# Stable, chronic in-vivo recordings from a fully wireless subdural-contained 65,536-electrode brain-computer interface device

**DOI:** 10.1101/2024.05.17.594333

**Authors:** Taesung Jung, Nanyu Zeng, Jason D. Fabbri, Guy Eichler, Zhe Li, Erfan Zabeh, Anup Das, Konstantin Willeke, Katie E. Wingel, Agrita Dubey, Rizwan Huq, Mohit Sharma, Yaoxing Hu, Girish Ramakrishnan, Kevin Tien, Paolo Mantovani, Abhinav Parihar, Heyu Yin, Denise Oswalt, Alexander Misdorp, Ilke Uguz, Tori Shinn, Gabrielle J. Rodriguez, Cate Nealley, Sophia Sanborn, Ian Gonzales, Michael Roukes, Jeffrey Knecht, Daniel Yoshor, Peter Canoll, Eleonora Spinazzi, Luca P. Carloni, Bijan Pesaran, Saumil Patel, Joshua Jacobs, Brett Youngerman, R. James Cotton, Andreas Tolias, Kenneth L. Shepard

**Author notes:** Contributed equally (Taesung Jung, Nanyu Zeng).

## Abstract

Minimally invasive, high-bandwidth brain-computer-interface (BCI) devices can revolutionize human applications. With orders-of-magnitude improvements in volumetric efficiency over other BCI technologies, we developed a 50-μm-thick, mechanically flexible micro-electrocorticography (μECoG) BCI, integrating a 256×256 array of electrodes, signal processing, data telemetry, and wireless powering on a single complementary metal-oxide-semiconductor (CMOS) substrate containing 65,536 recording channels, from which we can simultaneously record a selectable subset of up to 1024 channels at a given time. Fully implanted below the dura, our chip is wirelessly powered, communicating bi-directionally with an external relay station outside the body. We demonstrated chronic, reliable recordings for up to two weeks in pigs and up to two months in behaving non-human primates from somatosensory, motor, and visual cortices, decoding brain signals at high spatiotemporal resolution.

## INTRODUCTION

In electrophysiology, a fundamental trade-off exists between the invasiveness of the recording device and the spatiotemporal resolution and signal-to-noise ratio (SNR) characteristics of the acquired neural signals, ranging from scalp electrode discs to penetrating silicon probes. Non-invasive techniques such as electroencephalography (EEG) do not require surgery but can only capture limited spatiotemporal dynamics of brain activity ^1^. Recording from penetrating electrodes allows resolving extracellular action potentials from individual neurons ^2–4^; however, invasive microwires can cause tissue damage ^5–7^ and compromise long-term recording stability ^8–10^.

Electrocorticography (ECoG) is an intracranial approach that uses non-penetrating electrodes embedded in thin, flexible substrates that conform to the curvilinear surface of the brain. ECoG records population-level signals, averaged across local neurons. Importantly, because the electrodes sit on the cortical surface, ECoG minimizes brain tissue damage while being able to acquire higher signal-to-noise ratio (SNR), higher bandwidth, and more spatially localized signals compared to EEG ^11^. *In vivo* studies have shown that ECoG recordings taken from subdural arrays can remain stable for more than a year ^12,13^, demonstrating the technology’s potential usage in chronic applications. Furthermore, advances in microfabrication techniques continue to improve the spatial resolution of the ECoG electrode array. Current leading-edge micro-ECoG (μECoG) arrays implement more than a thousand recording sites with sub-millimeter electrode pitch on a single substrate ^14–17^. Such progress is encouraging since higher spatial resolution μECoG has been shown to improve the accuracy of sensory ^18^, motor ^19^, and speech decoding ^20^, more precisely map out epileptiform waveforms ^16^, and even record neural spiking activity ^21,22^.

Yet, current leading-edge high-resolution, multi-channel implantable electrodes, including but not limited to μECoG, remain separate from the electronics required for signal conditioning and data transmission. Traditionally, long percutaneous cables have been used to connect the implant to external rack-mount electronics ^23–27^, but the use of cables restricts the movement of subjects and increases the risk of infection and tissue damage ^28,29^. More recently, wireless electronics have been implemented to mitigate these concerns, by either using short percutaneous connection to a wearable headstage ^30–35^ or implementing a fully subcutaneously integrated system ^36–39^.

Current multi-channel, high-bandwidth wireless electronics, however, rely on the assembly of discrete, often commercial off-the-shelf components that lead to bulky form factors ^30–39^, which complicate surgical placement, removal, and revision ^40,41^. Combining discrete components also results in suboptimal electrical performance compared to what is achievable with application-specific integrated circuits (ASICs) and is hindered by limited interconnect density. This often becomes the bottleneck constraint on the number of recording channels and the scalability of these devices.

To implement the densest neural implants, electrodes and electronics need to be merged onto a single substrate. This has been the key to the success of Neuropixels ^23,24^, which feature penetrating neural electrodes capable of simultaneously recording from several hundred channels, with each probe equipped with thousands of electrodes and a wired connection to the probe. Here, we significantly extend the scale and nature of this CMOS integration of electronics and electrodes with the development of a device that we have named the Bioelectronic Interface System to the Cortex (BISC). BISC monolithically integrates a 256×256 high-resolution μECoG electrode array with front-end analog electronics, an on-chip controller, wireless powering, a radio frequency transceiver, and antennas onto a single CMOS substrate ^42^. For recording, BISC includes front-end circuitry for signal amplification and filtering and a back-end analog-to-digital converter (ADC). BISC also supports stimulation by including programmable bipolar constant-current sources. Our entire device is a 12×12 mm chip whose total thickness is rendered to be less than 50 μm after die thinning and passivation, giving the device enough mechanical flexibility to follow the contour of the brain.

This extreme miniaturization of function allows the BISC chip to be inserted under the dura on the pial matter using relatively simple and efficient surgical procedures, including the ability to replace chips at the same recording location after months, if necessary. Wireless powering and bi-directional communication are provided by a wearable device positioned directly outside the skin over the implant site. We call this device a “relay station” (**Fig. 1A**) because it wirelessly powers and communicates with the BISC implant while itself capable of being an 802.11n WiFi device. Recording fidelity and long-term efficacy of our device are demonstrated through a series of *in vivo* experiments on porcine and non-human primate (NHP) subjects. We implanted our device over multiple anatomical areas of the cortex and demonstrated high-quality chronic recording and accurate decoding of somatosensory, motor, and visual information. For example, from macaque visual cortex recording, our device captured complex spatiotemporal patterns of stimuli-induced traveling waves with spatial features on the scale of a few hundred microns and decoded stimulus orientation at a rate of 45 bits/sec. This minimally invasive, high-bandwidth chip, with its packaging-free, monolithic CMOS design and orders-of-magnitude improvements in volumetric efficiency and channel count compared to existing approaches, offers a scalable, cost-effective solution poised to revolutionize BCIs.

**Fig. 1.**
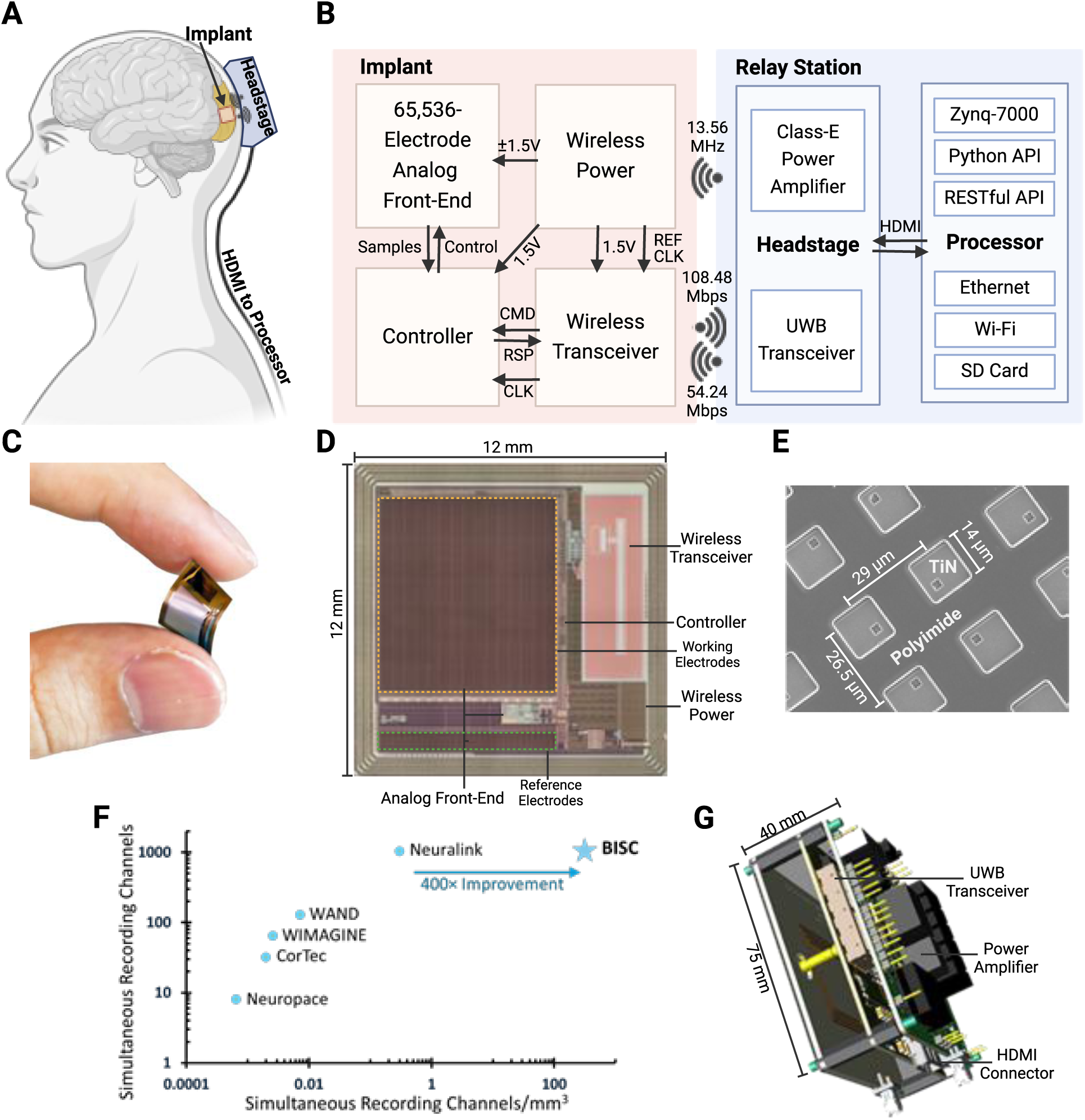
The BISC implant and relay station. (**A**) Concept diagram of the BISC implant and relay station. Relay station’s headstage provides wireless power and bi-directional communication from outside the body. The headstage module connects to the processor module (not shown) via HDMI cable. (**B**) Schematic of the overall system. The implant has four main modules: analog front-end (AFE) for recording and stimulation; wireless power for power transfer through an inductive link; wireless transceiver for bi-directional data telemetry; and controller for AFE configuration and data packetization. The headstage module receives a clock from the processor module; data moves between the headstage and processor modules with the processor module controlled by a computer through wired or wireless by means of Python/RESTful APIs. (**C**) Mechanical flexibility of the BISC implant. (**D**) Layout of the BISC implant. (**E**) Scanning electron microscopy (SEM) image of the titanium nitride (TiN) electrodes. (**F**) Comparison of our work with other competing wireless BCI devices. (**G**) 3D diagram of the headstage.

## RESULTS

### BISC implant form factor

A key metric for implantable BCI devices is “volumetric efficiency,” a measure of how much function can be achieved per amount of tissue displaced by the implant. Key to the volumetric efficiency of the BISC system is the chip, which constitutes the entire implanted system. This chip integrates a 256×256 microelectrode array (MEA) with a 26.5×29 μm pitch that results in a total array area of 6.8×7.4 mm (**Fig. 1C-E**). Each electrode is 14×14 μm in area and composed of titanium nitride (TiN), fabricated as part of additional microfabrication steps that follow the standard semiconductor foundry process (see **Methods** and **Figs. S3-S5**). As part of this “post-processing” which follows chip fabrication in a commercial CMOS foundry, the silicon substrate is also thinned to less than 25 μm, which, combined with the back-end metal stack, results in an overall thickness of approximately 40 μm (**Fig. 1C**, **Fig. S1B,D**). Polyimide is used to encapsulate the front-side, and parylene the back-side, resulting in a total device thickness of less than 50 μm. The final device has a bending stiffness of approximately 130 µN·m and, if crystalline defects are appropriately controlled during thinning, can be bent to a radius of curvature of about 1 mm for a strain of 1% in the silicon layer without fracture.

A fully processed BISC device has a total volume of 7.2 mm^3^. Its MEA provides a total of 65,536 recording channels, from which up to 1024 channels can be recorded simultaneously. We compare the form factor of our work with other state-of-the-art wireless brain-computer interfaces (BCIs). In this case, we define the volumetric efficiency as the number of simultaneous recording channels per unit implant volume (**Fig. 1F**, **Table S2**). The unique fully-integrated architecture of BISC allows it to achieve better than 400× improved volumetric efficiency over the closest competitor.

### BISC implant ASIC design

In the absence of any on-chip compression during recording, the number of simultaneous recording channels and the sampling rate are constrained by the uplink data bandwidth budget of 108 Mbps. To utilize the coverage and density of all the 65,536 available recording channels while preserving the ability to capture the full temporal dynamics of neural activity, we designed our device to digitize and transmit from a spatially programmable subset of either 256 channels at 33.9 kS/s or 1024 channels at 8.475 kS/s.

Every group of 2×2 neighboring electrodes in the array share the same recording pixel circuitry, resulting in 16,384 total pixels. When recording from 256 electrodes, the active subset can be programmed to any rectangle of 16×16 pixels, provided that their horizontal and vertical addresses are uniformly spaced. In this mode, only one of the four electrodes from each pixel is recorded. When recording from 1024 channels, the same 16×16 pixel addressing constraint applies, but now recording is taken from all four electrodes in each pixel (see **Methods** and **Supplementary Discussion S1**).

Each pixel performs signal amplification, chopping, and anti-aliasing filtering (AAF) and contains the associated digital logic to control these functions while occupying an area of only 53×58 μm (**Fig. S17**). This constrained area prevents the implementation of the capacitor-based filtering ^43^ or servo-loop-feedback-based or high dynamic range direct-quantization-based circuit topologies ^44^ generally used to prevent saturation in the presence of electrochemical DC offsets (EDO). We instead rely on the use of TiN electrodes, known to form non-Faradaic, capacitive interfaces ^45^ to reduce EDO. Electrochemical impedance spectroscopy (EIS) characterization confirms these properties for the BISC electrodes across the frequency range from 0.1 Hz to 1 MHz with an impedance magnitude of 205 kΩ at 1kHz, which is equivalent to an electrode capacitance of approximately 0.77 nF (**Extended Data Fig. 1A**, **Fig. S4**).

Instead of a traditional amplifier, the BISC pixel implements an integrator to provide area-efficient AAF from boxcar sampling principles ^46^ with an effective cut-off frequency at 15.1 kHz **(Extended Data Fig. 1B**, **Figs. S17** and **S26**). The use of an integrator also helps to mitigate open-loop gain variation across pixels (**Extended Data Fig. 1C**). All active pixels are time-multiplexed to share a single back-end programmable gain amplifier (PGA) through which a common gain is configured for all active pixels (**Extended Data Fig. 1B**, **Figs. S17, S18** and **S26**). For high-pass filtering, all pixels implement a variable pseudo-resistor that is globally configured across all active pixels (**Extended Data Fig. 1D**, **Fig. S26**). Input-referred noise integrated from 10 Hz to 4 kHz was measured to be 7.68 μV_RMS_ and 16.51 μV_RMS_ when recording from 256 and 1024 electrodes, respectively (**Extended Data Fig. 1E** and **F** and **Supplementary Discussion S3**).

The PGA output is digitized with a pair of interleaved 10-bit successive-approximation-register (SAR) analog-to-digital converters (ADCs) running at 8.68 MS/s. Ten samples of digitized data are grouped into a 125-bit packet by the on-chip controller (see **Supplementary Discussion S2**). The non-data bits in the packet are used for synchronization and error correction coding (**Fig. S23**). The transceiver has uplink and downlink data rates of 108 Mbps and 54 Mbps, respectively, and supports time-division duplexing with a single on-chip antenna by utilizing a transmit/receive (T/R) switch (**Supplementary Discussion S1** and **Fig. S20**).

Wireless power transfer (WPT) utilizes near-field inductive coupling at 13.56 MHz (**Supplementary Discussion S1** and **Figs. S21** and **S22**). This choice of frequency keeps the specific absorption rate below 2 W/kg ^47,48^ (**Fig. S22**). The system clock of the implant is derived from the WPT carrier, eliminating the need for a bulky crystal oscillator. BISC has a total power consumption of less than 64 mW, satisfying the thermal budget guideline of 0.5 mW/mm^2^ for neural implants ^49,50^ (**Fig. S22**).

### Relay station design

The relay station serves both to wirelessly power the implant and to transfer data between BISC and a computer base station, which can ultimately take the form of a smartphone. It is a two-part system that consists of a headstage and a processor module (**Fig. 1B**; see **Methods**), all designed from commercial off-the-shelf components.

The headstage has a wearable form factor (75×75×45 mm, 151 g) with a printed circuit board (PCB) stack-up that includes a powering coil and ultra-wideband (UWB) antenna (**Fig. 1G**, **Fig. S2**). For wireless powering, we use a tunable power amplifier to drive a custom spiral coil at 13.56 MHz that inductively couples to the receiving coil on the implant. For data telemetry, we use an impulse-radio ultra-wideband (IR-UWB) transceiver centered at 4 GHz with on-off-keying (OOK) modulation, where a custom dipole antenna on the headstage communicates with a monopole antenna on the implant.

The processor module controls the headstage through a standard high-definition-multimedia-interface (HDMI) cable. The processor module powers and configures PCB components on the headstage, sends queries and commands to the implant and receives responses and recorded data from the implant. Recorded data are either saved in the processor module’s non-volatile memory or re-directed to the computer base station over wired or wireless Ethernet. The processor module is based on a Xilinx Zynq-7000 system-on-chip (SoC) that includes a processing system (PS) unit and a programmable logic (PL) unit (**Fig. S24**). The PS integrates a dual-core ARM Cortex-A9 processor, runs the Linux operating system, and interfaces with a secure digital (SD) card used as the main non-volatile memory. The PL integrates specialized hardware to handle the bitstream of recorded neural data and store it in the ARM processor’s main memory. In addition, the specialized hardware on the PL embeds a high-level application programming interface (API) to generate and deliver pre-configured sequences of time-sensitive commands to the implanted device. We designed the API to be reconfigurable, leveraging its implementation on the PL, allowing it to be updated at any time. We implemented a Python software framework that uses the API and runs under the Linux operating system using PYNQ to control the specialized hardware in the PL. To support an interactive user environment, we designed a graphical user interface (GUI) that runs on the computer base station and interacts with the Python framework on the relay station via a RESTful API (**Fig. S25**). The GUI interacts with the relay station through the high-level API and displays neural data from the implant in real time.

Our bi-directional communication protocol uses a custom 125-bit packet tailored to accommodate the sampling rate and ADC resolution of BISC (**Fig. S23**, **Table S1**). The use of a custom protocol between BISC and the relay station results in more energy-efficient operation of this link when compared to the use of standard protocols such as Bluetooth or 802.11. Moreover, radio transmission between the BISC implant and the relay station only needs to be over a few centimeters, which allows our device to be a high-bit-rate radio transceiver while consuming less than 64 mW of total power (less than 13 mW peak power in the transceiver itself).

### Surgical approaches for implant

The thin, mechanically flexible form factor of the BISC implant enables relatively simple surgical implantation when compared to any device using penetrating intracortical electrodes or percutaneous connections. Device sterilization is performed with ethylene oxide (EtO; see **Methods**). In a technique common to both the pig and non-human primate (NHP) models, a standard craniotomy on the scale of 25 mm by 23 mm is made adjacent to the implantation site. The dura was then carefully elevated using sutures or forceps and linearly cut to provide a clear path for implantation. A commercial strip electrode is used as the insertion shuttle (see **Methods**, **Fig. S7**). The BISC chip was placed on top of the guide and then inserted under the dura, ensuring it rested directly on the pial surface. Importantly, the dural incision was made adjacent to the implantation site to avoid cuts or sutures directly over it. After securing the BISC chip in place, the dura was sutured, the skull was repositioned, and the incision was closed.

Furthermore, the procedure was designed to facilitate easy upgrading or replacement of the BISC chip. After several months of implantation, we reopened the skull in a NHP, made a new dural incision over the initial dural incision, and replaced the existing BISC chip in less than 10 minutes. This rapid, safe, and effective upgrade process is a significant advantage over other invasive BCI methods, where device replacement is more complex and time-consuming.

### Somatosensory evoked potential in porcine model

*In vivo* studies reported in this paper have focused on the recording capabilities of BISC; subsequent studies will focus on stimulation. Our device was first subchronically validated by implantation over somatosensory cortex in a porcine model. The pig brain’s size and gyrencephalic surface more closely resemble the human neocortex than those of commonly used small laboratory animals ^51^. Two weeks after the implant, we recorded somatosensory evoked potentials (SSEP) from the anesthetized animal in response to peripheral stimulation (see **Methods**). Five weeks after the implant, the whole brain was extracted postmortem for histological examination (**Fig. S9**).

For the SSEP recording session, percutaneous electrical stimulation was delivered to the median nerve and to four distinct locations of the snout (**Fig. 2A**) which is known to have a large somatotopic representation in the pig brain ^52^. The recording was taken from 16×16 channels, configured in the sparsest mode that covers the whole array. Data from these channels were low-pass filtered (300 Hz, eight-order Butterworth, zero-phase), subtracted by their baselines, and then down-sampled to 2.11 kS/s (see **Methods**). Trial-averaged channel responses (**Fig. 2B** and **C**, **Fig. S8**) show wave complexes whose polarization and depolarization peak timings are similar to those found in previous porcine SSEP studies ^53,54^. The spatial arrangement of the normalized peak response (**Fig. 2D**, **Video S1**) is also consistent with the previously reported somatotopy ^52,55,56^.

**Fig. 2.**
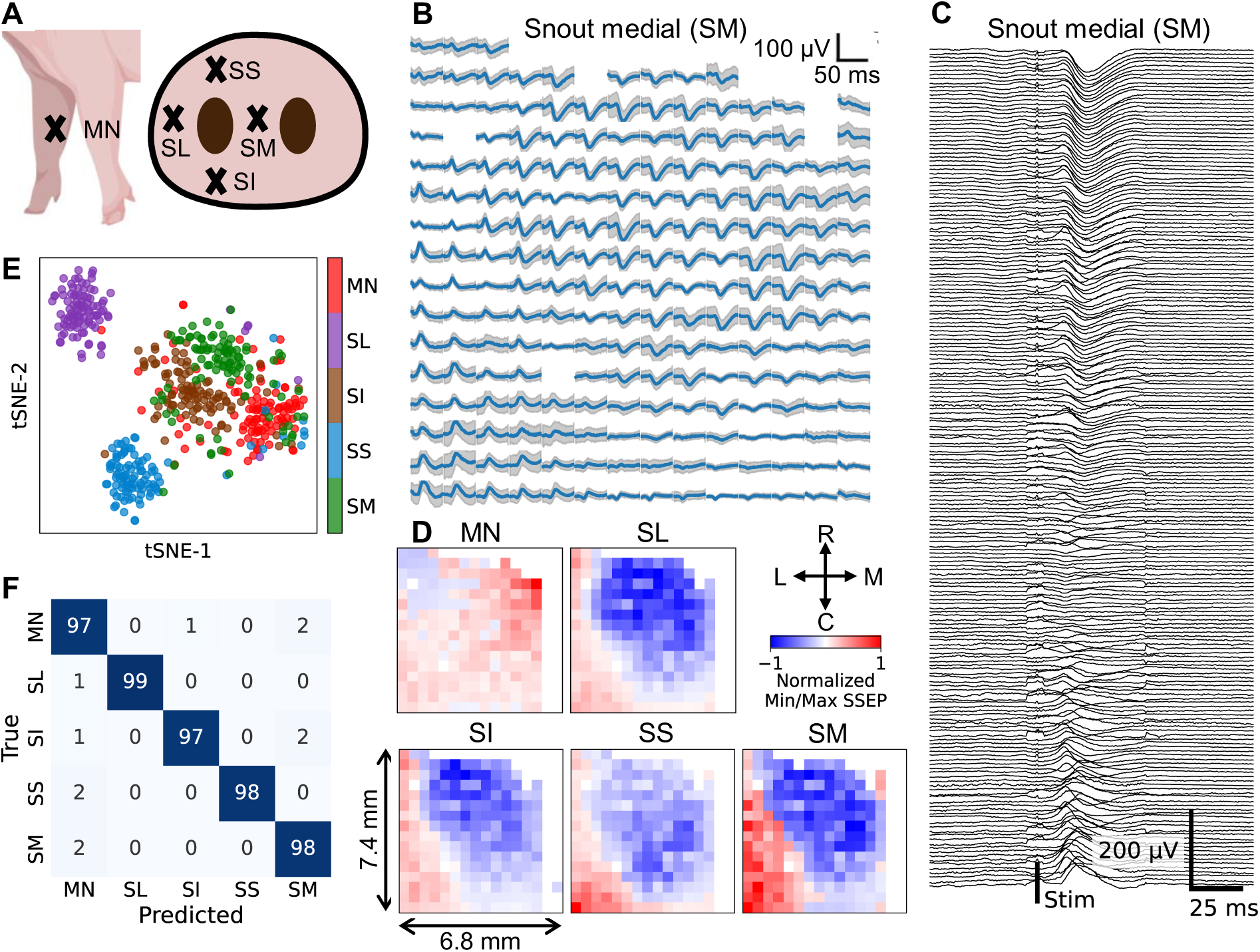
BISC recordings over somatosensory cortex from a porcine model. (**A**) Location of stimulation (MN: median nerve, SS: snout superior, SL: snout lateral, SI: snout inferior, SM: snout medial). (**B**) Example of stimulation evoked somatosensory evoked potential (SSEP), trial averaged (n = 100). Shaded area indicates SD. (**C**) Representation of (B) aligned on a shared time axis. (**D**) Spatial map of normalized SSEP extrema. Positive value indicates that peak SSEP magnitude was greater than the trough magnitude. Negative value indicates that trough SSEP magnitude was greater than the peak magnitude. Cross indicates orientation of the chip in (B) and (D) (R: rostral, C: caudal, L: lateral: M: medial). (**E**) State-space representation of the SSEPs using t-distributed stochastic neighbor embedding (t-SNE, n = 500). (**F**) Prediction of stimulation location using a linear discriminant classifier (n = 500).

Decoding peripheral stimulation site from SSEP was assessed through two approaches: visualizing the data projected into two-dimensional space and constructing a classification model (see **Methods**). For both analyses, we first used principal component analysis to project the z-scored spatiotemporal waveforms into a lower-dimensional space, retaining components that explained 80% of the variance (see **Methods**). These extracted components were projected onto a two-dimensional space using t-distribution stochastic neighbor embedding (t-SNE) ^57^. This projection (**Fig. 2E**) results in clearly separable clusters, indicating that the measured SSEP can be used to effectively distinguish between peripheral stimulation sites. Decoding performance was quantified using a linear discriminant model with 10-fold cross-validation that resulted in an overall accuracy of 97.8 ± 1.7 % (mean ± SD, n = 500) (**Fig. 2**F).

Coronal sections were collected from the post-mortem extracted brain for histological analysis using three markers: hematoxylin and eosin (H&E), NeuN, and Iba1 (**Fig. S9**). Sections taken directly under BISC showed no significant pathology by H&E or NeuN. Iba1, however, revealed a mild microgliosis extending from the superficial cortex to the subcortical white matter. Near the perimeter of our device, there was a small lesion on the surface of the cortex, consistent with a mechanical injury incurred during electrode placement. No pathological changes were seen in sections taken as control samples from the occipital cortex.

### Motor cortex recording in NHP

Prior to the chronic NHP study, BISC was first validated through acute recording using a glass artificial skull mounted in a permanent craniotomy over the motor cortex region of a behaving NHP subject. In this setup, the glass module is hermetically secured to a base ring affixed to the subject’s skull using mechanical screws, eliminating the need for additional surgery for device implantation. The subject was seated in a primate chair, facing the experimenter who manually held a wand at the subject’s full reach distance. The subject was trained to asynchronously reach, grab the wand, and then retract its arm without the use of an explicit cue to prompt its movement, with the arm location triangulated from multiple cameras (**Fig. 3A**; see **Methods**).

**Fig. 3.**
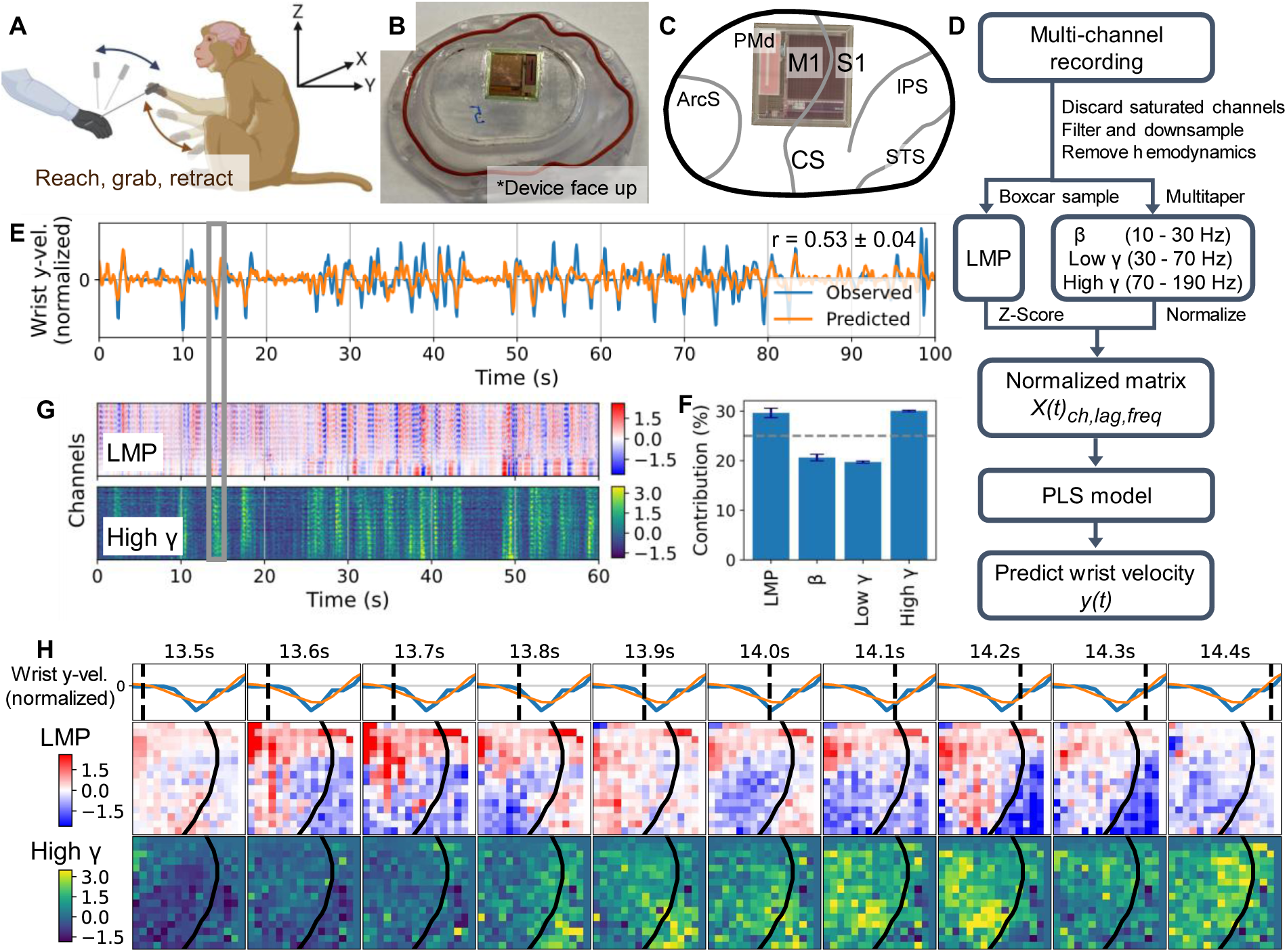
BISC recordings over motor cortex from a behaving non-human primate. (**A**) Behavioral task depiction. The subject was trained to asynchronously reach and grab a “wand” held by the experimenter without an explicit cue. (**B**) Device placement on the artificial skull. (**C**) Registration of the device with respect to the brain surface anatomy. (**D**) Motor feature decoding pipeline. A linear partial least squares (PLS) regression model uses spatial-temporal-spectral input 𝑋(𝑡) to predict the subject’s motor feature, 𝑦(𝑡). (**E**) Representative example of continuous decoding of normalized wrist velocity in y-direction (front-back). (**F**) Spectral contributions to the decoder from each frequency band. Error bars indicate SE, and dashed line indicates chance level. (**G**) Representative example of spatiotemporal dynamics of local motor potential (LMP) and high γ band, both z-scored. Time axes in (E) and (G) are shared. (**H**) Representative example of frame-by-frame spatiotemporal progression of LMP and high γ band, both z-scored. Plotted time window is indicated by the vertical gray box over (E) and (G). Dashed lines indicate time instants of each frame, and solid curves over the frames represent the central sulcus (CS) in (C).

For this experiment, the BISC device was attached directly to the artificial skull (**Fig. 3B**) and positioned over the central sulcus (CS), with most electrodes over the primary motor cortex (M1) but a significant portion over the primary somatosensory cortex (S1) (**Fig. 3C**).

While the subject was performing the behavioral task, BISC recordings were taken from 16×16 channels configured in the sparsest mode covering the whole array. The fidelity of measured multi-channel local field potentials (LFPs) was assessed by building a continuous velocity decoder using a linear model (**Fig. 3D-F**; see **Method**s) and visualizing the spatiotemporal dynamics of selected frequency bands (**Fig. 3G-H**).

The decoder was built by first identifying channels (n = 180 out of 256) that consistently remained non-saturated throughout the experiment (see **Methods**). Data from these channels were band-pass filtered (0.3 to 300 Hz) and down-sampled to 2.11 kS/s. Recordings contained a strong hemodynamic rhythm around 3 Hz, which we associated with the heart rate (180 beats/minute). This rhythm was removed by subtracting the corresponding time series components computed through space-time singular value decomposition (**Fig. S10**) ^58^. The pre-processed recordings were further grouped into four frequency bands: local motor potential (LMP), β (10 to 30 Hz), low γ (30 to 70 Hz), and high γ (70 to 190 Hz). Low-frequency LMP was extracted by applying boxcar averaging with a 50 ms window and then z-scoring ^59^. The other three bands were extracted by applying multitaper estimation with a 200-ms window and 10-Hz half-bandwidth and then normalizing ^60^. Finally, the time history of features from (t −0.47) s to (t + 0.47) s in 52 ms time steps were used for decoding, resulting in 13,680-dimensional vector 𝑋(𝑡) (180 channels, 19 time lags, four frequency bands) as model input for decoding motor feature 𝑦(𝑡).

We decoded the arm velocity using partial least squares (PLS) regression because it is effective in handling data whose predictor is highly correlated and has a large dimension compared to the number of observations ^61^. The optimal number of PLS components was determined by calculating the minimum predictive error sum of squares across 5-fold cross-validation (**Fig. S10**). When decoded against the normalized y-dimension (front-back) wrist velocity, feature prediction resulted in Pearson’s correlation coefficient of 0.53 ± 0.04 (mean ± SE), illustrated by an example time segment shown in **Fig. 3E**. Spectral contributions to the decoder from each frequency band were found by computing the sum of the relative weight of coefficients associated with each band (**Fig. 3F**).

The two bands with the highest contributions – LMP and high γ band – are further visualized by plotting the multi-channel power over time (**Fig. 3G**) which shows fluctuations synchronized to the subject’s motion. Spatiotemporal progression of these bands in a selected one-second-long time window (**Fig. 3H**) shows spatially localized activity in both M1 and S1, with LMP displaying patterns that resemble phase reversal across the CS boundary. A more detailed version of this plot with a finer time resolution and an extended time range can be found in **Video S2**.

### Visual cortex recording in NHP

The stability of chronic neural signals from BISC was validated through long-term (up to 64 days) studies from the visual cortex of an adult macaque monkey. The chip was placed near the border of V1 and V2, partially covering V4 as well **(Fig. 4A**). Over the span of the study, we conducted multiple experiments that involved three different visual stimulus paradigms: gratings, random dots, and natural images.

**Fig. 4.**
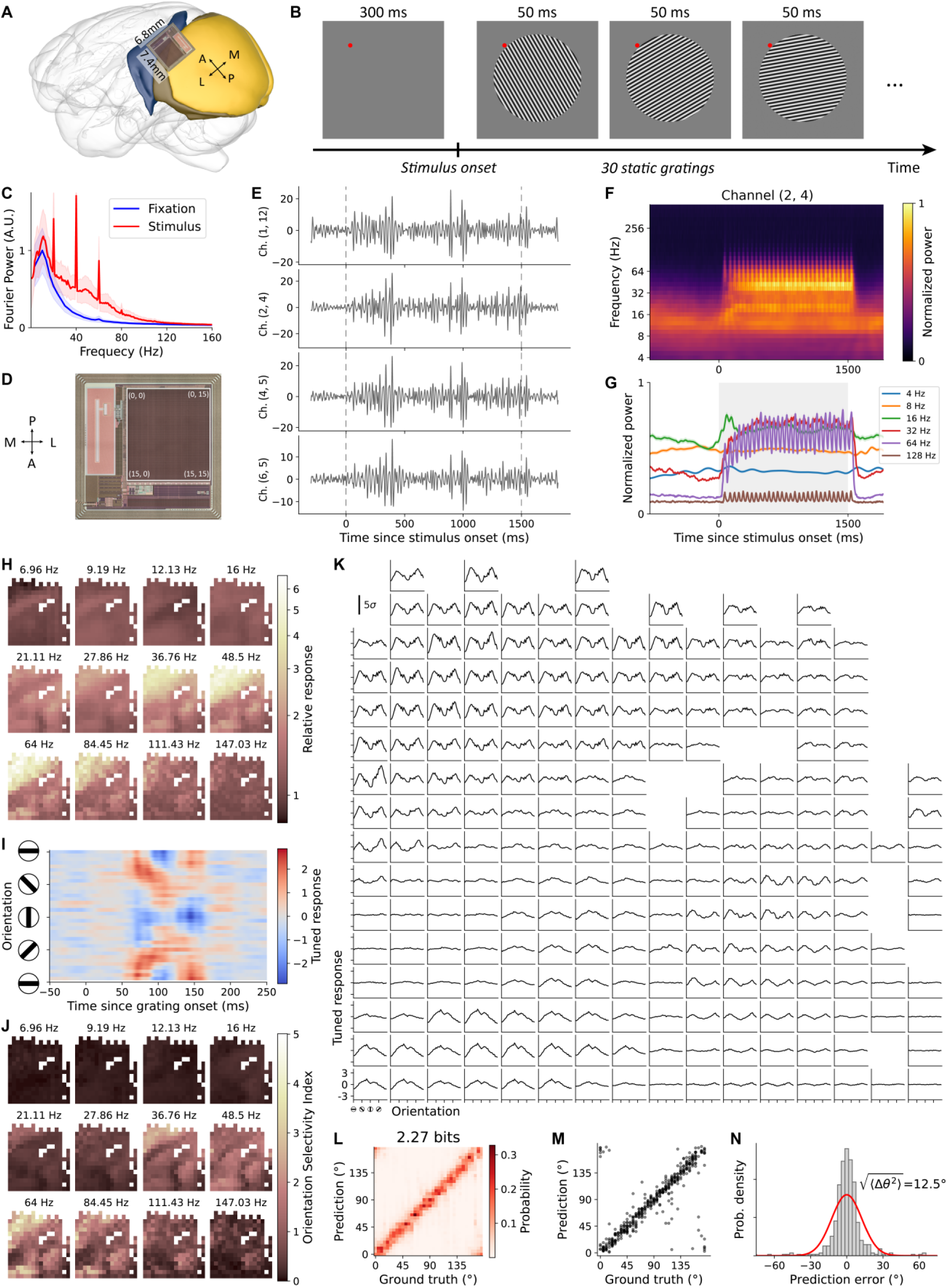
BISC recordings over visual cortex with grating stimuli. (**A**) Chip position on the cortex. The chip was placed on the border between V1 and V4 areas. Note that a mirrored image is used here as the chip is facing down. (**B**) A trial in the grating session. The monkey is required to fixate for at least 300 ms before visual stimuli are presented on the monitor. 30 static gratings with random orientations are displayed in one trial, each lasting 50 ms. (**C**) Spectrum comparison between fixation period and stimulus period. Fourier transform is applied to responses in the time window between −300 and 0 ms (blue) and the time window between 500 and 1500 ms (red) relative to stimulus onset. Solid lines are average over channels and trials; shaded areas mark the standard deviation of trial-average over channels. (**D**) Channel coordinates in full chip recording mode. We simultaneously record 256 channels at 33.9 kS/s from the full chip. (**E**) Filtered responses of example channels. For four channels that are strongly responsive to grating stimuli, responses after removing Fourier components below 20 Hz and above 90 Hz for one trial are shown. Vertical lines indicate the onset of stimulation. (**F**) Trial-averaged spectrogram of one channel. We applied wavelet transformation on the recorded signal and obtained time-varying power for different frequency bands. (**G**) Temporal profiles of selected frequency bands. Six rows from (F) are plotted as solid lines. Shaded area is the standard error of mean across trials. Gray background marks the period when grating stimuli are presented. (**H**) Responsiveness maps of different bands. Responsiveness for each channel in given frequency bands is defined as the signal power during the time window from 500 to 1500 ms divided by that during −300 to 0 ms. (**I**) Orientation tuning of an example channel. We computed the grating-triggered-average response conditioned for each orientation. The simple average contains a strong orientation-independent component that reflects the switching of grating every 50 ms (see Supplementary). The heatmap shows orientation tuned components after the removal of orientation-independent component. (**J**) Orientation selectivity maps of different bands. Orientation selectivity index is defined based on the tuned response during the time window from 88 to 112 ms after grating onset (see Methods). (**K**) Orientation tuning curves of all channels from 64-Hz band. Tuned response averaged over the time window from 88 to 112 ms as a function of grating orientation is shown for all channels. The unit of each channel’s response is the standard deviation 𝜎 across trials computed during fixation period. (**L**) Orientation decoder performance. We trained a decoder that takes raw responses from all channels in the time window from 0 to 200 ms after a grating onset, and predicts the grating orientation. The decoder outputs a distribution over orientations. The average decoder output is shown for each orientation on a hold-out testing set of trials. Mutual information between prediction and response is computed after discretizing orientation into 36 bins. (**M**) Point estimation on testing set. Circular mean of decoder outputs is computed as point estimations. (**N**) Histogram of prediction error. The difference between point estimation and ground truth is gathered for all gratings in testing trials.

In the grating sessions, the monkey fixated for at least 300 ms to initiate a trial in which 30 static gratings of random orientations were presented consecutively (**Fig. 4B**). Each grating lasted 50 ms. BISC recordings were taken from 16×16 channels configured in the sparsest mode covering the whole array. Approximately one-third of them, primarily those overlying V1, showed increased power for Fourier components between 20 Hz and 90 Hz (**Fig. 4C**). The peaks at 20 Hz and harmonics in the power spectrum response reflect the grating changing every 50 ms. We selected four example channels labeled by their chip coordinates (**Fig. 4D**) to show their band-pass filtered responses in one trial (**Fig. 4E**). It is evident that activity in these channels is elicited by the onset of the visual stimulus and returns to baseline after the stimulus. We further applied Morlet wavelet transformation (see **Methods**) to acquire the spectrogram of each channel. The averaged spectrogram of one channel after aligning at the stimulus onset is shown (**Fig. 4F**), along with six individual bands with central frequencies evenly spaced on a logarithmic scale (**Fig. 4G**). We observed different responses in each band, defined as the ratio of the mean response in the time window from 500 to 1500 ms relative to the stimulus onset over the mean response in the time window from −300 to 0 ms. The resulting response maps are shown for multiple bands (**Fig. 4H**).

For each frequency band, we computed the grating-triggered-average response by aligning response segments according to the grating onset and obtained the average band-passed scaled response conditioned on each grating orientation (see **Methods**). The result shows a strong untuned component corresponding to the grating switching (**Fig. S11**). After removing it, the residual tuned components show that channel responses during the time window from 50 to 200 ms after grating onsets are tuned to the grating orientation. A typical channel is shown in **Fig. 4I**. We take the average response between 88 and 112 ms after grating onset to measure a tuning curve for each channel and define the orientation selectivity index as the difference between the maximum and the minimum value. The orientation selectivity index maps of multiple frequency bands are shown in **Fig. 4J**. Tuning curves for the 64-Hz band are shown in **Fig. 4K**. The channels with pronounced tuning and greater orientation selectivity indices correspond to those overlying V1. As expected, we observe strong orientation tuning in the gamma band.

We then built an orientation decoder from these responses which takes raw responses (with no filtering) of all channels in the time window from 0 to 200 ms after grating onset and returns a distribution of grating orientation. We trained the decoder as a classifier in which orientation is discretized into 36 bins. The decoder is a multi-layer convolutional neural network (CNN) that uses one-dimensional (1D) convolution along the temporal dimension. The performance of a hold-out set (containing trials not used during training) is shown in **Fig. 4L**. The mutual information 𝐼(𝜃; 𝒓) = 𝐻(𝜃) − 𝐻(𝜃|𝒓) between predicted orientation 𝜃 and BISC responses 𝒓 is computed as approximately 2.27 bits, which constitutes a data rate of 45 bits/sec at a 50-ms frame rate (see **Methods**; **Figs. S14** and **S15**). We found that the decoder prediction approximates the ground truth values (**Fig. 4M**), with a root mean squared error of 9.1° (**Fig. 4N**).

We mapped spatial receptive fields (RFs) using a random dot experiment, where single 0.51° dots appeared on a uniform gray background, changing location and color (black/white) every 50 ms (**Fig. 5A**). Dots were presented within a 6°×6° rectangular field, centered 1.5° right and 3° below fixation. The monkey maintained fixation for 1500 ms to receive a juice reward. After removing Fourier components outside the 20-to-90-Hz band, the band-passed responses for all non-saturated channels are shown for one trial (**Fig. 5B**).

**Fig. 5.**
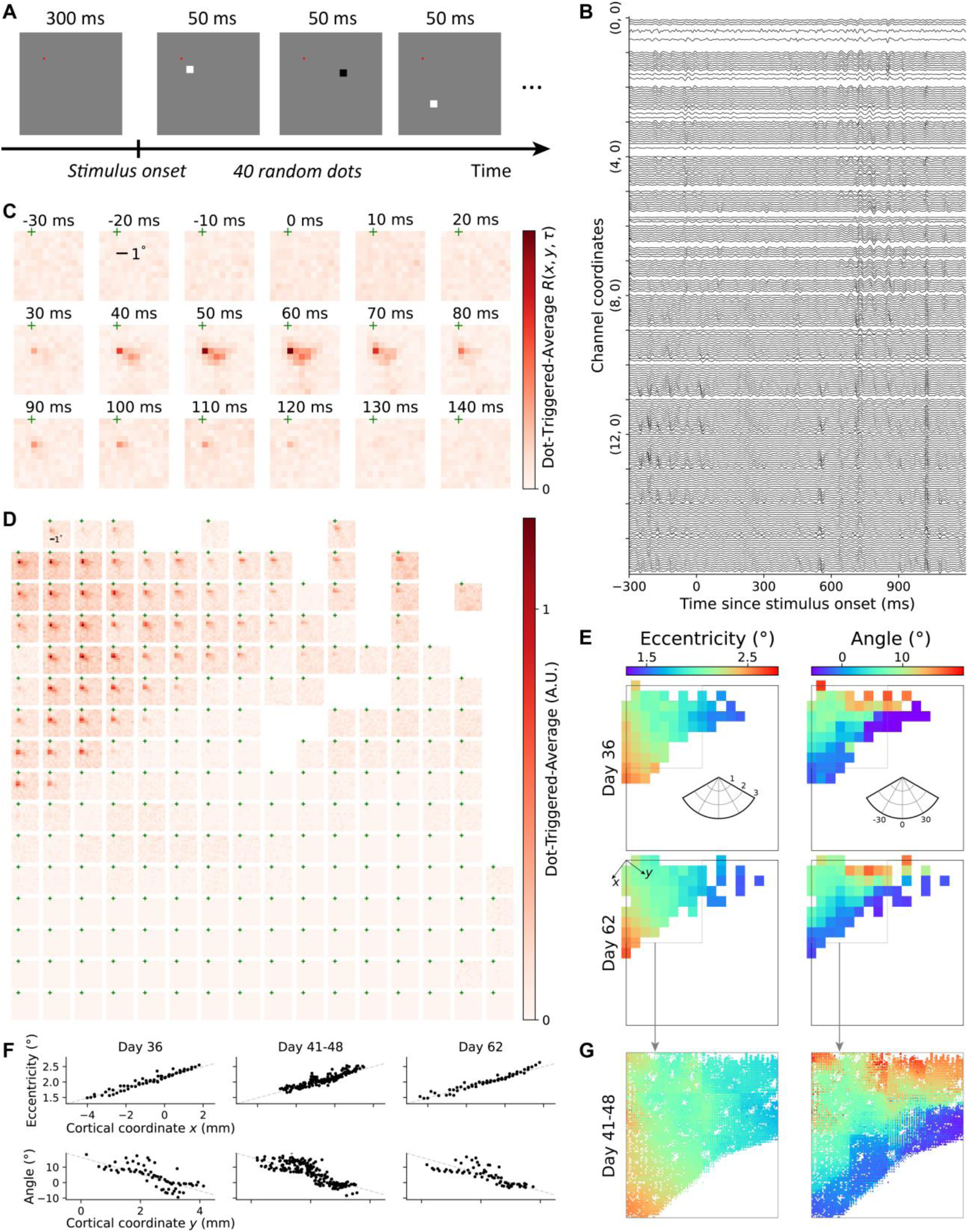
BISC recordings over visual cortex with dot stimuli. (**A**) A trial in the dot mapping session. 40 random dots of either black or white color is shown in the area [-1.5°, 4.5°]×[0°, 6°] (horizontal×vertical), each lasting 50 ms. (**B**) Filtered response of all non-saturated channels from a full chip recording session. We show their responses after removing Fourier components below 20 Hz and above 90 Hz for one trial. (**C**) Spatial-temporal receptive field (RF) of one example channel. We filtered the response by a Morlet wavelet of which the central frequency is 64 Hz, and computed the dot-triggered-average of signal power. Times are relative to dot onset and cross symbol represents the fixation location. (**D**) Spatial RF of all non-saturated channels. Spatial RF is computed by taking the temporal average of dot-triggered-average responses. (**E**) Chronic stability of retinotopic maps. A 2D Gaussian fit is computed for the spatial RF of each channel separately, and we take the fitted center as RF center. Channels with good Gaussian fit (fraction of unexplained variance smaller than 0.8) are shown for recordings from day 36 and 62 after surgery. (**F**) RF locations projected on cortical axes at Day 36, Days 41-48, and Day 62 after surgery. We denoted the direction along which RF eccentricity changes fastest on the chip as ‘x’, and the orthogonal direction as ‘y’. RF centers are plotted against the channel projections on ‘x’ and ‘y’ separately. Overall, the eccentricity changes roughly at the rate 0.17 visual degree per millimeter. Across days, the receptive field changed by less-than-0.05-degree in eccentricity and less-than-2.7-dgree in angle on average for all channels for both dense (Days 41-48) and sprase (Days 36 and 64), demonstrating the stability of BISC chronic recordings. (**G**) Zoom-in retinotopic map for the upper-left quarter of the chip. We performed 16 dense recording sessions over 8 days, with 1024 channels simultaneously recorded in each session.

We computed the RF by dot-triggered averaging on wavelet-transformed responses. Multiple wavelet central frequencies were tested (**Videos S3** to **S8**) and almost all of them show clean structure for channels in the V1 area. An example channel is shown in **Fig. 5C** for the 64-Hz band. This channel is responsive to a dot shown in a lower-right location approximately 40 to 80 ms after dot onset. By averaging the spatial-temporal receptive field over time, we can compute an estimate of the spatial RFs, which are shown with a shared color scale (**Fig. 5D**). While channels in V1 have compact RFs, the presented dot size is too small to invoke spatially structured response from channels outside of V1. We fit two-dimensional (2D) Gaussian functions to the computed RFs and plot the retinotopic map using the fitted Gaussian centers (**Fig. 5E**). We examined the chronic stability of these retinotopic maps by comparing these maps at Day 36, Days 41-48, and Day 62 after surgery. The receptive fields for these channels remained stable over this period with a less-than-0.05-degree change in eccentricity and less-than-2.7-degree change in angle on average for all channels from Day 36 to Day 62 for both dense (at Days 41-48) and sparse (at Days 36 and 62) recordings, demonstrating the stability of BISC chronic recordings (**Fig. 5F**).

To exploit the dense 65,536-channel structure of BISC, we densely recorded (minimum electrode pitches at 26.5 µm by 29 µm) from the top-left quadrant of the electrode array through repeated, contiguous recording blocks of 1024 channels. The combined results (**Fig. 5G**) are consistent with what we found in the sparse full-array recording sessions while providing a much higher resolution retinotopic map.

Next, we hypothesized that these spatially dense recordings capture fine-grained spatiotemporal dynamics of LFPs that cannot be resolved in the sparse configuration. Specifically, we hypothesized the existence of ‘traveling waves’—spatiotemporally coherent patterns of propagating neural oscillations—given recent findings showing that such waves can encode task-related spatial information^62^. We focused on identifying traveling waves in the gamma band (30-90 Hz), given the task-related oscillations in this range (**Fig. 4F-J**). To measure gamma-band traveling waves, we extracted the instantaneous phase of the gamma oscillation at each channel in the dense configuration. Then, we measured the topography of the gamma traveling waves, using circular statistics to extract the instantaneous direction and strength of the gamma phase gradient at each contact (see Methods). With this approach, we compared the strength of gamma traveling waves across dense recordings taken from 16 different locations of the array during dot viewing. The strongest traveling waves were recorded from the top-left corner of the BISC array, corresponding to posterior medial recordings over the operculum in V1 (see red outline in Fig. 6Ai). Our subsequent analysis (**Fig. 6Aii, B–E**) focuses on recordings from this section of the array.

**Figure 6.**
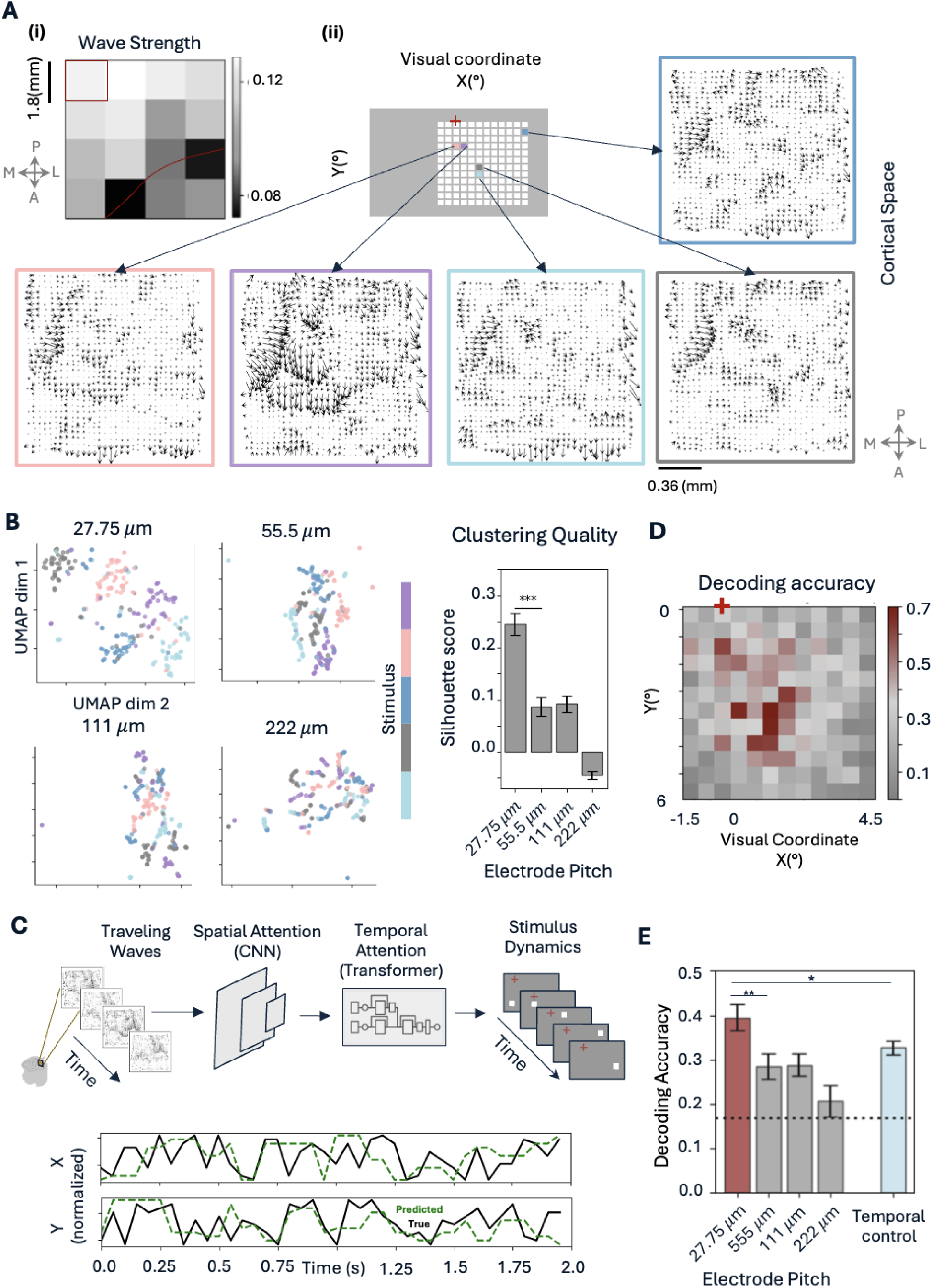
Spatially dense BISC recordings over visual cortex capture complex traveling wave patterns that encode the locations of viewed dot stimuli. (**A**) (**i**) The mean strength of traveling waves at each of 16 dense recordings taken from subsets of the BISC array during the dot mapping experiment. These 16 recordings are also presented in Fig. 5G; red line indicates the V1-V2 boundary identified from the retinotopic map. Recordings over V1 show higher wave-strength compared to those over V2, consistent with the enhanced functional role of V1 compared to V2 in processing simple visual features such as dot locations. (**ii**) Each dot stimulus location evokes a distinct pattern of traveling waves. Traveling wave topographies captured by 1024-channel, dense recordings taken from the top-left subset of the array shown in panel A(i) (red box), displayed for each of five example dot locations. Traveling waves are estimated from the gradient of the phase of oscillations in the gamma band (30-90 Hz) in the 40-90 ms interval relative to the dot onset, with this interval chosen as it evokes the maximum neural response (Fig. 5C). Note the spatially complex shape of the traveling waves, with even adjacent dots evoking spatially distinct patterns. (**B**) Separation of the traveling wave patterns between different dot locations visualized using Uniform Manifold Approximation and Projection (UMAP). Low-dimensional UMAP embeddings are shown for four different channel resolutions as defined by average electrode pitch. Colors indicate dot stimulus locations. Note that dot locations are clearly separated in UMAP space for 100% sampling. Separation decreases for spatial sampling at lower resolutions. Separability of dot locations is quantified using the silhouette score, with higher scores reflecting greater separability at higher spatial resolution (t-test, t(527) = 5.71, p <0.0001). (**C**) Decoding dot locations from the traveling waves at locations shown in (A) using a hybrid CNN-Transformer model. Predicted x- and y-coordinates closely match true dot locations, as shown in the accuracy trace plots. (**D**) A heatmap displays prediction accuracy for individual dot locations, illustrating decoding variability across the visual field. Dots presented closer to the center of the visual field (dark red) exhibit higher decoding accuracy, matching the receptive field maps. (**E**) Decoding accuracy as a function of the spatial sampling of the BISC channels, showing the importance of the BISC chip’s high-density recordings for recording relevant spatially precise features of traveling waves (t-test, t(285)=2.6, p<0.001). Temporally shuffled controls confirm the temporal dependencies of these high-resolution traveling waves, as temporal shuffling of the order between consecutive stimuli reduces the predication accuracy for individual dots (t-test, t(219)=2.0, p<0.05). Error bars indicate standard error of the mean across cross-validation folds.

First, we compared the spatial topographies of the traveling waves induced by dot stimuli (**Fig. 6Aii**). This analysis revealed distinct wave patterns on the cortex that differed according to the location of the presented dot in the visual field. Examining these signals more closely, we tested whether spatially dense BISC recordings revealed additional detailed stimulus-relevant information that was not evident at lower resolutions. For five example dot locations, using UMAP clustering analysis (see **Methods**), we found that spatially dense BISC recordings more effectively distinguished the traveling wave patterns corresponding to different viewed dot locations, compared measurements of these traveling waves at lower resolutions (**Fig. 6B**).

To expand this approach and test whether traveling waves encoded information about dot locations across the visual field, we built a neural network decoder to predict the horizontal and vertical coordinates of the dot viewed by the monkey based on the spatiotemporal pattern of BISC- measured traveling waves. This decoder uses a hybrid neural network architecture, including a convolutional neural network (CNN) that assesses the topography of traveling waves at each moment and a transformer network to extract long-range temporal correlations. The model was trained by using the spatiotemporal sequence of traveling waves from each trial to predict the current dot location viewed by the monkey (**Fig. 6C**; **Video S9**; see **Methods**). We used this network to assess the statistical robustness of dot-specific traveling waves over time and compared the accuracy in predicting the coordinates of the viewed dot as a function of recording resolution.

By feeding our model with the sequence of gamma band traveling waves observed in each trial, we could reliably decode the coordinates of individual presented dots, with highest accuracy for dots near the center of the visual field (**Fig. 6D**). Decoding accuracy is significantly higher when the traveling waves are computed from full-resolution BISC recordings, compared to lower resolutions (**Fig. 6E**, full resolution versus quarter resolution: t-test, t(285)=2.6, p<0.001).

To test the temporal dynamics of traveling waves, we next assessed whether the predictive model for the current dot stimulus depends not only on the current traveling wave but also on the past history of traveling waves and stimuli. We tested this hypothesis by asking the decoding model to predict the dot location for a new traveling wave pattern immediately after being presented with a shuffled sequence of dots and matching traveling wave patterns. This shuffling procedure resulted in significantly decreased decoding accuracy (**Fig. 6E**, t-test, t(219)=2.0, p<0.05), indicating that the traveling waves at each moment are not only dependent on the currently viewed dot but also on the recent histories of stimuli and cortical activity. Thus, traveling waves encode information about the current stimuli and past events^63,64^, which can be extracted by the CNN and transformer network.

Lastly, we studied the non-linear characteristics in the responses of BISC recording channels when presented with colored natural images (**Extended Data Fig. 2A**). In this paradigm, we showed the fixating monkey a large number of colored natural images from the ImageNet database ^65^. In each of the five recording sessions with sparse recording over the entire array of 16 × 16 channels, 10,000 – 12,000 unique natural images were presented in blocks of 15 images with each image displayed for 120 ms ^66,67^. The images (10° by 10° in size) were centered at a location 3° to the right and below the fixation spot. We then applied zero-phase component analysis (ZCA) whitening across all channels and band-pass filtered the signal between 30 and 90 Hz (with a tenth-order Butterworth, zero phase); this was followed by squaring and averaging of the signal from 40 ms to 160 ms after the image onset. Through these processing steps, we derived a scalar magnitude from the gamma band of each channel, known to reflect the spiking activity of the adjacent neuronal population ^68^, in response to each natural image that was presented. We then fit a deep neural network model (**Extended Data Fig. 2B**) to the pairs of natural images and gamma-band responses to learn a stimulus-response mapping, i.e. a digital twin of each recorded channel, such that the model can predict the activity of any channel when presented with an arbitrary stimulus. Then, we selected and fine-tuned an ImageNet pre-trained CNN model ^69^ as a shared feature space, with learned channel-specific weights, to predict the responses of individual channels ^70^. Next, we tested the accuracy of our digital twin model on a held-out set of 75 natural images. Each image was repeated 40 times and was never shown to the model during training.

As a measure of predictive performance (**Extended Data Fig. 2C**), we computed the correlation between the model prediction and the averaged neuronal response to repeated presentations of the same test image, which resulted in a mean of 0.69 ± 0.14 (mean ± SD) across all selected channels. This is comparable or better than the performance of similar predictive models of isolated single neurons in macaque V1 ^66,67^. Additionally, we computed the explainable variance for each channel, which measures the response reliability to natural images. This metric is defined as the ratio of response variance across repeated presentations of the same image over the response variance across all images. (**Extended Data Fig. 2C)**. We only selected channels with an explainable variance larger than 0.1. Across all channels, we obtained an average explainable variance of 0.24 ± 0.09 (mean ± SD), slightly below the values of 0.31 and 0.32 that were reported for isolated single neurons in macaque V1 and V4, respectively ^66,67^. The spatial distribution of the model performance (**Extended Data Fig. 2D**) and response reliability (**Extended Data Fig. 2E**) across the recording array indicated that channels closer to V1 exhibit higher model performance and response reliability compared to those in higher visual areas. We then visualize the feature selectivity of each channel by using our predictive model (digital twin, **Extended Data Fig. 2F**). Traditionally, parametric stimuli or hand-designed images are used to investigate the visual feature selectivity of single cells or populations of neurons. The stimuli that elicit the highest responses are then used to determine the tuning functions of the neurons in question.

More recently, digital twin models have been used to synthesize optimal stimuli by iteratively optimizing a starting image such that the response of a neuron or channel is maximized. These optimized images are referred to as maximally exciting images (MEIs) and can be thought of as non-linear receptive fields ^71^, shedding light on the underlying neural response functions. MEIs have recently been used to reveal visual tuning characteristics in mouse V1 ^71–74^, as well as monkey V1 ^72^, and V4 ^66,75,76^. Here, we show the MEIs for individual channels to demonstrate their visual feature tuning. Remarkably, the MEIs (**Extended Data Fig. 2G**) reveal a hierarchy of complexity in visual feature tuning from area V1 (top left) to V2 (center) to V4 (bottom right). Area V1 is characterized, as expected from MEIs of single isolated neurons V1 ^72^, with oriented Gabor filters. Areas V2 and V4 are dominated by more complex features with dominant color-opponency (also observed in ^75^). The digital twin model is thus able to capture detailed non-linear visual response characteristics of the adjacent neuronal population of each recorded channel.

## DISCUSSION

This project pioneers a new paradigm in BCI devices that can be implanted entirely subdurally. Like traditional µECoG arrays, it relies exclusively on recording and stimulation from the surface, albeit at a dramatically higher electrode density than any previous system. The advantage this device has over those relying on intracortical electrodes, particularly in the context of eventual human translation, lies in its surgical insertion; its removal and replacement are very straightforward, and little to no cortical damage results from using the BISC design. In the future, we expect to transition to less invasive surgical procedures in which only a small slit is made in the cranium and dura, and the BISC implant is “slid” under the skull.

Positioning BISC on the pial brain surface allows it to naturally conform to the contour of the brain and move with the brain, bringing significant advantages over devices with components attached to the skull which introduce relative motion between the device and the brain. An additional benefit of subdural μECoG recordings over intracortical recordings for clinical translation BCI is their stability over time which was replicated in BISC.

BISC was designed with a front-end recording bandwidth sufficient to capture the temporal dynamics of neural spiking activity. Our *in vivo* experiments did not demonstrate the existence of large-scale spiking activity from BISC surface recordings. Future iterations of BISC focused on only LFP recordings can reduce the front-end recording bandwidth and better utilize the available 108 Mbps data bandwidth to simultaneously record from a larger number of channels. The architecture of BISC in which full recording channels are present for all electrodes aids in this alternate implementation. The high-density recording capability of the BISC device shows that even for LFPs, when measured with small surface electrodes placed directly on the pial surface, these signals exhibit precise spatiotemporal patterns at a scale of tens of microns. These patterns convey information about both current and past stimuli, as demonstrated by decoder performance trained on gamma-band traveling waves in the primary visual cortex.

Prior work has measured traveling waves with more widely spaced electrodes in scales of centimeters^4^ or millimeters^5^, but the findings here are the first to show that traveling waves exist in the primate brain on much finer spatial scales. The traveling wave patterns shown in **Fig. 6A** have curved propagation patterns that roughly match the spatial scale of orientation pinwheels^77^ and individual cortical columns^78^.

Earlier findings on traveling waves suggest that these signals are coarse plane waves^4^, showing how broad areas of cortex propagate information dynamically and synchronize their activity. By demonstrating that traveling waves exist on a much finer spatial scale, our findings suggest that traveling waves may propagate along intricate cortical maps and provide the opportunity to study how traveling waves interact with detailed cortical gradients. Given our demonstration that information can be extracted for decoding at the tens-of-microns scale in V1, it would be interesting to explore whether BISC can achieve similarly high-resolution decoding in other brain areas relevant for human BMI applications, such as speech or motor movement decoding. The feasibility of such applications likely depends on the topological organization of the local circuits and the underlying task demands.

Although it was not included in these *in vivo* studies reported here, BISC supports amplitude controlled, bi-phasic current stimulation that can be configured to be either cathodic-first or anodic-first, followed by a passive balancing phase (see **Methods**, **Fig. S19 and S26**). The stimulation circuits in each of the 16,384 pixels can be independently programmed (see **Supplementary Discussion S1**), limited only by the aggregate current that can be sourced or sinked at any given time. Future experiments with BISC will test these stimulation capabilities.

While the current version of BISC relies on on-chip electrodes, the electronics of BISC can be easily connected to polyimide “extenders”, which would allow BISC to be adapted for intracortical depth electrodes, large area surface electrodes, or both. These extenders can be attached to BISC through solder-bump attachment, thermosonic bonding, or with the use of anisotropic conductive films. Multiple BISC devices can also be tiled over the brain surface to achieve larger area coverage without the use of extenders. All these alternatives build on the technology validated in this work, a fully encapsulated, integrated, wireless, bidirectional neural interface system that can be placed subdurally and controlled from an antenna outside the scalp. The effectiveness of decoding depends on several factors, including the dimensionality of the problem we intend to decode (such as complex, high-dimensional fine motor movements or speech), the spatial organization of the covered brain regions, and the information carried by the LFP for each electrode. Depending on the application and the complexity of the problem, the BISC system is scalable and modular and can be adapted to meet these diverse needs effectively.

The current headstage design is quite large, which makes it rather awkward for freely moving animal studies and future human use. A significant reduction in volume can be achieved with the current bill-of-materials by combining all the electronics into a single stacked PCB connected with dual radio-frequency cables to a wearable antenna. Further reduction in the size and form factor of the relay station will come in the design of an ASIC that replaces most of the discrete PCB components.

BISC overcomes one of the critical barriers to the widespread clinical translation of many alternative technologies – percutaneous wires – while delivering unprecedented volumetric efficiency and is very timely, given recent advances in the performance of BCI systems for speech and motor control ^79–81^. The combination of the high-density wireless recording and stimulation capabilities of BISC with deep-learning-based methods is poised to revolutionize high-bandwidth bidirectional BCIs and holds tremendous promise for revolutionizing treatments of brain diseases ranging from depression and aphasia to motor disorders, strokes, and blindness.

## ACKNOWLEDGEMENTS

This work was partly supported by the Defense Advanced Research Project Agency (DARPA) under Contract N66001-17-C-4001, the Department of the Defense Congressionally Directed Medical Research Program under Contract HT9425-23-1-0758, the National Science Foundation under Grant 1546296, and the National Institutes of Health under Grant R01DC019498. We would like to acknowledge the use of facilities and instrumentation at the Columbia Nano Initiative, the CUNY ASRC, and the UPenn Quattrone Nanofabrication Facility. We also extend our thanks to Youry Borisenkov, Adam Banees, and Kukjoo Kim at Columbia University for help with chip processing and Tjitse Vandermolen and Kenneth Kosik at UCSB for many helpful discussion.

## AUTHOR CONTRIBUTIONS

KLS, NZ, TJ, and RJC conceived the project. NZ, TJ, GE, MS, KT, GR, JH, KLS, and RJC designed the implant circuit. JDF, JK, and HY post-processed the implant. NZ andTJ implemented the relay station hardware. GE, NZ, PM, RJC, SP, TJ, AM, and LC implemented the relay station software. TJ, NZ, and SP performed bench-top characterizations. BY, ES, TJ, NZ, KLS, RH, IG, and GE performed *in vivo* experiments on the porcine subject. TJ, BY, and PC conducted porcine data analysis and histology. BP, AD, KEW, NZ, and TJ performed *in vivo* experiments on the motor cortex of non-human primate. TJ, BP, and KEW performed motor cortex data analysis. AT, SP, KLS, RJC, TJ, NZ, GE, TS, GJR, and CN performed *in vivo* experiments on the visual cortex of non-human primate. ZL, KW, AT, SP, DO, RJC, AD, EZ, and JJ performed visual cortex data analysis. KLS, AT, BP, MR, JJ, and DY acquired funding. KLS, AT, BY, BP, RJC, LPC, and JJ provided supervision. TJ, NZ, GE, KLS, KW, ZL, AD, EZ, JJ, and SP wrote the manuscript with review and editing contributed by all authors.

## DATA AVAILABILITY

All recorded electrophysiological data relevant to the figures presented in this paper are available at https://github.com/klshepard/bisc. All other relevant data are available from the corresponding authors upon reasonable request.

## CODE AVAILABILITY

All scripts used for data analysis are available at https://github.com/klshepard/bisc. All other relevant codes are available from the corresponding authors upon reasonable request.

## SUPPORTING INFORMATION

Supporting information includes Discussions S1 through S3, Figures S1 through S26, Tables S1 and S2, Supplementary Videos S1 through S9.

**Extended Data Fig. 1.**
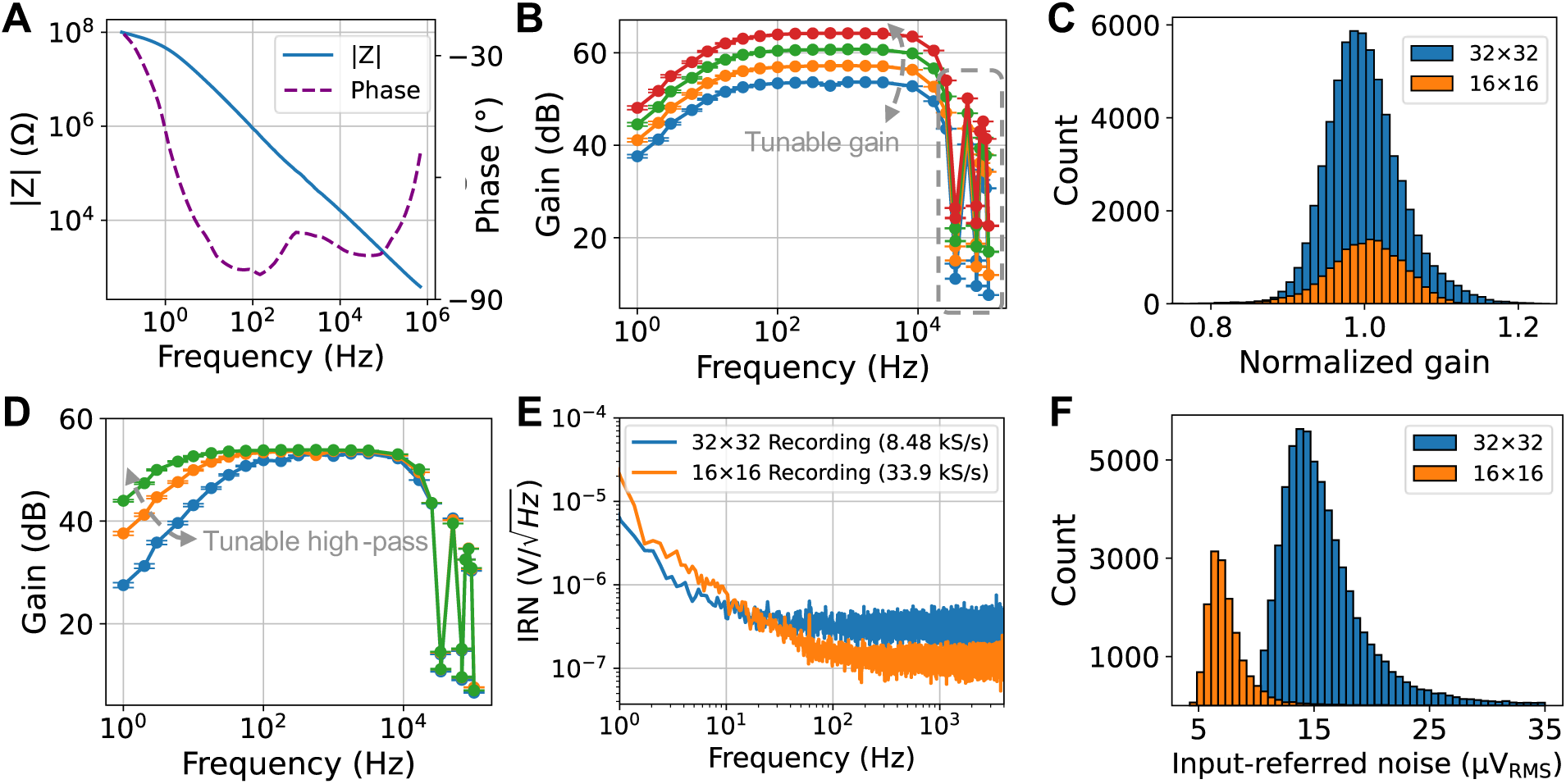
Bench-top i*n vitro* characterization of BISC implant. (**A**) Electrochemical impedance spectroscopy of titanium nitride electrode. (**B**) Frequency response across different gain configurations from a representative 16×16 recording. Note that gain is programmed through a single back-end amplifier that is shared by all pixels. Error bars indicate standard error (SE), and dashed rectangle marks the effects of boxcar sampling (flat band gains: 53.7 ± 0.20, 57.2 ± 0.21, 60.7 ± 0.20, 64.2 ± 0.19 dB, values: mean ± SD. n = 255, 255, 245, 235). (**C**) Histogram of channel gain variation for each recording mode (16×16 mode: ± 5.1%, 32×32 mode: ± 4.8%, values: SD. n = 15,163 and 62,245). (**D**) Frequency response across different high-pass (HP) filter configurations from a representative 16×16 recording. Error bars indicate SE (3-dB corner: 4.19 ± 2.28, 13.30 ± 2.37, 54.42 ± 1.98 Hz, values: mean ± SD. n = 244, 254, 256). (**E**) Input-referred noise (IRN) spectrum averaged over representative pixels (n = 10) for each recording mode. (**F**) Histogram of channel IRN for each recording mode, integrated from 10 to 4 kHz (16×16 mode: 7.68 ± 3.11, 32×32 mode: 16.51 ± 6.85 μV_RMS_, values: mean ± SD. n = 15,163 and 62,245).

**Extended Data Fig. 2.**
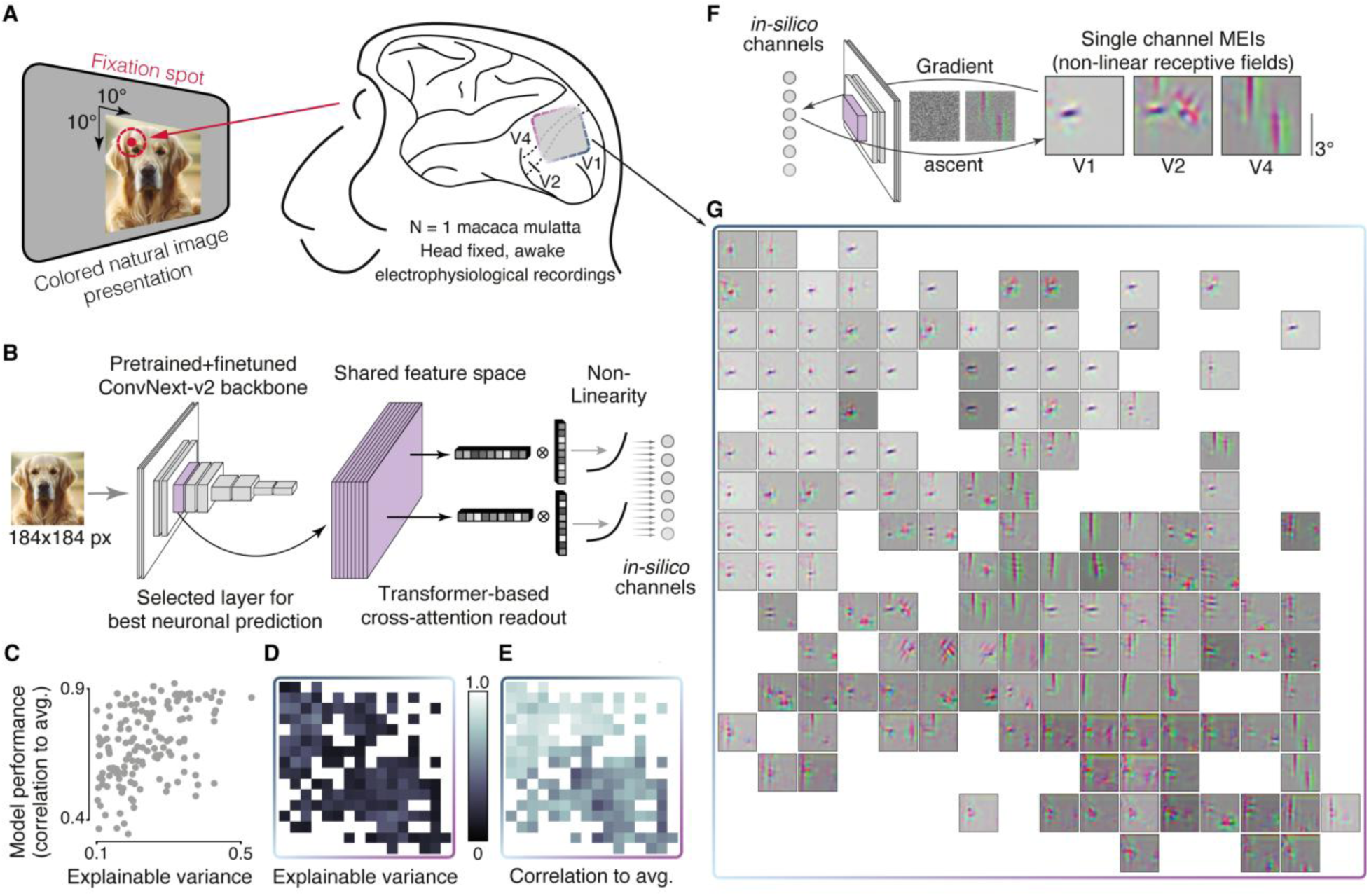
BISC recordings over visual cortex with natural images. (**A**) We presented static colored natural images, while the monkey was fixating (120 ms presentation time per image, 15 images per trial, 1200 ms inter-trial period). Images (10°×10°) were shown with their center 3° to the right and below the fixation spot. (**B**) Model architecture: The pre-processed stimuli (184 × 184 pixels) and neuronal responses were used to train a neural predictive model, which takes images as an input and outputs an estimate of the underlying neuronal activity. We passed the images through a ConvNext model, pre-trained on an image classification task to obtain image embeddings, i.e. a shared feature space. We then computed the neuronal responses by passing the feature activations to a transformer-based readout and a subsequent non-linearity. (**C**) Explainable variance, a measure of response reliability to natural images, plotted versus the model’s predictive performance (correlation between prediction and average neural response to repeated presentations) of all 144 channels (explainable variance 0.24 ± 0.09, and correlation to average 0.69 ± 0.14. values: mean ± SD). Only channels with an explainable variance greater than or equal to 0.1 are included in these analyses. (**D**) Visualization of the explainable variance as a function of channel position on the array (same spatial layout as in (D)). (**E**) Same as (D), but for the model’s predictive performance (correlation to average). (**F**) Schematic illustrating optimization of most exciting images (MEIs). By optimizing a random starting image to elicit the highest activity in one in-silico channel, the visual features that this channel is selective for can be exemplified. Three example MEIs from areas V1, V2, and V4 are shown. (**G**) MEIs for all 144 channels across the array which reliably responded to repeated image presentations. MEIs in area V1 are characterized by oriented Gabor filters, while the channels overlying area V2 and V4 exhibit more complex color opponent feature tuning.

## Methods

### BISC Implant Design

The BISC implant (**Fig. S1**), a custom-designed application-specific integrated circuit (ASIC) measuring 12 mm by 12 mm, was manufactured by the Taiwan Semiconductor Manufacturing Company (TSMC) in a 0.13-μm bipolar-CMOS-DMOS (BCD) process (**Fig. S1**). Post-processing steps, including thinning to 25 µm, electrode processing, and passivation, were conducted both in-house in the semiconductor processing facilities at Columbia University on coupons and at 200-mm wafer scale at MIT Lincoln Laboratories as described below.

The implementation of the implant followed a custom mixed-signal design flow. The digital circuits, which include the logic of the on-chip controller, were designed as finite state machines in SystemVerilog hardware description language (HDL). The HDL was subsequently synthesized into gate-level description using a logic synthesizer (Genus, Cadence Design Systems) and place-and-routed using a physical implementation tool (Innovus, Cadence Design Systems) using TSMC 0.13-μm-BCD standard-cell libraries. For analog circuits, schematic capture and layout tools (Virtuoso, Cadence Design Systems) were used for the circuit implementation. RF and microwave design tools (Ansys^®^ Electronics Desktop, Ansys; PathWave Advanced Design System, Keysight Technologies) were used for power coil and antenna designs.

The BISC implant features 65,536 surface titanium nitride (TiN) electrodes in its 16384-pixel array. We provide a basic overview of the design here with more details provided in **Supplementary Discussion S1**. BISC allows programmable selection of 1024 or 256 simultaneously recording electrodes from any of its electrodes with a set of rules.

- Every group of two-by-two electrodes (a pixel) is connected to an underlying pixel amplifier. Electrodes numbered from 1 to 65536 are connected to in-pixel amplifiers numbered from 1 to 16384 (sequentially from left to right and top to bottom). Electrodes 1, 2, 257, and 258 are connected to pixel-amplifier 1, and electrodes 65279, 65280, 65535, and 65536 are connected to pixel-amplifier 16384.
- A group of 16-by-16 in-pixel amplifiers can be used for each recording configuration. Both vertical and horizontal spacing (stride) of amplifiers in that group can be programmed individually from 0 to 7 pixel(s). For example, selecting a group with zero horizontal and vertical spacing (the densest spatial configuration) at the array’s top-left corner involves amplifiers 1 to 16, 129 to 144, …, and 1921 to 1936. In contrast, selecting a group with maximum spacing (seven horizontal and vertical) results in a configuration with the highest spatial coverage.
- Each electrode in a pixel can be either statically or dynamically multiplexed to the in-pixel neural amplifier. In static multiplexing, 256 electrodes (1 per pixel) are simultaneously read out at a sampling rate of 33.9 kS/s. In dynamic multiplexing, 1024 electrodes are read out simultaneously at 8.475 kS/s.

Each electrode is a 14-µm-by-14-µm square. In the densest configuration, with center-to-center electrode pitches at 26.5 µm by 29 µm, 32×32 electrodes cover an area of 0.85 mm by 0.93 mm. In contrast, the least dense configuration spaces the electrodes 424 µm by 464 µm apart, and 16×16 electrodes cover an area of 6.4 mm by 7.0 mm.

The reference input for the in-pixel neural amplifiers can be connected either to column-wise reference electrodes or to a global counter electrode; the latter acts as the ground reference potential for the chip. No significant differences were observed between these two schemes in our comparative evaluations. Consequently, the counter-electrode reference configuration was employed for all experiments conducted.

The in-pixel amplifiers are designed as chopped integrators, with their inputs biased by transistors in the weak-inversion region at the input. To avoid space-consuming DC blocking capacitors, the electrical double layer formed at the TiN electrodes is used as an implicit DC block. The pixel-amplifier inputs are biased to 0 V (ground) during operation. Chopping in the in-pixel amplifiers is used to reject intrinsic voltage offset and 1/𝑓 noise. The pixel amplifier outputs are multiplexed into a single two-stage programmable gain amplifier. The total voltage gain of the system can be programmed from 484 to 1620 V/V and high-pass corner can be programmed from 4 to 55 Hz. The low-pass corner of the system is 15 kHz. The outputs of the programmable gain amplifier are subsequently digitized by an 8.68 MS/s, interleaved 10-bit successive-approximation-register (SAR) analog-to-digital converter (ADC). When recording from 256 channels, the BISC implant has an input-referred noise of less than 5 µV_RMS_ across the frequency range of 10 Hz to 1 kHz and 10 µV_RMS_ from 0.3 Hz to 10 kHz. When recording from 1024 channels, the input-referred noise is less than 8 µV_RMS_ from 4 Hz to 1 kHz.

The implant also allows stimulation current driven from any group of electrodes with a set of rules.

- Stimulation can be driven from a minimum of one two-by-two electrode group (a pixel), up to the full array of 65,536 electrodes, acting as a macroelectrode.
- For monopolar stimulation, any combination of macroelectrodes can be used to drive current at a given time.
- For bipolar stimulation, macroelectrodes electrodes can be used to drive current either in-phase or out of-phase. Macroelectrodes driving opposite phases must have equal widths and have the same horizontal addresses, but their heights can be individually programmed.

The temporal profile for stimulation has three phases. The first two phases are anodic or cathodic, generating currents from two on-chip regulated current sources. During these phases, switches that connect the stimulation electrode to the corresponding current source are activated, allowing the current to flow from the source to the electrodes. Both anodic and cathodic current have the same amplitude which is programmable from 10 µA to 1.02 mA. The third phase is for charge balancing, in which the electrodes used for stimulation are grounded to ensure no accumulated charge on the electrodes. The duration of all three phases can be independently programmed from 0 to 350 µs. The compliance voltages of the anodic and cathodic current sources are ±1.4 V.

The BISC implant has a wireless transceiver with an on-chip slot monopole antenna, enabling wireless communication with the relay station. It operates as an ultra-wideband impulse radio, encoding digital data in short “bursts” of a 4-GHz sine wave. In this encoding scheme, a data “1” is represented by a burst, while a data “0” is the absence of a burst. The transceiver occupies up to 700 MHz of bandwidth in the unlicensed ultra-wideband frequency band and can support uplink data rates of 108.48 Mbps and downlink data rates of 54.24 Mbps. To allow full duplex communication using a single antenna, transmitting and receiving are time-division multiplexed for every bit. This allows us to control the timing precisely to stop a recording while the implant still transmits data to the relay station.

The implant features wireless power transfer that allows harvesting more than 64 mW from the integrated power coil through inductive resonance coupling. The AC power received on the powering coil is first rectified by an active rectifier and then regulated by regulators to support three 1.5-V power domains and one −1.5-V power domain.

### Relay Station Design

The relay station (**Fig. S2**) interfaces the BISC implant to a host computer, while providing wireless power to the implant, sending commands to control the implant’s behavior, collecting recorded data, and relaying everything to the host computer over wired or wireless Ethernet. The relay station has two parts: the headstage, where the wireless transceiver and power amplifier reside, and a processor module, upon which a Xilinx Zynq processor and logic translators reside.

The headstage is a wearable device (75 × 75 × 45 mm, 151 g) prototyped using off-the-shelf components and printed circuit boards. It establishes wireless communication with the BISC implant and supplies power to the implant through an integrated wireless transceiver and power amplifier. The headstage is connected to the processor module, which contains an FPGA (Snickerdoodle Black, Krtkl), with a standard HDMI cable which can be up to 5 m in length. The HDMI cable sends 12-V DC power to the power regulators on the headstage, delivering up to 1.2 A. The wireless transceiver on the headstage transmits high-speed digital data up to 108.48 Mbps using LVPECL over the differential twisted pairs on the HDMI cable to the processor board. A microcontroller (Teensy 4.1, PJRC) integrated on the headstage configures the wireless transceiver and the power amplifier. Communication between the microcontroller and the processor board is achieved using a two-wire UART protocol through the HDMI cable.

The BISC headstage is assembled from four separate printed circuit boards. The arrangement of the boards, from the bottom (facing implant) to the top, is as follows:

- *<u>UWB circular dipole antenna board.</u>* A dipole antenna is printed on a 0.2-mm-thick, single-layer FR-4 substrate. It is co-designed with the powering coil to optimize the bandwidth and radiation efficiency. The antenna board connects to the transceiver board with an in-series SMP adapter (19K104-K00L5, Rosenberger Group). A balun (BD3150N50100AHF, Anaren) is installed on this board to convert singled-ended RF signal from the SMP adapter to differential signal for the dipole.
- *<u>Power coil board.</u>* The power transmitting coil is a planar, square spiral, printed on a 1.6 mm thick, two-layer FR-4 substrate. It is fabricated with a 13 oz (0.455 mm) thick copper to implement a high-Q inductance for enhanced efficiency and minimal heat loss. The coil has an outer diameter of 3.6 cm and an inner diameter of 2.5 cm to allow clearance for the UWB dipole antenna. The coil has lumped impedance equivalent to 885 nH and 1.1 Ω with a self-resonance frequency well beyond (> 70 MHz) the link frequency. Measured linked efficiency is −10.5 dB at 1.5 cm distance, assuming an ideal conjugate matched driving source and a 75 Ω load at the receiver side which is equivalent to the overall circuit load. The coil is terminated with an edge-launch SMA connector and is driven by the power amplifier board via a short (75 mm) SMA cable.
- *<u>Transceiver board.</u>* The transceiver board is a 1.6-mm-thick four-layer printed circuit board. Rogers RO4003C is used as the top prepreg to allow good impedance matching and reduce power loss, while FR-4 is used as the core and bottom prepreg to lower the cost. Components installed on the transceiver board are for transmitting and receiving UWB pulses. The SMP connector to the antenna board, receiving chain, and transmitting chain are connected through a RF switch (HMC8038, Analog Devices). In the receiving chain, the RF signal first passes through a digitally controlled attenuator (HMC540SLP3E, Analog Devices) to prevent electromagnetic interference saturating the amplifiers. Subsequently, the signal is amplified 60 dB using two low noise amplifiers (CMD308P4, Qorvo) in series and filtered by a band-pass filter (B040MB5S, Knowles). An envelope detector (ADL6012, Analog Devices) then extracts the envelope of the incoming signal, followed by a threshold detector comprising two operational amplifiers (THS4304, Texas Instruments) and a comparator (TLV3604, Texas Instruments). The threshold detected signal becomes a digital short pulse if the implant is transmitting a “1” or stays constant if transmitting “0”. Multipath fading or reflection in the transmitting path can cause spurious pulses to be detected by the threshold detector. This issue is resolved using a self-resetting edge detector, which waits for 6 ns between detections to allow echoes to decay and resets the outputs after each detection. The edge detector is built from a LVPECL buffer (SY89327L, Microchip Technology), a D-flip flop (SY10EP51VMG, Microchip Technology), and a comparator (TLV3601, Texas Instruments). The final output from the edge detector is fed into a divide-by-2 divider (MC100EP32, Onsemi), converting the digital pulses into a non-return-to-zero data stream. This stream is then sent over the HDMI cable (RX) and sampled by the FPGA, and the original data is reconstructed by XORing the current and previous received bits. In the transmitting chain, a frequency synthesizer (ADF4351, Analog Devices) is used to generate the 4 GHz sine wave bursts. To transmit data “1”, the processor board sends a logic high through the HDMI cable (TX), and the RF output from the synthesizer is activated, driving the dipole antenna via the RF switch. The reference clock of the synthesizer and control signal for the RF switch are also sent over the HDMI cable (CLK, TR) by the processor board. Additionally, this board houses the microcontroller which controls the attenuator and the reference voltage of the threshold detector. DC power, reference clock and other control signals are passed to the power amplifier board via a board-to-board connector.
- *<u>Power amplifier board.</u>* The power amplifier board implements a class-E amplifier to drive the power coil at 13.56 MHz. The amplifier uses a single-ended gallium nitride (GaN) transistor (EPC2051, Efficient Power Conversion) as the active switch, loaded with a standard matching network ^82^ that shapes the impedance of the coil. In an inductively coupled system, link efficiency and tuning of its resonant frequency depend on a number of dynamic variables such as the coil-to-coil alignment, electromagnetic properties of the environment, operating mode of the implant etc. To compensate for these variables in real time, the board employs two feedback mechanisms to keep the link operating at its optimum. First, it controls the radiation magnitude by periodically reading out the level of power received by the implant and adjusting the supply voltage of the class-E amplifier through an I2C configurable regulator (TPS65400, Texas Instruments). Second, it prevents de-tuning of the resonance by adjusting the series capacitance of the load network using a reconfigurable capacitor bank ^83^. By monitoring the current consumed by the class-E amplifier under different load conditions, the 4-bit (1pF LSB) bank is configured to keep the link resonance stable at 13.56 MHz.

The processor module runs both firmware and the Linux operating system on the FPGA and ARM processor of a Zynq-7020 SoC. We designed application programming interfaces using Python programming language and Xilinx PYNQ libraries. Our Python software running on Linux (Ubuntu 18.04) provides methods to fully control the BISC implant, stream the recording data over ethernet or store it on the secure digital (SD) card, and control the microcontroller on the BISC headstage.

The BISC headstage and processor module presently only work with one implant at a time. However, multiple devices can be implanted if a spacing of at least 5 mm is maintained between devices. In this scheme, an implant would be selectively powered up by positioning the headstage over it.

### BISC implant post-processing

#### Coupon processing

A coupon consisting of four reticles (16 total die) was scribed out from the original 300-mm BISC wafer from TSMC using a diamond scribe. **Fig. S3** shows a schematic of the process flow through a cross-section of the microelectrode array (MEA) region. A layer of photoresist (AZ P4620, MicroChemicals) was spin coated onto the coupon at 3000 rpm, exposed on a contact mask aligner (MA-6, Karl Suss), and then developed (AZ 400K, MicroChemicals). Etching the silicon oxide and nitride layers with a plasma etcher (Oxford Instruments) exposed the Al redistribution layer (RDL) at the electrode sites. The RDL was wet etched in Al etchant (Type A, Transene) to expose the underlying Ta diffusion barrier layer over the redistribution via (RV) layer. A brief Ar ion sputter to remove the Ta oxide layer was followed by sputtering in the same vacuum chamber (Orion 8 Dielectric Sputter Chamber, AJA International) of a 240 nm layer of titanium nitride (TiN; 120 min, 0.33 Å/s, 20 sccm Ar, 3 mTorr, 175W) from a TiN target onto the Ta, and the resist was lifted off in photoresist remover (Remover PG, Kayaku).

TiN was chosen as the electrode material due to its biocompatibility, its rough surface morphology providing a reduction in impedance (Z = 205 kΩ at 1 kHz for a 14 µm × 14 µm electrode) compared to a smooth electrode material such as gold (Z = 3.1 MΩ at 1 kHz for a 14 µm × 14 µm electrode), its strong adhesion for chronic implantation and stimulation, its compatibility with CMOS processing, and its capacitive non-Faradaic current properties ^84^. Characterization results of the BISC TiN electrodes are shown in **Fig. S4**.

After electrode fabrication, a 2.5-µm-thick polyimide encapsulation layer (PI2610, HD Microsystems) was spin coated onto the front surface after functionalizing the surface with an adhesion promoter (VM652, HD Microsystems) and cured at 350 °C for 30 min. The TiN electrodes were exposed using oxygen plasma etching (Oxford Instruments). After a brief Ar ion clean, a second 360 nm TiN layer was sputtered on top of the first TiN layer using the same deposition parameters, and the AZ P4620 mask was lifted off in Remover PG. The die were then separated with a dicing saw (DISCO Corporation) and bonded frontside down on a glass carrier using an instant adhesive (Loctite^®^ 460, Henkel) for thinning on a grinding and polishing tool (X-Prep^®^, Allied High Tech) to a final silicon substrate thickness of ∼25 µm. Chips were then loaded into a parylene coating chamber (Specialty Coating Systems) to encapsulate the backside with 10 µm of parylene C. The parylene was trimmed along the edge to leave about a 1 mm overhang for handling and the chip was released from the glass carrier by dissolving the adhesive in acetone.

#### Wafer-scale processing

Whole 8” wafers can be processed in a similar way to individual reticles starting with TiN electrode fabrication and polyimide front surface encapsulation. Assuming the entire wafer is dedicated to BISC chips, a single wafer yields up to 200 fully contained devices ready for sterilization and implantation (**Fig. S5**).

During mechanical thinning, grinding induced defects are introduced into the silicon which can serve as nucleation sites for bending induced fracture growth and device failure. Extensive polishing is needed to remove these defects. With this in mind, for the wafer scale thinning process, we developed a wet etch thinning technique using an isotropic silicon wet etch consisting of hydrofluoric, nitric, and acetic (HNA) acids. The device wafer was bonded to a silicon carrier wafer using an organic adhesive (WaferBond^®^ HT-10.11, Brewer Science). The wafer edges were clamped using an O-ring seal, and the wafer surface was covered in etchant which was stirred and maintained at room temperature to give a uniform etch rate of 4.28 µm/min across the wafer. The remaining silicon thickness was monitored during etching, and the wafer was removed and rinsed in DI water once this value reached 25 µm.

The chips were carefully aligned using features visible through the thinned backside of the wafer and singulated by laser dicing (DISCO Corporation) through the thinned wafer with 100-µm-wide dicing lanes, stopping within the WaferBond adhesive layer. A 10-µm-thick film of parylene C was deposited onto the backside of the wafer in a room temperature chamber (Specialty Coating Systems) using an adhesion promoter (Silane A174, Sigma-Aldrich). The wafer was then loaded into an excimer laser dicing tool (IPG Photonics) and was aligned as before. The parylene within the dicing lanes was laser cut with a dicing width of 60 µm to leave a 20 µm overhang of parylene along the chip edges. The wafer was then submerged in WaferBond remover (1-dodecene, Brewer Science) to dissolve the underlying adhesive. The individual chips were collected and rinsed in acetone and isopropanol and dried with nitrogen.

### Mechanical flexibility of implanted devices

To render BISC chips compatible with *in vivo* implantation (i.e., skull closure with minimal tissue damage and displacement and maximal conformability), we removed the bulk of the rigid silicon substrate to reduce volume and achieve sufficient mechanical flexibility. Any desired thickness of silicon can be removed using the mechanical or wet etch thinning processes described here. We identified a 10-μm thickness limit based on the depth of an n-type buried layer in the CMOS process. In the case of mechanical thinning, consideration must be made for grinding induced subsurface damage. We found that ∼25 µm was a safe stopping point to guarantee chip functionality while also providing sufficient flexibility, while the 15 μm thickness was achievable with wet etching.

#### Device mechanical stiffness

The final implanted device is a thin plate comprising a multi-material stack of polyimide, metal interconnects and vias surrounded by interlayer dielectrics (BEOL), the silicon substrate, and finally a parylene backing layer. By summing the stiffnesses of each of the four layers, we can estimate a total device bending stiffness of ∼130 µN·m according to:

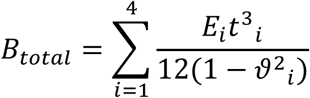

where *E* is Young’s modulus, *t* is the layer thickness, and 𝜗 is Poisson’s ratio ^85,86^.

By encapsulating our device in polyimide and parylene, we reduce the material stiffness mismatch between the device surface and the surrounding tissue. The main implication of reduced *device-scale* stiffness which we achieve from die thinning lies in device tissue conformability (see discussion below). Reduction in tissue damage along the device edges (see histological results), may also be aided by reduced device stiffness.

#### Device conformability to cortex

Three regimes can be considered to understand what happens mechanically to BISC upon implantation. These regimes depend on the relative magnitudes of the curvature of the cortex (*r_cortex_*) and the elasto-capillary length (*L_EC_*) of the device. We model the BISC implant as a flexible device with hydrophilic surface and no built-in stress, wrapping around a cylinder with water as the wetting liquid (surface tension, 𝛾 = 72 mN/m). Through equating surface energy reduction with bending energy increase, we find that *L_EC_* is given by:

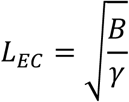

where *B* is the bending stiffness. The radius, *r_min_,* below which spontaneous wrapping does not occur is given by:

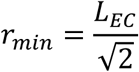

Regime 1 (*L_EC_* < *r_cortex_*). In this regime, the device spontaneously conforms to the cortical surface driven by capillary forces. Take for example a 5-µm-thick parylene device for which *B* equals ∼0.035 µN·m, giving *L_EC_* = 0.7 mm and *r_min_* = 0.5 mm. In this case, the device conforms to even the smallest features such as sulci in the cortex, with curvatures down to ∼1 mm.

Regime 2 (*L_EC_* > *r_cortex_*). In this regime, the device does not spontaneously bend to conform to the cortical surface. In the limit of zero flexibility, the device may be pushed into the compliant tissue by an outside force, e.g., skull reattachment, potentially leading to significant tissue damage especially as device volume increases.

Regime 3 (*L_EC_* ≈ *r_cortex_*). This regime represents the BISC devices (*L_EC_* ≈ 40 mm, *r_min_* ≈ 30 mm) in the brain regions studied in this paper (*r_cortex_* ≈ 30 mm). As a demonstration of BISC conformability, **Fig. S6A** shows a BISC chip spontaneously wrapping around a glass beaker with a radius of curvature of 30 mm with water as the wetting liquid. Importantly, this natural conformability indicates that outside mechanical forces are not needed push down the BISC device to make it conform to the cortical surface.

#### Bending induced fracture

Due to the nature of its atomic bonding, silicon is inherently prone to brittle fracture. For ideal silicon samples with minimal defects, the maximum sustainable bending strain (ε_𝑚𝑎𝑥_) is ∼1% ^87^. The strain in a material bent under uniaxial stress applied to a radius of curvature *r* is given by

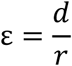

where *d* is the distance between the neutral plane and the plane of interest ^88^. Setting ε = ε_𝑚𝑎𝑥_ and taking *d* equal to the distance of maximum strain furthest away from the neutral plane in a slab of thickness *t*, i.e., *d* = *t*/2, we can calculate the smallest achievable radius of curvature *r* = 𝑟_𝑓𝑟𝑎𝑐𝑡𝑢𝑟𝑒_for an ideal silicon sample via

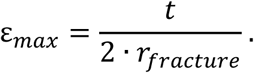

For a multimaterial stack, *d* should be taken as the distance from the neutral plane where strain is zero. For the BISC device, denoting *h* = 0 as the top of the polyimide on the front surface of the device, we can estimate the neutral plane position assuming zero slip and no debonding between adjacent layers to be at *h* = 20 μm, which is 5 μm into the silicon layer ^89^. This produces asymmetry in the mechanical stability of the device under opposing bending directions, since the silicon is more likely to fracture under tension than compression. When the parylene is under tension, 𝑟_𝑓𝑟𝑎𝑐𝑡𝑢𝑟𝑒_is calculated to be 1.5 mm, while when the polyimide is under tension, the value is 0.5 mm.

The maximal bending strain can be greatly reduced by defects in the sample and depends on the details of device processing. The effect of defects on device frature is typically quantified using Weibull distrubiton ^90^. For our process, curvatures as small as 10 mm 𝑟_𝑓𝑟𝑎𝑐𝑡𝑢𝑟𝑒_ are achievable (**Fig. S6B**). However, such small radii of curvature should not normally be encountered during or after implantation (see section above on conformability); once implanted, the device is not subject to any macroscopic bending lengths that would result in fracture. This is consistent with our experiments, which have never witnessed a fracture mode failure *in vivo*. Handling during implantation is the only point at which fracture could occur. However, surgical procedures allowed for consistent implantation without requiring the device to be bent beyond the minimal radius of curvature.

### Porcine somatosensory cortex recording experimental procedure

All porcine experiments were performed in compliance with the National Institutes of Health Guide for the Care and Use of Laboratory Animals and were approved by the Columbia University Institutional Animal Care and Use Committee (protocol AC-AABQ0559).

#### Surgical procedures

The BISC implant chips and all surgical tools were sterilized with ethylene oxide (EtO) prior to implantation. The surgical incision and craniotomy were customized to the number of devices and implant target. For the porcine model sensorimotor cortex experiment, two BISC devices were implanted over sensorimotor cortex of each hemisphere of a female Yorkshire pig (18 weeks old, 23.9 kg). The procedure was performed under general anesthesia. For placement of bilateral devices, a 5-cm curvilinear bicoronal incision was planned just rostral to the anatomic sensorimotor cortex. The incision was injected with 2% lidocaine, incised sharply and the scalp reflected posteriorly. Following hemostasis, a bifrontal craniotomy was performed. Burr holes were placed with slotting over relevant venous structures and connected with a cutting drill. Then, the bifrontal craniotomy bone flap was elevated, and epidural hemostasis was obtained. For each BISC chip, the dura was tented and a coronally oriented 15 mm linear incision was made to allow access to the subdural space. A commercial four-contact subdural strip electrode (PMT Cortac^®^ 2110-04-032, PMT^®^ Corporation) was used in conjunction with commercial EEG acquisition equipment (XLTEK Protektor32, Natus Neuro) and peripheral subdermal needle electrodes to identify the location of somatosensory cortex ^55^. The same strip was used as a shuttle to slide our implant, from rostral to caudal in the subdural space targeting the cortical location of maximal somatosensory evoked potential (SSEP) response (**Fig. S7**). The relay station was then draped and brought into the sterile field to test the device prior to dural closure. Once recording and stimulation function were confirmed, the dura was reapproximated and closed in watertight fashion with 4-0 Nurolon sutures. The craniotomy bone flap was microplated back into place with the nonmetallic cranial fixation system (Delta Resorbable Implant System, Stryker). The galea was closed with 3-0 Vicryl sutures and the skin with 3-0 nylons and surgical glue. The animal was recovered under veterinary observation.

#### SSEP recording

Two weeks after the implant, SSEP was recorded from the subject (28.5 kg) under stable anesthesia. Drug dosages of 17.5 – 28.1 µg·kg^-1^·h^-1^ Fentanyl, 4.9 mg·kg^-1^·h^-1^ Ketamine, 1.6 – 2.6 µg·kg^-1^·h^-1^ Dexmedetomidine were administered. The choice of drug was influenced by the subject’s allergic reaction to Propofol.

Subdermal needle electrodes (RLSND110-1.5, Rhythmlink^®^) were used in conjunction with a commercial stimulator (XLTEK Protektor32, Natus Neuro) to deliver electrical stimulation. Needles were placed on median nerve and four different snout locations (Fig. 3A) with local reference pairing on the contralateral side of the implant. To invoke SSEP, we used a train of anodic pulses continuously running at 2.79 Hz with pulse width of 0.3 ms. Stimulation amplitude was gradually increased with increments of 0.5 mA until distinct twitching responses were observed (**Fig. S8B**). For each location, we collected approximately 200 SSEP trials, where a trial refers to the delivery of each anodic pulse. Recording configuration used 16×16 channels spatially programmed in the sparsest recording mode, minimum programmable gain amplifier (PGA) gain of 53.7 dB, and high-pass filter cut-off of 13 Hz.

#### Histology

Five weeks after the implant, we extracted the brain post-mortem for histological analysis. Coronal sections were taken from under the BISC electrode, near the rostral edge of the implant where there was gross evidence of a mechanically induced injury, and from the ipsilateral occipital lobe as a control. Sections were stained with three different markers: hematoxylin & eosin (H&E, H: NC2072425, Fisher. E: 6766008, Fisher), NeuN (MAB377, Millipore), and Iba1 (NB100-1028, Novus Biologicals) (**Fig. S9**). Although a small lesion was found near the rostral edge of our implant indicative of a mechanically induced placement injury, tissue section from directly under the electrode array showed no significant pathological change except for a mild microgliosis.

### Porcine somatosensory cortex recording data analysis

#### Data pre-processing

Non-functioning recording channels were identified from the baseline recordings that were taken without running any electrical stimulation. They were identified by labeling channels whose waveform root mean square (RMS) values are below a heuristic threshold of 1.7-bits after low-pass filtering (300 Hz, eight-order zero-phase Butterworth) or whose waveforms remained consistently saturated throughout the recordings. The BISC 10-bit ADC has an output range of [0, 1023]. For porcine SSEP data analysis, we defined the non-saturated code range for the BISC ADC as [3, 1020] with any data point outside of this range considered saturated. By this means, 25 out of 256 channels were labeled as either dead or saturated and were eliminated from further analysis.

For preprocessing, recordings were first divided into 500 ms segments for each SSEP trial centered on the time of stimulus delivery (from −250 to 250 ms with the stimulation at t = 0) and were linearly detrended. For each trial, a channel recording was marked as invalid if its segmented waveform contained one or more saturated data points with invalid segments discarded from further analysis. Common average referencing was applied by re-referencing against the mean of 10 channels with the lowest average RMS activity across the five different stimulation locations. Recordings from these 10 channels remained consistently non-saturated throughout the experiment.

For synchronization between the commercial stimulator and the BISC system, the stimulator was programmed to deliver a 0.3-ms wide electrical pulse to the BISC relay station 55 ms after each peripheral stimulation. Offline analysis, however, revealed that electromagnetic radiation from these pulses were recorded as significant artifacts affecting segments from 50 to 100 ms. For visualization purposes, these artifacts were removed by taking singular value decomposition of recording segments from 50 to 100 ms and then removing a heuristically chosen number (n = 15) of principal component time series from the original data. This artifact removal, however, only affects the plot visualization in Fig. 3C and Fig. S8C. It does not affect any other visualization or analysis presented in this paper, as they do not utilize recording segments beyond 50 ms. As the last pre-processing step, all valid segments were downsampled to 2.11kS/s with anti-aliasing filtering with the mean of baseline window (from −50 to −5 ms) then subtracted.

#### SSEP visualization and analysis

Time-domain SSEP waveforms from 0 to 50 ms (Fig. 3B**, Fig. S8A**) were plotted by averaging the channel recordings over 100 trials with channels with fewer than 100 trials excluded from this analysis. Waterfall plots (Fig. 3C**, Fig. S8C**) are a different representation of the same trial-averaged data (n = 100), linearly detrended and aligned to the shared time axis from −50 to 100 ms. They were generated by arranging the mean channel responses sorted by their peak amplitudes inside the SSEP window, defined to be from +5 to +45 ms. The peak amplitudes here are either positive or negative extrema, whichever is higher in absolute magnitude. Colored spatial maps (Fig. 3D) show the peak response of each channel inside the SSEP window, after normalizing by the overall peak response of the array. Frame-by-frame temporal dynamics of these spatial maps are provided as a movie (**Video S1)**.

For further analysis, only channels (n = 158) that remained consistently non-saturated throughout the experiment were considered. 100 valid trial recordings from each stimulation location were first z-scored. Then, their lower-dimensional features were extracted through principal component analysis (PCA), taking only components that cumulatively explain 80% of the variance. These features were further reduced to two-dimensional space (Fig. 3E) by applying t-distributed stochastic neighbor embedding (t-SNE; Barnes-Hut method, perplexity = 30). The same features were used to train a classifier using linear discriminant model (LDA; singular value decomposition method) with stratified 10-fold cross validation (Fig. 3F). All analyses in this section used Python v3.10.0, numpy v1.23.5, scipy v1.10.0, and scikit-learn v1.2.1.

### NHP motor cortex recording experimental procedure

All NHP motor cortex experiments were performed in compliance with the National Institutes of Health Guide for the Care and Use of Laboratory Animals, and were approved by the University of Pennsylvania Institutional Animal Care and Use Committee (protocol 807341).

BISC device was attached to the artificial skull referred to as the BrainPort (Fig. 4B) using clear epoxy (Loctite^®^ EA M-31 CL, Henkel) and was EtO sterilized after curing. Registration of the device (Fig. 4C) was determined by comparing the ground truth images of the BrainPort over the cranial window, before and after the device attachment.

During behavioral tasks, our NHP subject, a male rhesus macaque (Macaca mulatta, 11 years old, 13 kg) was seated in a primate chair with its head motion restricted but otherwise free to move. It was trained to reach and grab the target using its right arm which was contralateral to the device placement. The target in our experiment was a wooden stick referred to as the wand, manually held near the full reach distance of the subject. Post-experiment, we discovered that there may have been too little variation in the wand position. As a consequence, the subject’s normalized wrist velocities in y (front-back) and z-direction (up-down) were found to be highly correlated (Pearson r = 0.96). It may also be the reason that our feature decoder does not perform well in x-direction (left-right), as the wand was consistently presented in proximity of the median plane of the subject.

The reaching tasks were asynchronous and did not require an explicit cue, and the subject received food rewards for correct behaviors. Experiment was broken down into multiple sessions with rests between them, with each recording session lasting up to two minutes. Because the subject was not water restricted, it did not engage in tasks most of the time, resulting in 44 clean reaches over aggregate duration of approximately six minutes.

Three machine vision cameras filmed the experiment from three different angles. An open source deep learning framework, DeepLabCut ^91^, was used to label the wrist position of the subject at 60 Hz frame rate.

### NHP motor cortex recording data analysis

#### Data pre-processing

Position tracking was lost at times when the subject’s wrist moved out of the camera field-of-view, particularly when the arm was retracted beneath the metal plate of the primate chair. Short time gaps (< 0.5 s) in the position data were linearly interpolated, and the resulting data was smoothed using a moving average filter with 200-ms window. Remaining gaps were categorized into medium (< 2 s) and long (> 2 s) gaps, where medium gaps were interpolated using cubic spline function and long gaps were linearly interpolated. To extract velocity, derivatives of this data were taken and then smoothed again using a moving average filter with 200-ms window. Finally, the resulting wrist velocity in each dimension was upsampled to 200 Hz and then normalized by dividing by its standard deviation.

Throughout the experiment, BISC recordings were taken from 16×16 channels spatially programmed in the sparsest mode that covers the whole array, each channel sampled at 33.9 kS/s, with gain of 53.7 dB, and high-pass filter cutoff frequency of 13 Hz. For motor data analysis, we defined the non-saturated range as [3, 1020] out of the full ADC code range of [0, 1023]. For each session, a channel is considered non-saturated if less than 1% of data points are saturated.

Time points with packet losses were filled in by time domain linear interpolation, up to 11 consecutive samples (< 0.33 ms). Short segments up to three consecutive saturated samples (< 0.09 ms) were also linearly interpolated. Then, saturated channels were spatially imputed by taking the average of non-saturated channels in their immediate vicinity (Euclidean distance = 1) only if more than one non-saturated neighboring channels existed. The resulting data was band-pass filtered (0.3 – 300 Hz) with an eighth order zero-phase Butterworth filter and downsampled to 2.12 kS/s. Across sessions, an average of 15.0 ± 4.58 standard-deviation (SD) channel recordings were recovered through spatial imputing. Unrecovered saturated channels (45.8 ± 13.1 SD) were visualized as carrying all-zero data (Fig. 4H) and excluded from further analysis.

Unlike our other *in vivo* experiments, our motor cortex recordings contained significant hemodynamic artefacts, with a dominant frequency around 3 Hz and its harmonics. To remove this rhythm, the downsampled recordings were band-pass filtered in the frequency range of hemodynamic rhythm (3 – 12 Hz) with an 8^th^ order zero-phase Butterworth filter. The filtered recordings were treated as a space-time matrix which was decomposed using singular value decomposition (SVD). The resulting principal component (PC) time series were ranked according to their 3-Hz spectral power strength. Finally, a heuristically chosen number of PCs (five ranked components) were subtracted from the original downsampled recordings.

By means of this PC removal, we were able to preserve spectral content in the affected frequency range (**Fig. S10B**) which would not have been possible with a more aggressive approach such as band-stop filtering. **Fig. S10A** shows an example channel (address: (3, 8)) recording after hemodynamics removal. Spectrogram was computed by extracting spectral power through multitaper estimation (200 ms window, 10 Hz half-bandwidth) and then z-scoring for each frequency bin (0, 20, …, 200 Hz).

#### Motor feature decoder

To build a continuous motor feature decoder, as described in the main text, the recordings were further grouped into 4 frequency bands (LMP, β, low γ, high γ) and 19 time history bins (t-0.47, t-0.42, …, t+0.47 s). For model construction, only channels that were consistently non-saturated across sessions were used (n = 180), resulting in a 13,680-dimensional spatio-temporal-frequency vector 𝑋(𝑡) (180 channels, 19 time lags, four frequency bands) as model input for predicting motor feature 𝑦(𝑡). Time resolution of feature decoding was kept at 100 ms.

We used a linear partial least squares regression (PLS) model whose hyperparameter – number of PLS components – was determined by finding the minimum point of predictive sum of errors (PRESS) across five-fold cross validation (**fig. S10C**). We built our model to decode wrist velocity of all three dimensions and evaluated its performance using Pearson’s correlation coefficient. It performed best on decoding y-direction velocity (0.53 ± 0.04 SE) and next on z-direction velocity (0.50 ± 0.04 SE). Velocity in x-direction could not be decoded reliably; speed was decoded instead (0.50 ± 0.06 SE).

The y-direction velocity decoder was further analyzed by computing the relative contribution of each frequency band. Denoting our model as the following linear combination where ε(t) represents the residual error,

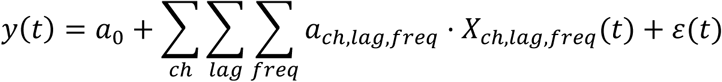

the relative contribution of each band 𝑤_𝑓_ was calculated as follows (Fig. 4F):

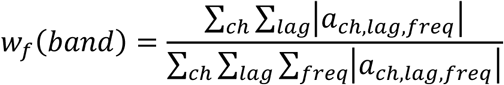

All analyses in this section used Python v3.10.0, numpy v1.23.5, scipy v1.10.0, and scikit-learn v1.2.1.

### NHP visual cortex recording experimental procedure

#### Surgical procedures

All experimental procedures were approved by the Baylor College of Medicine Institutional Animal Care and Use Committee (protocol AN-4367). Here, all behavioral and electrophysiological data were obtained from a healthy, male rhesus macaque (Macaca mulatta) monkey aged 19 years and weighing 13.85 kg during the study period. The animal was housed individually in a large room adjacent to the training facility, along with around ten other monkeys, permitting rich visual, olfactory, and auditory interactions, on a 12-hour light/dark cycle. Regular veterinary care and monitoring were provided.

All surgical procedures on monkeys were conducted under general anesthesia, adhering to standard aseptic protocols. Prior to surgery, the monkey had been fitted with a custom titanium headpost, which secured the head in a fixed position to facilitate accurate eye movement tracking. Pre-medication involved administering atropine (0.05 mg/kg) followed by sedation using a combination of ketamine (10 mg/kg) and dexmedetomidine (0.015 mg/kg). Anesthesia during surgery was maintained with isoflurane, adjusted between 0.5% and 2% as necessary.

The surgical site was prepared in a stereotaxic frame, and the implantation area was meticulously cleaned with alternating applications of betadine and alcohol prior to the initial incision. Based on precise stereotaxic coordinates targeting the primary visual cortex and higher visual areas V2 and V4, an incision was made through the epidermis to expose the underlying muscle. Large, curved hemostats were applied to clamp the muscle tissue above the intended removal site to enable the relay station headstage to be positioned on top of the skin close to the BISC implant (1.5 to 2 cm distance between top of the skin and the implant). Following 15 seconds of clamping, the targeted muscle was excised using a #10 blade scalpel.

A craniotomy measuring 2.5 cm by 2.3 cm was performed to access the underlying brain area for the BISC device implantation. The removed bone was preserved in sterile saline. Prior to dura mater incision, the animal was hyperventilated to maintain CO_2_ levels between 25-30 mm Hg. A 1.8 mm straight incision was made in the dura mater close to the posterior ridge of the craniotomy to create a pocket for the BISC device, which was then carefully slid under the dura with the electrode side facing the brain surface. A commercially available deep brain stimulation guide was used to accurately slide and position the BISC chip under the dura. The implanted device’s functionality was confirmed through successful communications with the headstage.

After testing the communication between the headstage and the chip, the dura was closed using 3-0 polyglycolic acid suture, and the previously removed bone flap was repositioned. Burr holes were drilled into both the bone flap and adjacent skull, and suture was used to secure the bone in place, promoting a flat, well-aligned healing surface. The overlying skin was then repositioned, and the device functionality was confirmed once more before suturing the skin closed with 3-0 braided vicryl sutures.

Post-operatively, the animal was weaned off isoflurane and closely monitored during recovery by both lab and veterinary staff at Baylor College of Medicine. Analgesics were administered for up to seven days post-surgery to manage pain.

The relatively straightforward surgical procedure and minimal invasiveness of the BISC chip enable us to replace the BISC chip after months by following the same procedure as above: recutting the dura over the old incision, removing the chip with graphite forceps, and then using the DBS guide to reposition a new chip. The data presented here from the visual cortex are from one such replacement chip

#### Electrophysiological recordings

Electrophysiological data from BISC were recorded by a custom LabVIEW (National Instrument) software system. Data were transferred over an Ethernet network connection from the relay station processor module to a Windows PC, while the chip was configured to record in either 16×16 electrodes full chip coverage mode or 32 × 32 electrodes dense coverage mode. Full chip coverage mode was recorded at about 34 kS/s per electrode whereas the dense coverage mode was recorded at about 8.5 kS/s per electrode. The LabVIEW software system implemented a trial-based state machine responsible for initiating and terminating behavioral trials, tracking eye movements and behavior, dispensing reward, controlling visual stimulation, time stamping and database logging various events for post-acquisition synchronization and analyses. A real-time display of acquired signals is also available in LabVIEW. A pseudorandom digital signal is recorded together with the BISC data, and the same signal is separately recorded together with the synchronizing photodiode signal described below. The presence of the pseudorandom signal in both recordings allows for the precise synchronization of BISC data with visual stimulation.

Visual stimuli were rendered on a dedicated graphics workstation and displayed on a 16:9 HD widescreen LCD monitor (23.8”) with a refresh rate of 100 Hz at a resolution of 1920 × 1080 pixels and a viewing distance of 100 cm (resulting in ∼ 63 px/°). The center of the screen was aligned with the monkey’s root of the nose. A photodiode was attached in the top-left corner of the display monitor and utilized to detect a synchronizing signal presented together with the visual stimuli. In experiments conducted with gray scale stimuli, the monitor was gamma-corrected to have a linear luminance response profile.

In experiments conducted with color stimuli, the monitor’s color channels were calibrated by measuring their spectra at 16 contrast levels equally spaced in the range from 0 to 255, without equalizing the outputs ^92^. The spectra were corrected for the spectrometer’s (USB2000+, Ocean Optics) sensitivity, allowing us to compare the relative intensities accurately. Intensity values for each color were derived by integrating the spectra over a 200 nm range. These data helped develop contrast-intensity curves and allow for the computation of an inverse gamma curve for linearization using a unified look-up table. After linearizing, the spectra were re-measured to confirm the uniformity of the color outputs. Mixed color outputs were also examined by testing all combinations of the three-color channels at four contrast levels, and the resultant spectra were documented. Finally, we compared the measured intensities from the mixed tests against predicted values based on single-color data. The strong correlation confirmed the linearity and minimal interaction between the channels, validating the calibration process. This streamlined approach ensures accurate color reproduction, essential for reliable experimental results.

A camera-based, custom-built eye tracking system verified that the monkey maintained fixation within ∼ 0.95° around a ∼ 0.15°-sized red fixation target. Offline analysis showed that monkeys typically fixated much more accurately. After the monkey maintained binocular fixation for 300 ms, a visual stimulus appeared. If the monkey fixated throughout the entire stimulus period, they received a drop of juice at the end of the trial.

#### Grating experiment

In this experiment, the monkey maintained binocular fixation on a red 0.16° fixation spot at the center of the screen for at least 300 ms to initiate a trial. In each trial, 40 static gratings of random orientations were presented consecutively (Fig. 5B) on a uniform gray background with a grating size of 6°. The fixation spot was at the center of the screen and the grating was centered 2° to the right and 2° below the fixation spot. Each grating had a normalized background luminance of 0.5 and contrast of 0.95 and was presented for 50 ms. The monkey was rewarded with juice if it maintained fixation during the whole trial which lasted 1500 ms.

#### Dot-mapping experiment

In this experiment, the monkey also maintained binocular fixation on a red 0.16° fixation spot at the center of the screen for at least 300 ms to initiate a trial. In a trial, a single 0.51° dot was presented on a uniform gray background, changing location and color (black or white) randomly every 50 ms (Fig. 6A). The fixation spot was at the center of the screen and the dots were presented in a rectangular field (6°×6°) centered at 1.5° to the right and 3° below the fixation spot. The monkey was rewarded with juice if it maintained fixation during the whole trial which lasted 1500 ms.

#### Natural image experiment

We sampled a set of 24,075 color images from 964 categories (∼ 25 images per category) from ImageNet ^65^ and cropped them to keep the central 420 × 420 pixels. Images were scaled up to 630 × 630 pixels using bilinear interpolation. All images had eight-bit intensity resolution (values between 0 and 255) in each color channel. We then sampled 75 as the test set with the remaining 24,000 images as the training set. The monkey maintained binocular fixation on a red 0.16° fixation spot at the center of the screen for at least 300 ms to initiate a trial, and images were shown at a size covering 10° (screen resolution of 63 pixels per visual angle) centered at 3° to the right and below the fixation spot. The rest of the screen was kept gray (128 intensity). During a recording session, we recorded ∼1000 successful trials, each consisting of uninterrupted fixation for 2.4 seconds including 300 ms of gray screen (128 intensity) at the beginning and end of the trial, and 15 images shown consecutively for 120 ms each with no blanks in between. Each trial contained either training or test set images. We randomly interleaved trials throughout the session so that our test set images were shown ∼40 times. The training set images were sampled without replacement throughout the session, so each image was effectively shown once or not at all.

### NHP visual cortex recording data analysis

#### Channel selection

Prior to any analysis on recording data from grating and random dot experiments, we first identified the saturated channels. In this case, from the BISC 10-bit ADC range of [0, 1024), the range [8, 1016) was chosen to represent the non-saturated interval. For a given channel, the response in one trial was considered non-saturated when fewer than 0.1% of recorded signals was in saturated region, and a channel was considered as non-saturated when it is saturated in fewer than 1% of trials. An intersection of non-saturated trials of each non-saturated channel was defined as the non-saturated trials for the experiment session (about 300 trials in each grating session, 400 trials in each random dot session). All analyses were performed on non-saturated channels and trials.

We compared Fourier spectrum during fixation period and grating stimulus period for each non-saturated channel (**Fig. S11**). For frequencies evenly sampled in the range from 0 to 200 Hz, we computed the fraction of frequencies at which the channel responses in two periods differ by more than one standard deviation measured across trials. All channels with a fraction greater than 0.1 are labeled as responsive channels. Only responsive channels are used to compute the average spectrum in Fig. 5C. These channels are located at the V1 area of cortex.

#### Wavelet transformation for band-pass filtering

We applied Morlet wavelet transformation on the raw response from one channel to get time-varying band-passed signals. The complex wavelet is implemented by scipy.signal.morlet with scaling factor s=0.5. An example of a wavelet with central frequency of 64 Hz is shown in **Fig. S12**. The absolute value after wavelet transformation is defined as the signal for the corresponding frequency band.

#### Response scaling

Since the gains of each channel are not necessarily identical, we shifted and scaled band-passed responses before performing any tuning analysis, i.e. grating-triggered-average or dot-triggered-average. For a given channel, we first computed the mean response during the [-300, 0] ms window relative to stimulus onset for every trial. We define the average and standard deviation over trials as the baseline response and scaling unit, respectively. Band-passed responses were converted by the corresponding affine transformation.

#### Removing orientation-untuned component

The raw grating-triggered-average result reveals an orientation-independent component that mostly encodes the grating switching every 50 ms. The same example channel as in Fig. 5I is shown in **Fig. S13**. We computed the average over all orientations and all cycles and termed the residual as orientation tuned component (Fig. 5I).

#### Orientation selectivity index

Proper scaling of band-passed responses enables us to perform comparisons between channels. For each channel, we computed the temporal average during the interval from 88 to 112 ms after each grating onset conditioned on the grating orientation, denoting the result as 𝑓(𝜃). The orientation selectivity index is defined as difference between maximum and minimum value:

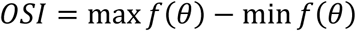

#### Orientation decoding

The orientation decoder was built on raw channel responses without any wavelet transformation. We first preprocessed the data by performing principal component analysis (PCA) to remove some correlations. The first 96 principal components (PCs), which capture 93.4% of the total variance, were kept and scaled according to the PC variance to serve as the input to the decoder. The decoder took the responses from these 96 PCs during the interval from 0 to 200 ms after a grating onset and binned them into 400 bins (0.5 ms each). The decoder backbone was a three-layer 1D convolution neural network with the last layer outputting a distribution between 0° and 180° represented by logits in a given number of bins. When training the decoder as a classifier, the cross entropy was used as the loss objective. We compared the mutual information between decoder prediction and input for different bin sizes used in discretization (**Fig. S14**) and observed little change when bin size is no greater than 15°. When training the decoder as a regressor, the Euclidean distance between the circular mean and the target orientation in the complex space was used as the loss objective. A small regularization coefficient (0.02) for distribution entropy was used in training the regressor to avoid non-smooth prediction. Regressor decoder performance is shown in **Fig. S15**, similar to Fig. 5L**-M**.

#### Two-component analysis of raw receptive field

In addition to dot-triggered-average of responses filtered by wavelet, we also computed dot-triggered-average of unfiltered responses (Videos S3 to S8). We observed structured spatial-temporal receptive field 𝑅𝐹(𝑥, 𝑦, 𝜏) on more channels; many of them locate on V4 and are not picked up by RF analysis using filtered response. We found that these RFs can be roughly approximated by two components (**Fig. S16**). Each component has a Gaussian-like spatial profile and the temporal profiles differ from each other:

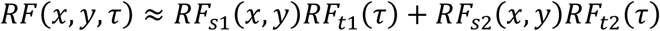

While spatial profiles 𝑅𝐹_𝑠1_(𝑥, 𝑦) and 𝑅𝐹_𝑠2_(𝑥, 𝑦) can be different for different channels, we assume 𝑅𝐹_𝑡1_(𝜏) and 𝑅𝐹_𝑡2_(𝜏) are shared by all channels. We computed the retinotopic map for both components separately. Component 1 was smaller in size, closer to the fixation point and shows up earlier than Component 2.

#### Identification of traveling waves

The very densely spatially sampled BISC recording channels make it an ideal candidate for detecting rich spatiotemporal neural activity such as traveling waves. We analyzed recordings from the visual cortex of the adult macaque monkey during the dot mapping experiment as described before. Traveling waves are brain oscillations that show individual cycles progressively propagating across the cortex in specific directions. We used a localized circular-linear regression approach to identify traveling waves in BISC recordings. In this approach, we assume that the relative phases of the BISC channels exhibit a linear relationship with channel locations *locally*^62,93,94^. This local circular-linear fitting of can reveal traveling waves that move at different directions across different areas of the electrode array, thus identifying complex patterns of traveling waves in addition to planar traveling waves^62^.

To identify these patterns, we first filtered the BISC signals in the gamma frequency band (30-90 Hz) by applying a 4^th^-order Butterworth filter, similar to analysis described in previous sections. We applied the Hilbert transform on each channel’s filtered signal to extract the instantaneous phase. To identify traveling waves, we used a series of two-dimensional (2-D) localized circular– linear regression to model the direction of wave propagation in a local cluster of channels in the 80-μm neighborhood of each BISC channel. The local regression determines the direction of local wave propagation in the cluster surrounding each channel, measuring whether the local phase pattern varies linearly with the channel’s coordinates in 2-D. Here, let 𝑥_𝑖_ and 𝑦_𝑖_ represent the 2-D coordinates and θ_𝑖_ the instantaneous phase of the *i-*th channel in a cluster. We used a 2-D circular-linear model

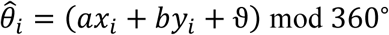

where *θ̂*_𝑖_ is the predicted phase, 𝑎 and 𝑏 are the phase slopes corresponding to the rate of phase change (or spatial frequencies) in each dimension, and ϑ is the phase offset. We converted this model to polar coordinates to simplify fitting. Let 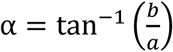 denote the angle of wave propagation and 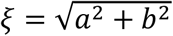 denote the spatial frequency. We fitted 𝛼 and 𝜉 to the distribution of phases at each time point by conducting a grid search over 𝛼 ∈ [0^∘^, 360^∘^] and ξ ∈ [0,6.2]. Note that 𝜉 = 6.2 corresponds to the spatial Nyquist frequency of 6.2°/μm, corresponding to the 29 μm spacing between neighboring channels.

We carried out a grid search in increments of 5° and 0.5°/μm for *α* and 𝜉, respectively. The model parameters (𝑎 = ξ cos(α) and 𝑏 = ξ 𝑠𝑖𝑛(α)) for each time point are fitted to most closely match the phase observed at each channel in the cluster. We computed the goodness of fit as the mean vector length of the residuals between the predicted (𝜃^_𝑖_) and actual (𝜃_𝑖_) phases^95^,

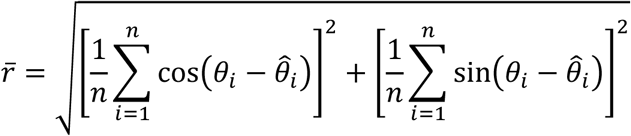

where *n* is the number of channels in each cluster. The selected values of 𝛼 and ξ are chosen to maximize 𝑟̅. This procedure is repeated for each cluster. To measure the statistical reliability of each fitted traveling wave, we examined the phase variance that was explained by the best fitting model. To do this, we computed the circular correlation, 𝜌_𝑐𝑐_, between the predicted (𝜃^_𝑖_) and actual (𝜃_𝑖_) phases at each channel:

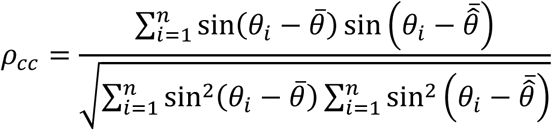

where bar denotes averaging across channels. We also refer to 𝜌_𝑐𝑐_ as the wave strength^93^.

#### Dimensionality reduction analysis for wave patterns

To visualize how the spatial patterns of traveling wave propagation differ between multiple viewed dot locations, we used UMAP^96,97^ to embed both data features and class labels into a low-dimensional manifold. Specifically, we set n_neighbors = 15, min_dist = 0.1, and metric = euclidean in the UMAP toolbox^97^; the supervision was introduced via a cross-entropy term that encourages separation among stimulus classes. These hyperparameters were selected after grid search on the validation dataset to balance local neighborhood preservation against global cluster separability. We applied this approach to downsampled datasets to test the contribution of high-density recordings; here wave features were recalculated for each subset to prevent information leakage from the full-resolution grid. After UMAP projection, each trial’s wave features were embedded in 2D, and stimulus class separation was quantified using the silhouette score:

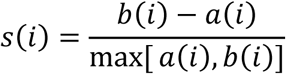

where 𝑎(𝑖) is the average distance of the i-th sample to members of its own cluster, and 𝑏(𝑖) is the lowest average distance of the i-th sample to any other cluster. Higher mean silhouette scores indicate greater separability. Scores were averaged over 300 random subsampling seeds per resolution, with variability assessed via standard error.

#### Dot location decoder: hybrid CNN-Transformer-based neural network model

To decode dot locations from BISC-recorded traveling-waves, we developed a hybrid CNN-Transformer architecture that predicts the viewed dot location for each traveling wave while accounting for spatial and temporal dependencies. To decode the spatial location of the stimulus from traveling-waves, we first passed the pattern of traveling wave propagation for each dot into a CNN module. The CNN comprised three sequential convolutional layers with filters of size 3 × 3, each followed by ReLU activation and batch normalization (momentum = 0.1). We employed a max-pool layer with a stride of 2 after the first and second convolutional layers, reducing the dimensionality of the spatial representation. The final feature map was flattened, yielding a 128-dimensional vector that served as input to the subsequent Transformer.

Across dot locations, the flattened CNN outputs were reorganized into a sequence of feature vectors corresponding to consecutive time windows (each ∼50 ms), which were then passed into a Transformer with two encoder layers. Each layer used multi-head self-attention (4 heads; embedding dimension = 128), followed by a position-wise feedforward layer (hidden dimension = 256). A dropout rate of 0.1 was applied to both attention weights and feedforward activations to prevent overfitting. The Transformer thus captured potential long-range temporal dependencies in wave propagation. We employed sinusoidal positional encodings^98^ to preserve the ordering of time windows.

We allocated 80% of the trials to training, 10% to validation (for hyperparameter tuning and early stopping), and 10% to hold-out testing. Network weights were optimized via the Adam optimizer (α = 0.001, β_1_ = 0.9, β_2_ = 0.999) with a batch size of 32. We trained for a maximum of 1000 epochs, applying early stopping if validation loss did not improve over 10 consecutive epochs.

#### Loss function and multi-task decoding

To facilitate training of the network on the dynamic stimulus detection task we used a common hybrid approach for multi-task learning^99^. The final output layer branches into two heads: (1) a classification head, predicting discrete stimulus locations (e.g., four classes for four possible stimulus sites), and (2) a regression head, estimating continuous coordinates in the stimulation array. The overall loss was:

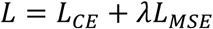

where 𝐿_𝐶𝐸_is cross-entropy loss for classification, 𝐿_𝑀𝑆𝐸_is mean-squared error between predicted and actual coordinates, and 𝜆 was set to 0.5 to balance the two objectives (tuned using grid search). During the initial training phase, noise regularization was implemented by adding 5% additional data augmentation to the input, a method used for enhancing model generalization in the context of time series prediction^100^ and image processing^101^.

Temporal shuffling control: To test the impact of temporal structure on decoding, we randomly permuted the order of presented dot time windows within trials while preserving spatial features. The effect of this shuffling was to specifically disrupt temporal correlations in neural signals between consecutive dot presentations. We repeated this procedure across all test trials 10 times to assess variability and ensure reliable performance estimates. A two-sided paired t-test (p < 0.01) was used to evaluate differences between the original and shuffled conditions. The number of iterations was chosen based on empirical stability in decoding performance across runs.

#### Deep neural network models trained on natural images

The deep learning based neural predictive model consisted of two main parts: A pretrained core that computes image embeddings (i.e. a shared feature map given an input image) and a readout that maps these features to the neuronal responses of a single recording channel. As a core, we selected ConvNext-v2-tiny ^69^, a recently published convolutional neural network model trained on ImageNet ^65^. We used the original neural network weights with fine-tuning in a two-step training process. As a readout, we fit an attention readout, described in detail elsewhere ^70^, to transform the core feature map into a scalar neural response for each recording channel.

In brief, the readout first adds a fixed positional embedding to the image embeddings with subsequent normalization through LayerNorm. Then, key and value embeddings are extracted by position-wise linear projections, both of which have parameters shared across all recording channels. Then, for each channel, a learned query vector is compared with each position’s key embedding using scaled dot-product attention. The result is a spatially normalized attention map that indicates the most important feature locations for a recording channel, given an input image. We then use this attention map to compute a weighted sum of the value embeddings, which results in a single feature vector for each neuron. Finally, a neuron-specific affine projection with exponential linear unit (ELU) non-linearity gives rise to the scalar predicted neuronal activity. We refer to this architecture as a cross-attention readout because the key and value embeddings are obtained from the image, whereas the query is learned for each recording channel separately.

The model is trained by minimizing the Poisson loss between recorded and predicted neuronal activity, identical to the procedures described in ^66^. Here, we first freeze the weights of the core and train the readout for 30 epochs. Then, we reduce the initial learning from 0.001 to 0.0001 and optimize the weights of the core and readout using the AdamW optimizer for a total of 200 epochs.

#### Explainable variance

As a measure of response reliability, we estimated the fraction of the stimulus-driven variability compared to the overall response variability. More specifically, we computed the ratio of each channel’s total variance minus the variance of the observation noise, over the total variance. To estimate the variance of the observation noise, we averaged the variance of responses across image repeats for all of the 75 repeated natural image test stimuli.

#### Generation of MEIs

We used the trained model to synthesize maximally exciting input images (MEIs) for each channel using regularized gradient ascent. Starting out with a randomly initialized Gaussian white noise image, we showed the image to the model and computed the gradients of a single target channel with respect to the image. To avoid high frequency artifacts, after each iteration, we applied Gaussian blur with an SD of three pixels to smoothen the image. Additionally, we constrained the entire image to have a fixed-energy budget, which we implemented as a maximum L2 norm of the z-scored image, calculated across all pixel intensities. We chose a total L2 norm of the MEI of 20, such that the resulting MEIs had minimal and maximal values similar to those found in our training natural image distribution. Additionally, we made sure that the MEI could not contain values outside of the eight-bit pixel range by clipping the MEI outside of the bounds that correspond to 0 or 255 pixel-intensity. We used stochastic gradient descent (SGD) optimized with learning rate of five and ran each optimization for 500 iterations, without early stopping.

## Supplementary Information for

### Supplementary Discussion S1. BISC implant design

As illustrated in **Fig. S1**, the implant circuitry consists of four primary top-level modules, each of which can be implemented and verified independently. These modules are the analog front-end (AFE), the controller, the wireless transceiver, and the wireless power transfer (WPT). The AFE is directly interfaced with the electrodes. It is responsible for sensing the neural signals on the electrodes, which are then amplified locally in the in-pixel amplifiers. The outputs from these amplifiers are transferred to a programmable gain amplifier (PGA) through column switches and subsequently digitized by a 10-bit successive-approximation-register (SAR) analog-to-digital converter (ADC). Additionally, the AFE is designed to drive stimulation currents through the electrodes from global current sources, utilizing the column and in-pixel switches. The controller plays a central role in managing the system. It sets all static configurations, including biasing and gain control for the AFE. Furthermore, it controls all dynamic switching signals into the pixel array and the column switches. The controller also receives digital samples from the ADC, packaging the data for transmission via the wireless transceiver, as well as receiving, depacketizing, and decoding serialized data from the wireless transceiver. While not explicitly shown in the diagram, the controller also controls the activation and deactivation of the voltage regulators for the AFE. The wireless transceiver module is responsible for the receiving and transmitting data through the UWB antenna. It also supplies the controller with a clock signal, which is generated in a phase-locked loop (PLL) using a reference clock from the WPT module. Lastly, the WPT module is designed to inductively receive RF power through a coil. This RF power is initially rectified to 2V, and then further regulated by three 1.5 V regulators dedicated to the AFE, the controller, and the transceiver. Additionally, the rectified 2V is converted to −2V, and then regulated to −1.5V specifically for the AFE.

### Design of the analog front-end

The analog front-end (AFE) is an analog mixed-signal module that serves as the interface between the electrodes and the controller. Within this module, various circuit blocks are implemented along two primary signal paths. The first is the recording path, which originates at the electrodes and progresses through an in-pixel amplifier (**Fig. S17A**), column switches, a programmable gain amplifier, and a 10-bit ADC, terminating at the controller. The design of the recording path was inspired by the architecture of an active-pixel sensor (APS) ^102^ photosensor ^103^, which utilizes a charge amplifier readout mechanism. The APS architecture was used to avoid long high-impedance routings from electrodes, which are susceptible to electromagnetic interference from the 13.56 MHz wireless powering frequency. Such interference, unless carefully filtered or mitigated, can lead to large offsets in the amplifiers due to circuit nonlinearity ^104,105^. To minimize the potential interference, the routings from the electrodes to the inputs of the in-pixel amplifiers are restricted to within 20 μm. Furthermore, the APS architecture relaxes the performance requirement of the shared back-end circuitry, allowing implementation of PGA and ADC with smaller area and lower power consumption.

The pixels and column switches in the AFE utilize a dual rail ±1.5 V power supply. This power supply configuration allows maximum safety because 0-V bias voltage can be applied on the electrodes and there is no DC voltage difference between the working, reference, counter electrodes, and the substrate. Potential current leakages are eliminated from all conductive parts of the implant. To support this, the pixels and column switches are implemented in a −1.5 V deep P-well (DPW), isolated from the rest of the circuits by an N–type buried layer (NBL). Each pixel is identical and occupies an area of 53 × 58 μm^2^ (**Fig. S17B**). In every pixel there are four 14 × 14 μm^2^ electrode pads on the redistribution layer (RDL). These electrode pads connect to the underlying circuits by redistribution via (RV).

For every pixel, the recording process utilizes a neural amplifier and multiple control switches. Given the high-density requirements of the APS architecture, the area available for the neural amplifier layout is significantly restricted. On the contrary, optimal noise and mismatch performance requires us to bias the metal-oxide-semiconductor field-effect transistors (MOSFET) in the weak inversion region, which demands a large area ^106^. To address this challenge, we made four design decisions aimed at maximizing the area efficiency for recording circuitry. Firstly, we implemented the stimulation current sources as a global current generator outside the pixel array, incorporating only MOSFET switches and combinational logics within the pixels to realize stimulation. Additionally, the control signals for the stimulation circuits undergo global level shifting outside the pixel array. Secondly, for the neural amplifier, we adopted a DC-coupled architecture which eliminates the need for a dedicated large DC blocking capacitor by utilizing the double-layer capacitance formed between the electrodes and the electrolyte. Thirdly, we implemented a multiplexing technique for recording from the four electrodes in each pixel to connect to the neural amplifier, significantly reducing the number of amplifiers ^107^. Lastly, we implemented a differential integrator as the neural amplifier (**fig. S15B**) ^42^ leveraging the offset memorization principle ^108^ and boxcar averaging using integrating, sampling, and reset operations (**fig. S15E**) to provide anti-aliasing filtering without the need for bulky components. This approach utilizes chopping operation ^108^ that effectively cancels out the DC offset and 1/𝑓 noise inherent in the neural amplifier. The noise transfer function 𝐻_𝑁_ of the neural amplifier is given by:

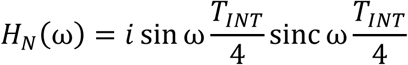

where 𝑇_𝐼𝑁𝑇_is the integration time of the neural amplifier. The sine function in the noise transfer function attenuates low frequency noise (**Fig. S17F**). Consequently, this approach achieves low noise and offset without relying on overly large input transistors. Common-mode feedback (CMFB) of the differential integrator can also be efficiently implemented using MOSFET switches and two metal-insulator-metal (MIM) capacitors by taking advantage of the reset phase ^109^. Moreover, the open-loop configuration of this design eliminates the need for large feedback components while still maintaining a well-defined gain and low-pass corner. The neural amplifier can be considered as a trans-capacitance amplifier and its voltage-to-charge gain 𝐴_𝑐_ can be calculated by:

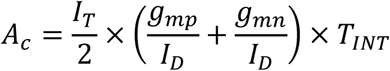

where 𝐼_𝑇_ is the tail current of the neural amplifier, 𝐼_𝐷_, 𝑔_𝑚𝑝_ and 𝑔_𝑚𝑛_ are the drain current and transconductances of the input PMOS and NMOS transistors. The 0-V biasing at the input of the neural amplifier is implemented using NMOS transistors configured as tunable pseudo-resistors. The gate voltage of these NMOS transistors is globally generated from a W-2W voltage digital-to-analog converter (DAC) ^110^. These pseudo-resistors in conjunction with the electrode capacitance allow the neural amplifier to reject up-to 25 mV DC-offsets in the electrodes and provide a tunable high-pass corner. The 3-dB low pass corner ω_lp_ of the neural amplifier is fixed and can be calculated as:

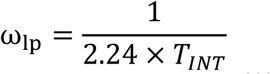

where coefficient 2.24 is the result of boxcar averaging ^111^ whose transfer function is a sinc function (**Fig. S15F**). In this design, 𝐼_𝑇_ is globally programmable from 0.625 to 4.375 µA, 𝐼_𝐷_ is half of 𝐼*_T_* 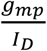 is 22 V^−1^, 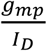 is 26 V^−1^, and 𝑇_𝐼𝑁𝑇_ is 29.38 µs. Since the input transistors are biased in the weak inversion region, their 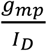 values stay nearly constant with differential values of 𝐼_𝐷_. Custom designed 3-V logical cells (**Fig. S17D**) are used to generate the control signals to the neural amplifier.

The programmable gain amplifier (PGA) (**Fig. S18**) consists of a charge amplifier stage (**Fig. S18A**) followed by a programmable gain stage (**Fig. S18B**). The operation of the charge amplifier involves an operational transconductance amplifier (OTA) switching between two sets of feedback capacitors. When one set of the metal-insulator-metal (MIM) capacitor is used in amplification, the other set is reset to ground and the common-mode voltage (VCM). This operation allows almost an entire clock cycle to be used for OTA settling, relaxing the bandwidth requirement of the OTA and reducing power consumption. The charge amplifier stage significantly reduces the nonlinearity of the PGA compared to a voltage amplifier because the metal-oxide-semiconductor (MOS) sampling capacitors in the pixels are highly nonlinear. Direct amplification of the voltage output from the pixels would limit the overall effective number of bits (ENOB) to seven bits. With the MIM capacitors for charge transferring, the system can have a linearity over 10 bits. The programmable gain stage utilizes a modified predictive switched-capacitor amplifier ^112^. The voltage gain of this stage is programmable from 1.5 to 5.0. The total voltage gain of the system 𝐴_𝑠𝑦𝑠_ is given by:

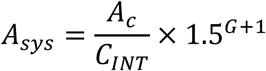

where 𝐶_𝐼𝑁𝑇_ is 5.44 pF and 𝐺 is a two-bit programmable value from 0 to 3.

Two 10-bit 8.67-MS/s SAR ADC ^113^ are used to digitize the outputs from the PGA in an interleaved operation. The two ADCs effectively work as a single ADC, but the interleaving relaxes the conversion time of the ADC and the settling time of the PGA. This reduces the power consumption of the ADC and PGA.

The recording process in pixel at row ‘r’, column ‘c’ is controlled by four row lines (PWR_R_ [r], CDS_R_ [r], REC_R_ [r], RST_R_ [r]) and five column lines (PWR_C_ [c], RST_C_[c], REC_C_[c], SEL0_C_[c], SEL1_C_[c]) as shown in **Fig. S17D**. These control lines operate the recording circuit in six phases. The first phase is power-on, and in this phase PWR_R_ and PWR_C_ are activated, turning on the bias currents in the neural amplifier. The second and third phases are the amplification phase in which auto-zeroing happens. They are differentiated by the toggling of the MOSFET chopper switches. When the CDS_R_ and RST_C_ are activated, a clock divde-by-2 circuit toggles its output, consequently, flips the polarity of the chopper. The fourth phase is readout, where REC_R_ and REC_C_ are activated, turning on the output MOSFET switches in the neural amplifier and column switches. The charges integrated by the neural amplifier are transferred to the programmable gain amplifier via the switches, and subsequently digitized by the ADC. The fifth phase is reset, where RST_R_ and RST_C_ are activated, resetting the charges on the integration capacitors through the reset MOSFET switches. The switched-capacitor common-mode feedback (CMFB) also refreshes its biasing point at this phase. Additionally, the electrode address for multiplexing will be updated from two column lines SEL0_C_[c], SEL1_C_[c]) and saved in two D-latches. If recording continues, after the reset phase the neural amplifier will go into the amplification phase. Otherwise, it goes into the last phase which is power-off. In the power-off phase all column and row lines are deactivated, and electrodes are electrically disconnected from the neural amplifier. The recording control lines are locally level shifted to ±1.5 V logic level in each pixel, and the logic circuits are custom designed to work under ±1.5 V.

For stimulation, there are 16,384 stimulation channels (one per pixel) and a bi-phasic current pulse can be distributed across a programmable set of pixels (all pixels generate the same temporal current profile) acting as a “macroelectrode” in either monopolar or bipolar configuration (see **Methods**, **Fig. S19**). The temporal profile of the bi-phasic simulation can be configured to be either cathodic-first or anodic-first, followed by a passive charge balancing phase. Stimulating pixel address can be quickly reprogrammed within 100 μs, which offers opportunities to inject high density spatiotemporal information into the brain.

The stimulation currents are generated globally from a stimulation current generator (**Fig. S19**), consisting of two regulated-cascode current sources. A stimulation-current digital-to-analog converter (DAC) provides programmable biasing of the current sources. The reference current is generated from a low-voltage bandgap reference circuit ^114^ via a master current DAC. The amplitude of the current output 𝐼_𝑜𝑢𝑡_ of the stimulation current generator in µA is:

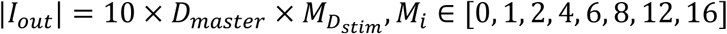

where 𝐷_𝑚𝑎𝑠𝑡𝑒𝑟_ and 𝐷_𝑠𝑡𝑖𝑚_ are three-bit values from 0 to 7.

The stimulation process in each pixel is controlled by a column line (PROG[c]) and three row lines (SRCS[r], SINK[r], CBAL[r]) routed to the pixels from the controller. These digital control lines are level-shifted to ±1.5 V logic levels before they enter the pixel array. Two current column lines (ANOD[c], CATD[c]) are routed into the pixels from a stimulation current generator through the column switches. Activation of the PROG line and one of the row lines by the controller initiates the stimulation sequence: the MOSFET switches connect the four electrode pads in the respective pixel to the ANOD for anodic current, the CATD for cathodic current, and ground for charge balancing.

To optimize power efficiency in the implant, both the recording and stimulation circuits are designed with the capability to be individually shut down by deactivating their respective bias currents. The implant is operated in a manner that ensures either the recording or the stimulation circuits are powered on at any given time, but not both simultaneously. Switching between recording and stimulation introduces a latency of approximately 1 ms to start up the circuits.

#### Design of the wireless transceiver

The wireless transceiver consists of ultra-wide-band (UWB) antenna and transceiver circuits. On-off keying (OOK) is used for data modulation. The wireless transceiver is designed for low power, single user, and short-range communication.

The UWB antenna, having a differential-fed slot design ^115^, was modified to work under brain environment (**Fig. S20A**). To achieve this, custom multi-layer human head models were built that incorporate various tissue characteristics, including tissue thickness ^116^ and dielectric constants ^117^. We performed electromagnetic simulation (Momentum, Keysight Technologies and Ansys HFSS, Ansys) to determine the parameters of the UWB antenna with the custom brain model. Given that both the UWB antenna and the transceiver circuits are custom designs for the implant, they were co-designed for optimal performance. The input impedance at 4 GHz was designed to be 50 + j16 Ω. The path loss is simulated to be 50 dB (Ansys HFSS, Ansys) for a total tissue thickness of 2 cm between the designed antenna and an ideal dipole antenna (**Fig. S20A**). A 700-MHz bandwidth is achieved from 3.6 to 4.3 GHz.

The UWB antenna is shared by the transmitter and receiver circuits using two sets of NMOS switches. To minimize insertion loss at the working frequencies (3.6 to 4.3 GHz), a floating-body technique ^118^ is used which takes advantage of the deep p-well (DPW) in the CMOS technology. Resistors are used in-series with their gates to reduce RF leakage to the DC control lines.

Duty-cycling is a proven technique to reduce power consumption of impulse radio UWB (IR-UWB) transceivers ^119,120^. The transmitter employs this technique by generating short RF pulses using a duty cycled LC complementary oscillator. The LC oscillator is tuned to resonate with the UWB antenna at 4 GHz without a local frequency reference. A digital pulse generator generates two short baseband pulses with 1.3 and 0.3 ns pulse width from an enable signal (**Fig. S20B**). When transmitting a data “1”, the LC oscillator’s tail current is turned on for 1.3 ns. To speed up the start-up of the LC oscillator, the KICK_N MOSFET switch is turned on for the first 0.3 ns. Conversely, for a faster shut-off, the shunt MOSFET switch is turned on while the RF switches are turned off. The LC oscillator delivers 1 V peak-to-peak differential signal into the UWB antenna when it is turned on. The designed transmitter transmits at maximum 108.48 × 10^6^ pulses per second and consumes 39 pJ for each pulse transmitted.

The receiver has a non-coherent energy detector architecture for its low implementation complexity ^121,122^. Consider a noise floor (N_0_) at body temperature of −173.6 dBm, a minimal signal-to-noise ratio (SNR_REF_) of 17 dB for a bit-error rate (BER) less than 1 × 10^-10^ ^123^, path loss (PL) of 50 dB, receiver noise figure (NF) of 5 dB and data rate (DR) of 54.24 Mbps, the minimal required transmit power (P_TX_) of the headstage is given by:

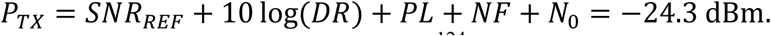

According to the FCC regulations on the UWB band ^124^, a maximum equivalent isotropic radiated power (EIRP) of −41.3 dBm/MHz can be transmitted in free air and measured over a 1ms time window. The maximum power (P_TX_MAX_) allowed to transmit for a 500-MHz bandwidth is given by:

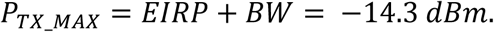

We also comply with the average power limit as P_TX_ is less than P_TX_MAX_. Also, the peak power limit of 0 dBm is satisfied as we operate in the high-data-rate regime ^122^. Since there is a −14.3 + 21.3 = 10 𝑑𝐵 link margin, we find that the design is robust against manufacturing variations and variations in the distance between the implant and headstage.

The UWB receiver features an RF architecture similar to that described in ^125^. The signal processing begins with amplifying the input signal from the UWB antenna by a low-noise amplifier (LNA), which comprises an input stage followed by two RF amplifier (RF AMP) stages (**Fig. S20C**) that provide 57 dB of voltage gain with in-band noise figure less than 5 dB. A self-mixing mixer based on double-balanced Gilbert-cell detects the signal’s envelope (**Fig. S20D**), which is low-pass filtered by a baseband amplifier (BB AMP) (**Fig. S20E**). Finally, a one-bit digitizer (1-b ADC) comprised of a sampling and subtraction stage (**Fig. S20F**) followed by a StrongArm comparator^126^ determines the received bit (“0” or “1”) based on a predefined voltage threshold.

A phase-locked loop (PLL) modified from ^127^ is used to generate a 208.96 MHz clock from the 13.56 MHz powering frequency, which is divided down to generate all the clocks used in the implant. A 4-stage differential ring oscillator is used which allows timing control at the resolution of 0.567 ns. A digital controller inside the wireless transceiver controls the start-up of the PLL and timing generation.

The timing diagram of the UWB transceiver is shown in **Fig. S20G**; the LNA is powered on 4.61 ns and the mixer and based-band low-pass amplifier are powered on 6 ns for every 18.44 ns. The entire transceiver consumes 12.2 mW when transmitting at 50% of “1”s.

#### Design of the BISC implant controller

The digital controller on the BISC implant is responsible for managing the wireless communication, device configuration, and dynamic data acquisition. It operated from the 1.5-V power domain on the chip and consumes a peak power of 10 mW during recording. The controller consists of a main decoder that translates the instructions received by the wireless transceiver into actions on the device. The decoder can decode only one instruction at a time and while an instruction is being executed, other instructions cannot be accepted by the controller.

The decoder interfaces to two specialized control blocks:

- *<u>AFE control.</u>* AFE control sets the static configuration for recording and stimulation functions and outputs dynamic control signals that trigger the switching activity in the AFE.
- *<u>Communications control.</u>* This block receives data samples from the AFE and packetize them for transmission via the wireless transceiver. During a recording, every new sample of data is immediately added to the next packet. When a packet is full, it is passed to be streamed through the wireless transceiver, while the preparation of the next packet begins. The communication control can also be configured to packetize and transmit other types of data regarding general information on the current state of the device. The packetizing follows the specialized communication protocol described in more detail in **upplementary Discussion S2**.

The instruction set architecture (ISA) of the controller consists of seven instructions (**Table S1**) that can be classified into three types: dynamic instructions, static instructions, and the query configuration instruction.

Dynamic instructions include recording, stimulation, power-on, and halt. Each of these instructions initiates an event that starts or stops an ongoing process. The recording instruction starts a process that updates the control lines going to the AFE which allow time-multiplexed recording from a designated subset of channels from the microelectrode array. The stimulation instruction initiates multi-phasic stimulation through a pre-configured subset of electrodes for a pre-configured amount of time. The power-on instruction powers on the pixel-level recording amplifiers, which is required prior to recording the electrodes before recording. The halt instruction stops an ongoing recording.

Static instructions include configuration and programming. The configuration instruction can program up to 64 bits of configuration in a given instruction. These configuration registers, which number 122 bits in total, control the functioning of finite state machines (FSMs) in the controller or are distributed to other functional blocks on the BISC implant, such as the WPT, the AFE, and the UWB transceiver in order to set longer-term states that are infrequently updated. The programming instruction sets the address of electrodes used for stimulation and their initial polarity, along with additional configuration information that are stored in the controller’s registers. When stimulation is initiated, this information is used to gate specific control signals sent to the AFE.

The query configuration instruction allows one to request access to the contents of the configuration registers.

#### Design of the WPT circuit

The WPT circuit receives power from the relay station by means of 13.56 MHz inductive coupling, converting the received power to regulated ±1.5V supplies (**Fig. S21**). RF inductive power received by the power coil (**Fig. S1C**) is first converted to a DC voltage using an active rectifier. This DC voltage, which can be anywhere between 2 and 3.3V to support full functionality of the device, is further converted to both +1.5V and −1.5V supplies. To generate −1.5V, the rectifier output voltage goes through an additional stage (**Fig. S21B**) prior to the regulation stage.

The output voltage level of the rectifier is a function of the amount of power received by the BISC implant and the amount of power consumed by it. In a practical setting, both factors are dynamically changing. For example, the former depends on the coil-to-coil alignment between the device and the relay station headstage, and the latter depends on the device’s mode of operation. For thermal and specific absorption rate (SAR) characterization, we assumed that our device is in a recording state, which is the most power-hungry mode of operation.

Ideally, the power delivered to the device should be sufficient but no more than what is necessary to support full operation, since excess power received becomes converted to heat. This means that under optimum operation, the rectifier output voltage is stabilized to near 2V. To achieve this, we periodically read out the rectifier output in a three-bit digitized format (0: 2V, 1: 2.1V, …, 7: 2.7V) using the query configuration instruction to the BISC implant and adjust the magnitude of transmitted power from the headstage.

Coil-to-coil link efficiency is a key metric in inductive WPT design. A more efficient link not only allows a longer operating distance, but also helps to keep the power radiated by the headstage to stay below the SAR safety exposure limit (2 W/kg) set by the FCC ^47^. We characterized our link efficiency by measuring S-parameters of the two coils with chicken breast as a tissue phantom (Fig. 20A). We assumed an ideal conjugate-matched impedance on the transmitting side (Port 1) and a 75 Ω load on the receiving side (Port 2) which is the periodic steady-state (PSS) simulated linear-load equivalent of the time-varying rectifier input impedance.

Temperature of the chip was measured with a thermal imaging camera (FLIR E5-XT, Teledyne FLIR) under different power delivery conditions (Fig. 22B). Measurements were taken in an ambient setup with no active air flow at 25 °C, after the device operating in recording mode reached thermal equilibrium. For this measurement, we used a sample that is fully passivated on both sides. In recording mode, our device consumes 63.5 mW, including the coil loss. SAR simulation (Ansys^®^ Electronics Desktop, Ansys) assuming 1.5-cm implant depth using a six-layer brain model ^128^ demonstrates that our system can operate below the 2 W/kg limit set by the FCC for wireless devices used against the body operating below 6 GHz (Fig. 22D).

### Supplementary Discussion S2. BISC wireless link protocol and software stack

The BISC wireless link protocol consists of both uplink and downlink protocols. The downlink protocol packetizes the information transmitted from the relay station to the implant, while the uplink protocol packetizes the information sent from the implant to the relay station. Both types of packets hold a total of 125 bits. Packets from the relay station are received by the BISC implant bit-by-bit and stored in a shift buffer that matches the size of one packet.

Encoder modules in the BISC implant and in the relay station prepare the packets before transmission. The packets contain the information to be transmitted, along with several fixed-size segments of bits that are used for communication synchronization which we denote as “sync bits” occupying unique positions inside each packet. In addition, the encoder adds eight bits of error correction code (ECC) to implement single error correction, double error detection (SEC-DED), computed from the information in the packet and the sync bits. Decoder modules in the implant and the relay station are in charge of probing the shift buffer at each clock cycle, searching for the sync bits, and identifying when a full packet is present on the device. When a packet is identified, it is copied from the shift register and forwarded for further processing.

**Fig. S23** shows the packet structure for both the downlink and the uplink. In both packets, the parity bits hold the ECC code, and the preamble bits are unused. In the downlink packet, the OPCODE is a unique code that identifies the instruction (**Table S1**). The OPERAND bits include an address to a block in the controller and any other parameters that are needed to execute the command. In the uplink packet, the preamble bits are used to synchronize the order of recorded values during recording; otherwise they are unused. The DATA bits hold the information from the implant to be processed by the relay station.

**Fig. S24** presents the software stack in the BISC system and the hardware interface. The software stack is executed on the processing system (PS) of the relay station. The top layer of the stack is the RESTful API that implements an interface between an external client to the lower software layers in the stack. The commands sent to the relay station from an external client trigger Python applications that allocate memory through direct memory access (DMA) and make the preparations to intercept data from the implant. When the preparations are done, the applications use system calls to drivers, which access memory-mapped registers to configure the hardware in the programmable logic (PL) and retrieve the addresses for the allocated memories. The Python applications and drivers are implemented on top of a PYNQ ^129^ overlay embedded in the Linux operating system. PYNQ provides libraries that abstract the interaction with the hardware interface between the PS and the PL. Once the memory-mapped registers are configured with the information to execute high-level commands from the embedded API in the PL, the hardware in the PL executes the commands, prepares sequences of instructions to be sent to the implant, and then forwards them to be processed by the BISC controller. The PL communicates instructions to the implant through a designated module that interfaces with the transceiver on the headstage. The PL incorporates a data management module to intercept incoming data from the implant and store it using DMA in the memories allocated by the Python applications. The PL and the PS interact through interrupts that signal when the execution of the high-level commands is completed.

**Fig. S25** shows the graphical user interface (GUI) of the software used for recording.

### Supplementary Discussion S3. Bench-top *in vitro* characterization

BISC recording and stimulation (Fig. 2, **Fig. S26B-G)** were characterized from a fully wireless*, in vitro* measurement setup (**Fig. S26A**). A custom 3D printed mold was used to create two separate chambers filled with electrolyte: one over the pixel array and the other over the reference/counter electrodes. Silicone sealant (Kwik-Cast™, World Precision Instruments) was used to attach the device to the mold to prevent any leakage between the two chambers. Because of small device feature size, however, the sealant inevitably flowed over to a small portion of the array, encasing some electrodes (< 5%) at the edge of the array.

A sample processed with titanium nitride electrodes and front-side passivation was used for the measurement. Because of hydrophobicity of the polyimide used for front-side passivation, we needed to apply droplets of isopropyl alcohol over the electrodes as the “wetting layer” prior to filling the chambers with 1× phosphate-buffered saline (PBS) solution. For *in vivo* experiments, this step was not needed because EtO sterilization significantly reduced the hydrophobicity of the polyimide. Ag/AgCl pellet electrodes (EP1, World Precision Instrument) were used as the electrochemical interface between the 1× PBS and electrical cables.

For recording frequency response and pixel gain characterization (Fig. 2C**-E**, **Fig. S26B-E**), electrical cables were driven by a 400 µV_PP_ sinusoid (AWG3102C, Tektronix. In conjunction with −40 dB attenuator). For noise characterization (**Fig. 2B** and **F**), the two Ag/AgCl electrodes were shorted together. For both measurements, relay station headstage and the sample being measured were placed inside a Faraday cage. For stimulation characterization (**Fig. 2G** and **H**, **Fig. S26F** and **G**), electrical cables were connected to a low noise current amplifier (SR570, Stanford Research System) to measure current waveforms.

---

**Fig. S1.**
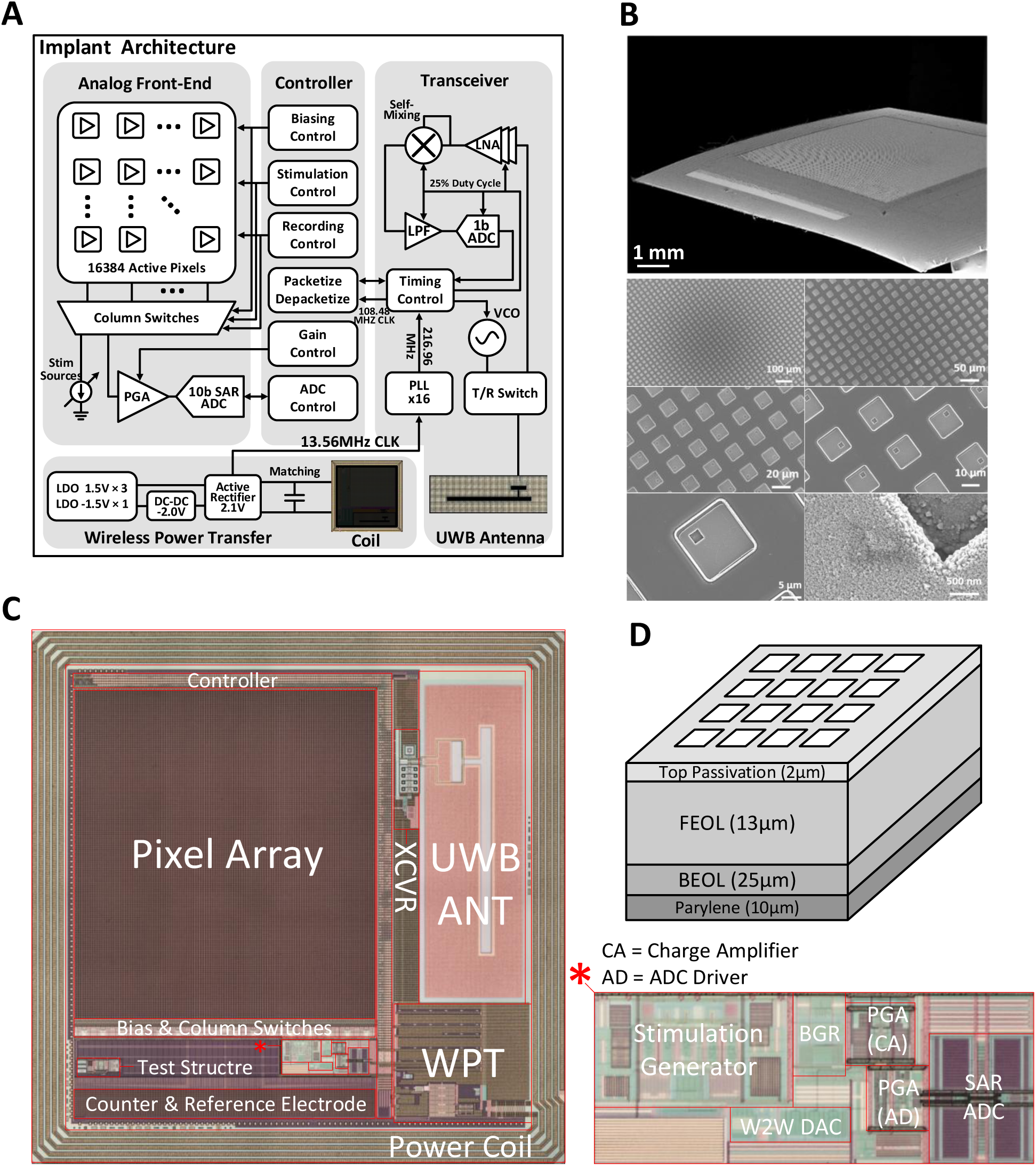
Architecture of the implant. (**A**) Functional diagram of the modules on the implant. (**B**) Cross-section and electrode SEM image. (**C**) Die photo showing functional components on the integrated circuit. (**D**) Stack-up of the implant.

**Fig. S2.**
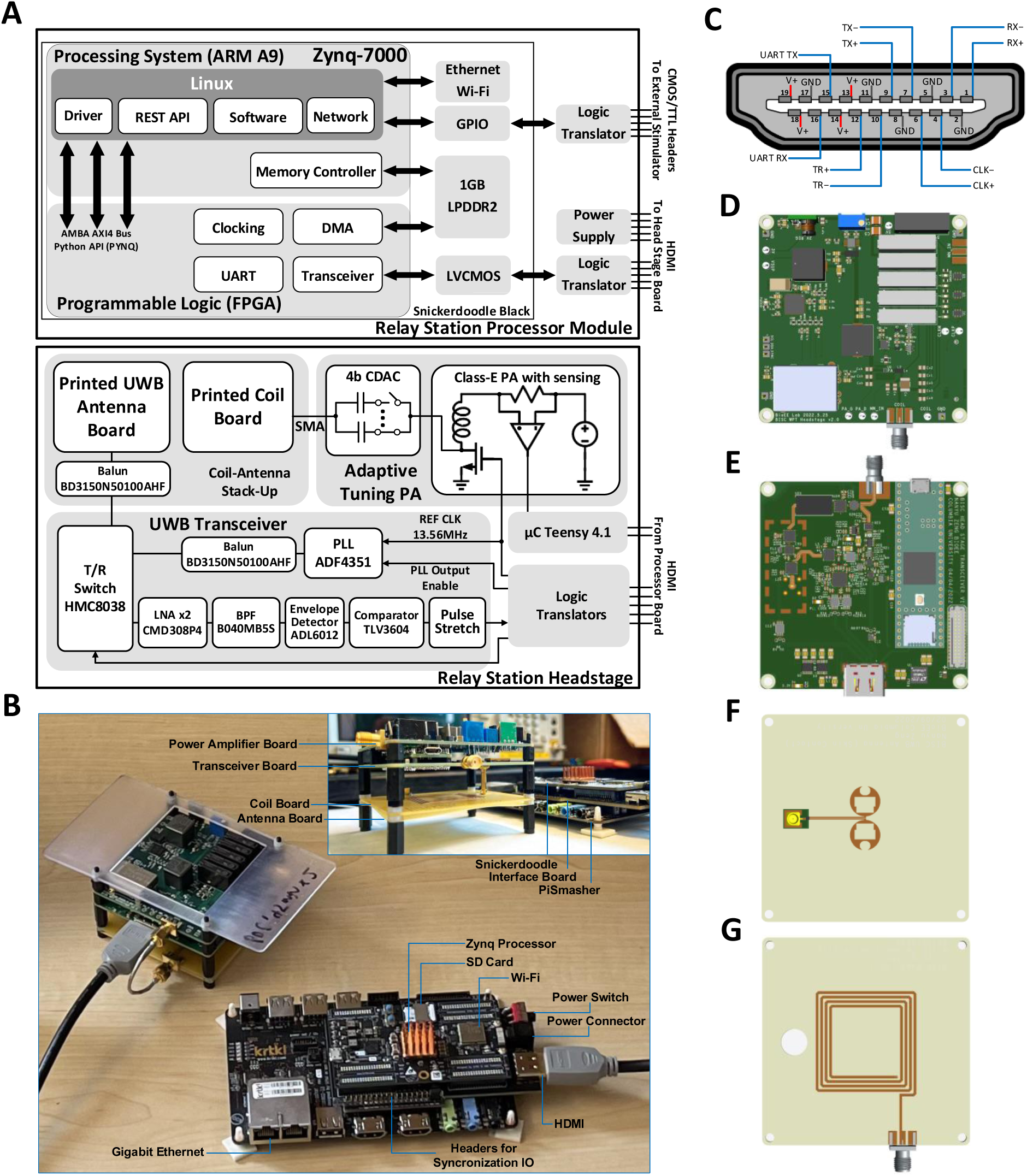
Architecture of the relay station. (**A**) Functional diagram of the modules in the relay station. (**B**) Photos of the relay station with cabling. (**C**) HDMI connector pinout diagram. (**D**) Power-amplifier board. (**E**) Transceiver board, with a Teensy 4.1 board (the top SMA connector is used for testing only). (**F**) UWB antenna board. (**G**) Coil board.

**Fig. S3.**
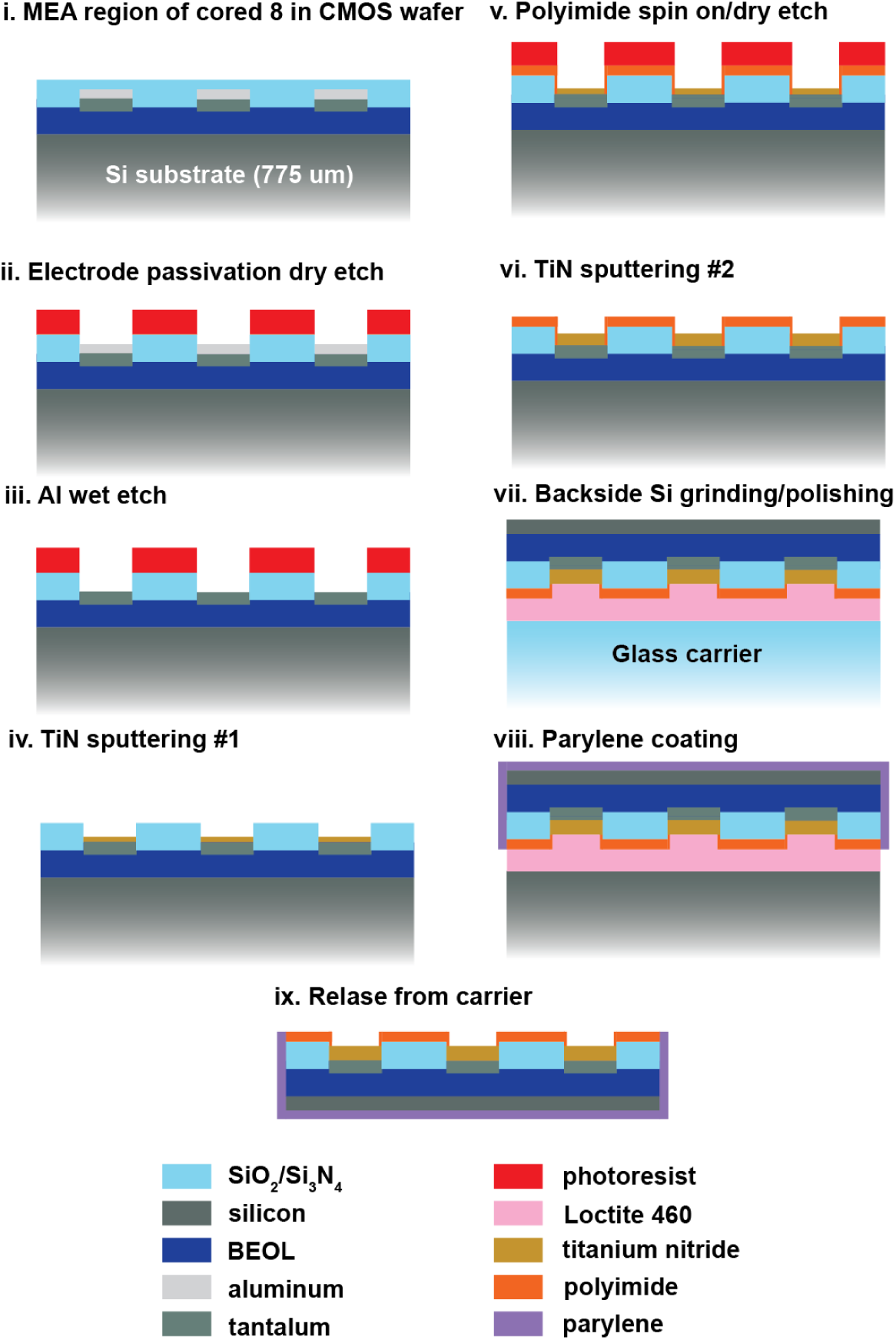
Coupon processing. A cross-section through the microelectrode array (MEA) region shows the pristine wafer with bulk Si substrate (i) post-processed to expose the electrodes (ii and iii) followed by lift-off patterning of a protective TiN layer (iv), spin-on polyimide encapsulation (v), surface impedance optimization with a second lift-off patterned TiN layer (vi), coupon dicing, thinning to remove the bulk of the substrate Si down to ∼25 µm RST (remaining silicon thickness) (vii), and finally parylene encapsulation (viii) and solvent release (ix).

**Fig. S4.**
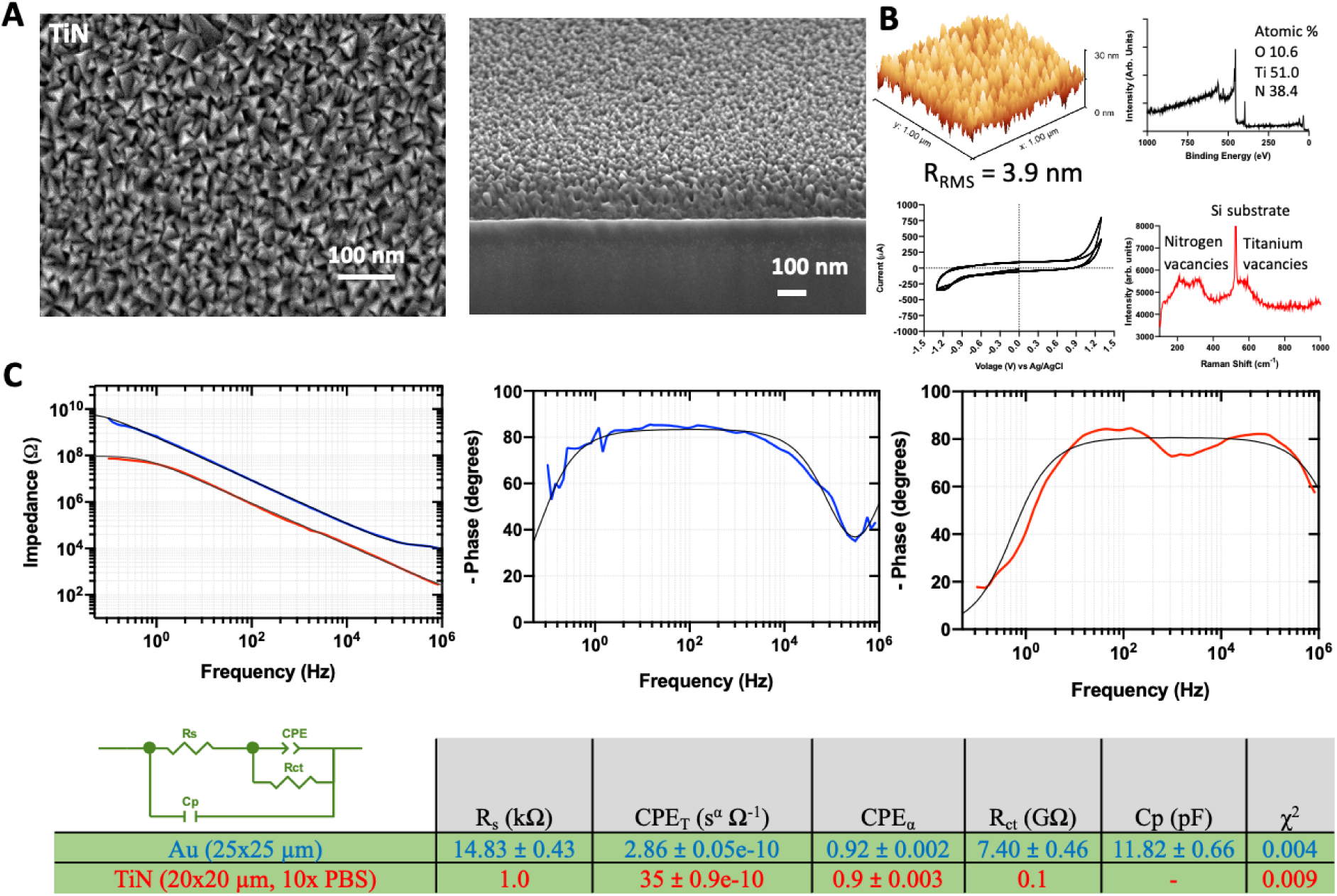
Titanium nitride characterization. (**A**) The characteristic roughened morphology of the deposited TiN films was observed with SEM. SEM images of TiN films were acquired on a FEI Helios dual beam FIB-SEM. (**B**) Surface roughness of the deposited TiN was quantified using AFM giving a root mean square roughness of 3.9 nm. Surface roughness was measured with a Bruker Dimension Icon AFM in tapping mode. Raman scattering was measured on a Renishaw inVia Raman microscope. XPS elemental analysis was performed on a PHI VersaProbe II with peak analysis done using PHI MultiPak software. (**C**) Electrochemical impedance spectroscopy was used to verify the reduction in electrode impedance from the TiN films. A 20 mV amplitude sinusoidal applied voltage was swept from 10^6^ to 10^-1^ Hz with a large area Pt coil counter electrode and Ag/AgCl reference electrode. A 20 µm × 20 µm working TiN test electrode (red) was measured in 10× PBS. For comparison, a 25 µm × 25 µm Au electrode (blue), measured in 1x PBS is shown as well. The TiN CV curves were measured using a 1 cm^2^ TiN working electrode, Ag/AgCl pellet reference electrode, and Pt coil counter electrode in 1× PBS between 1.3 and −1.3 V at a scan rate of 100 mV/s. Electrochemical measurements were made using a CH Instruments 760D potentiostat and curve fitting to the equivalent circuit model shown here consisting of a double layer capacitor (constant phase element *CPE* with leakage resistance *R_ct_*) at the electrode surface, solution resistance *R_s_*, and parasitic capacitance *C_p_* was done with ZView software (Scribner Associates).

**Fig. S5.**
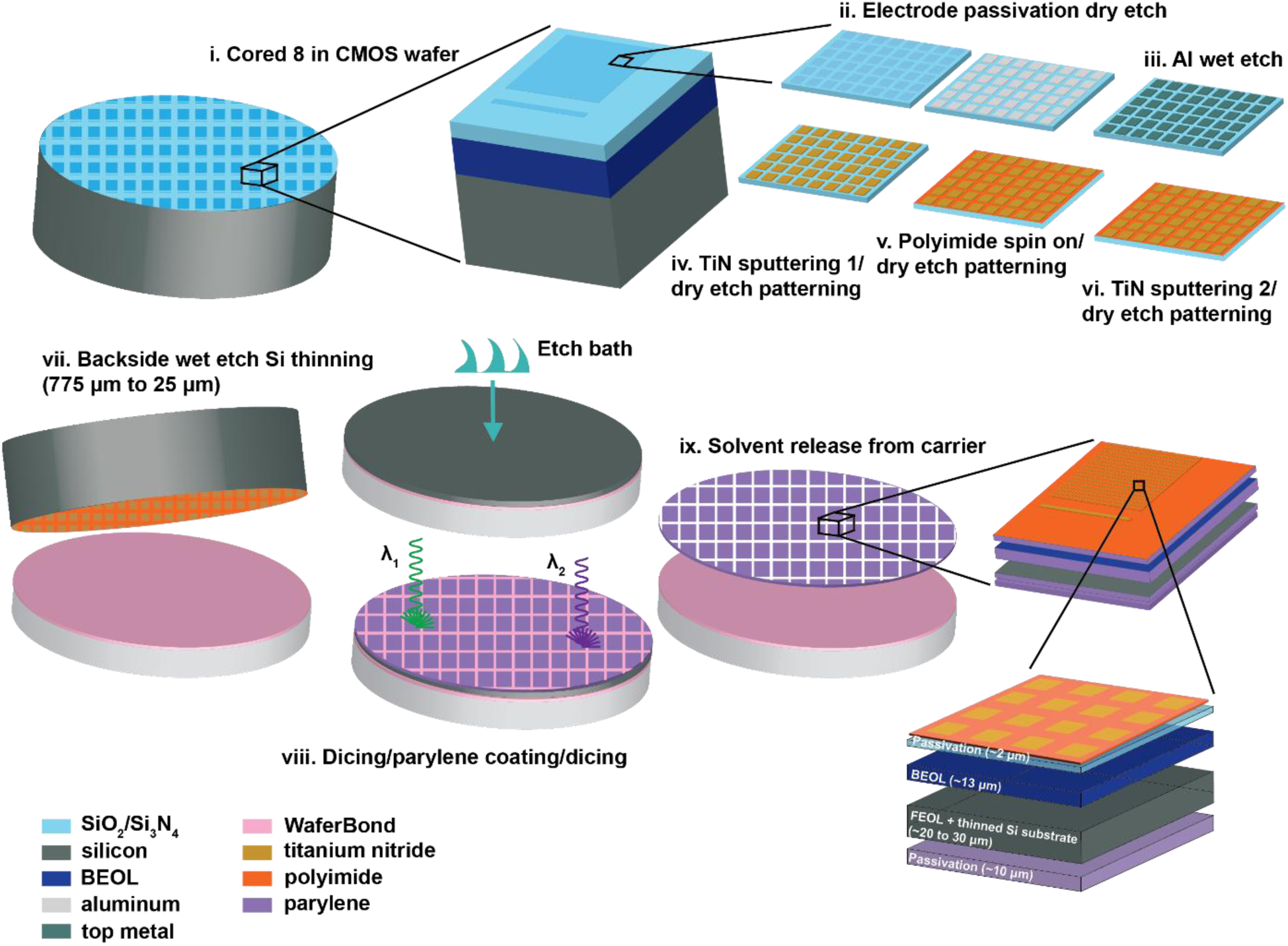
Wafer-scale processing. Proceeding in a similar manner to coupon processing (i – vi), 200-mm (8”) wafers cored from 300-mm foundry wafers receive a wet etch thinning step (vii) followed by laser dicing (viii) to yield more than 100 implant chips per 200-mm wafer.

**Fig. S6.**
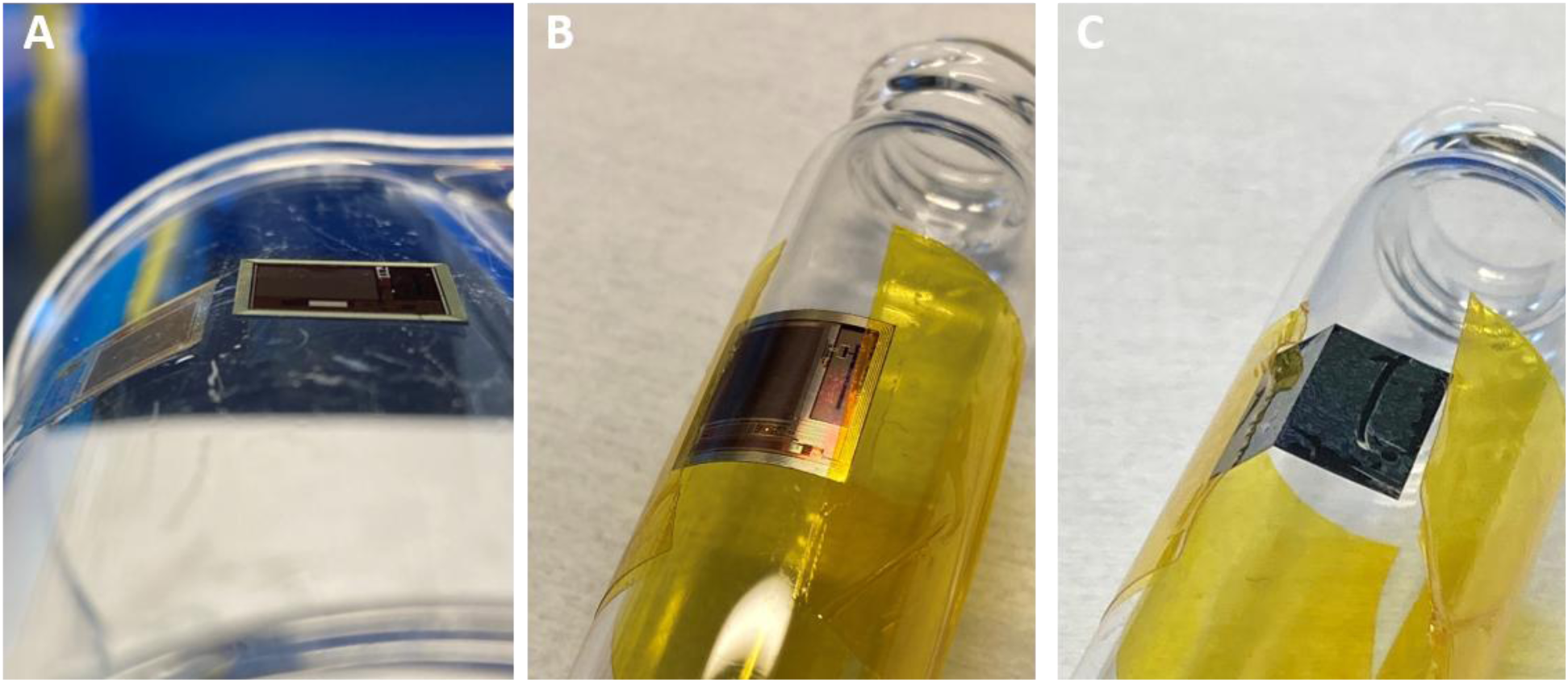
Mechanical flexibility of thinned BISC chips. (**A**) Image of a thinned chip (left) spontaneously conforming to the surface of a glass beaker with a radius of curvature of ∼30 mm using water as the wetting liquid next to an unthinned, rigid chip (right). (**B**) Image of a chip bent to a curvature of ∼10 mm in the more robust direction (front surface under tension) below the spontaneous elasto-capillary length cutoff. Kapton tape is used to supply the necessary bending force. No fracturing occurs in the silicon down to this curvature. (**C**) Bending down to 10-mm radius in the opposite direction where the silicon experiences greater tensile stresses due to the position of the neutral plane, resulting in brittle fracture.

**Fig. S7.**
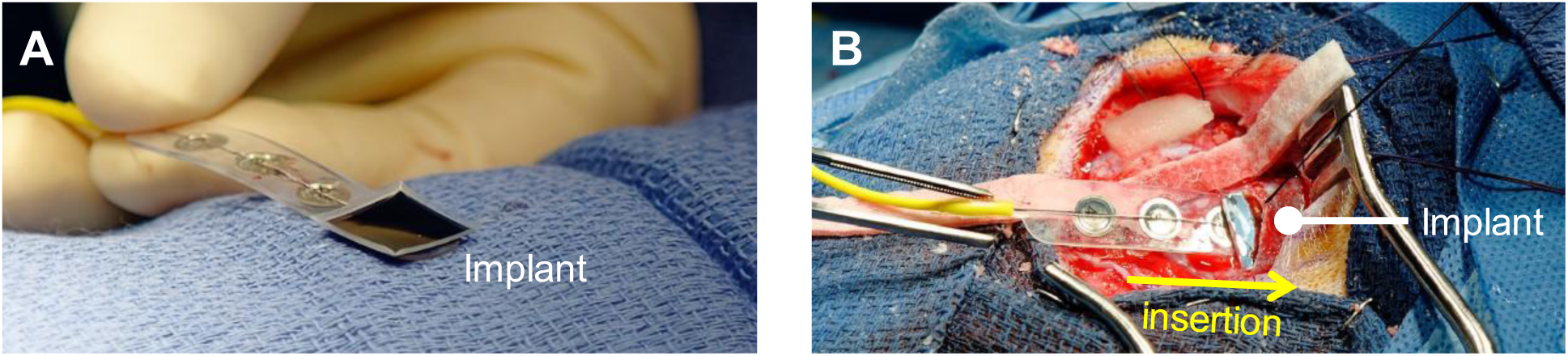
Surgical insertion of the implant. (**A**) BISC device placed faced down on commercial strip electrodes, used as the insertion shuttle. (**B**) Device insertion to the subdural space.

**Fig. S8.**
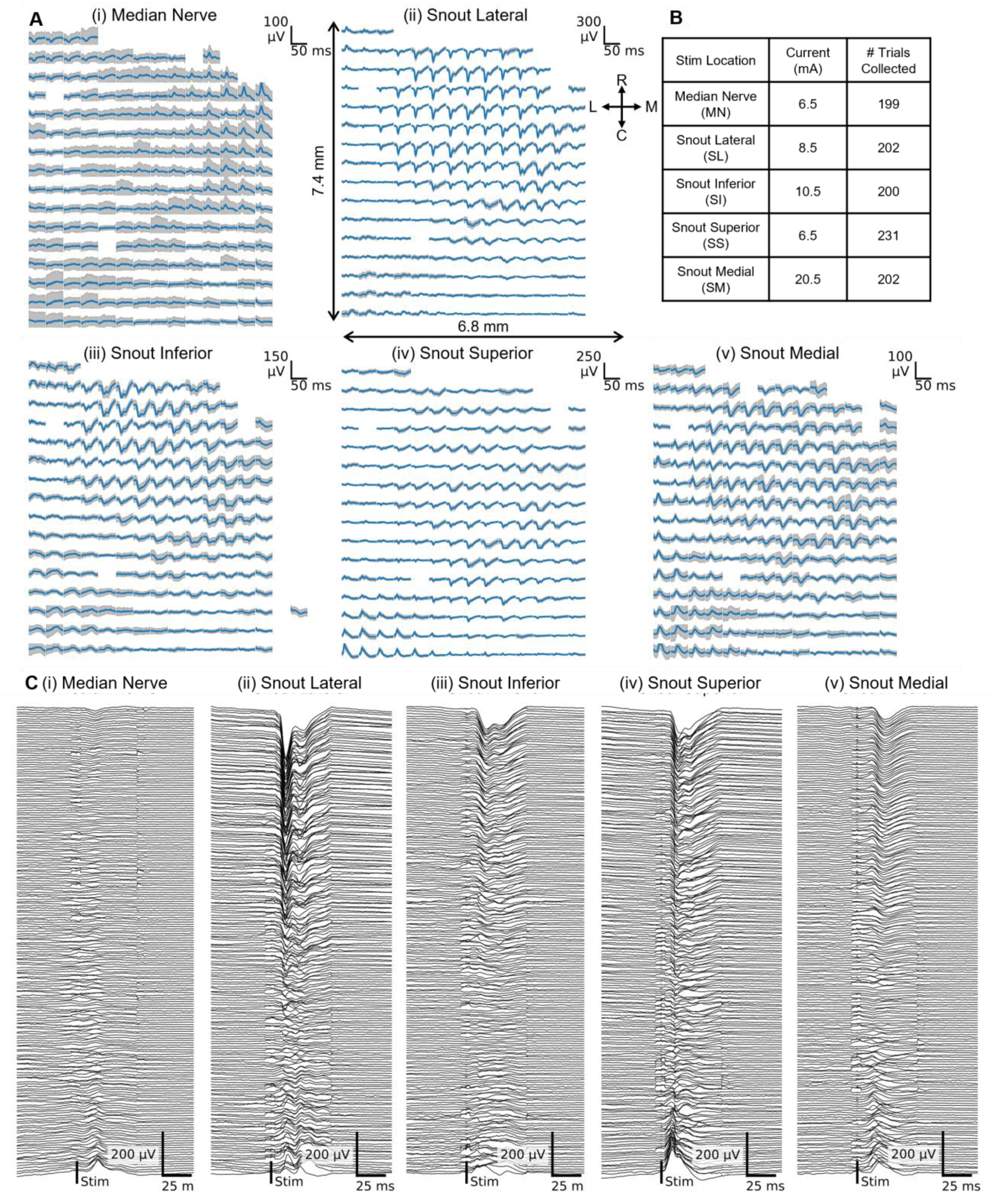
Somatosensory evoked potential (SSEP) recording from porcine model. (**A**) Trial averaged (n = 100) SSEPs evoked from five different stimulation locations. Gray shade indicates SD. (**B**) Stimulation current amplitude (2.79 Hz, pulse width 0.3 ms) applied to each location, and total number of SSEP trials recorded from each location. (**C**) Representation of (A) aligned on the same time axis.

**Fig. S9.**
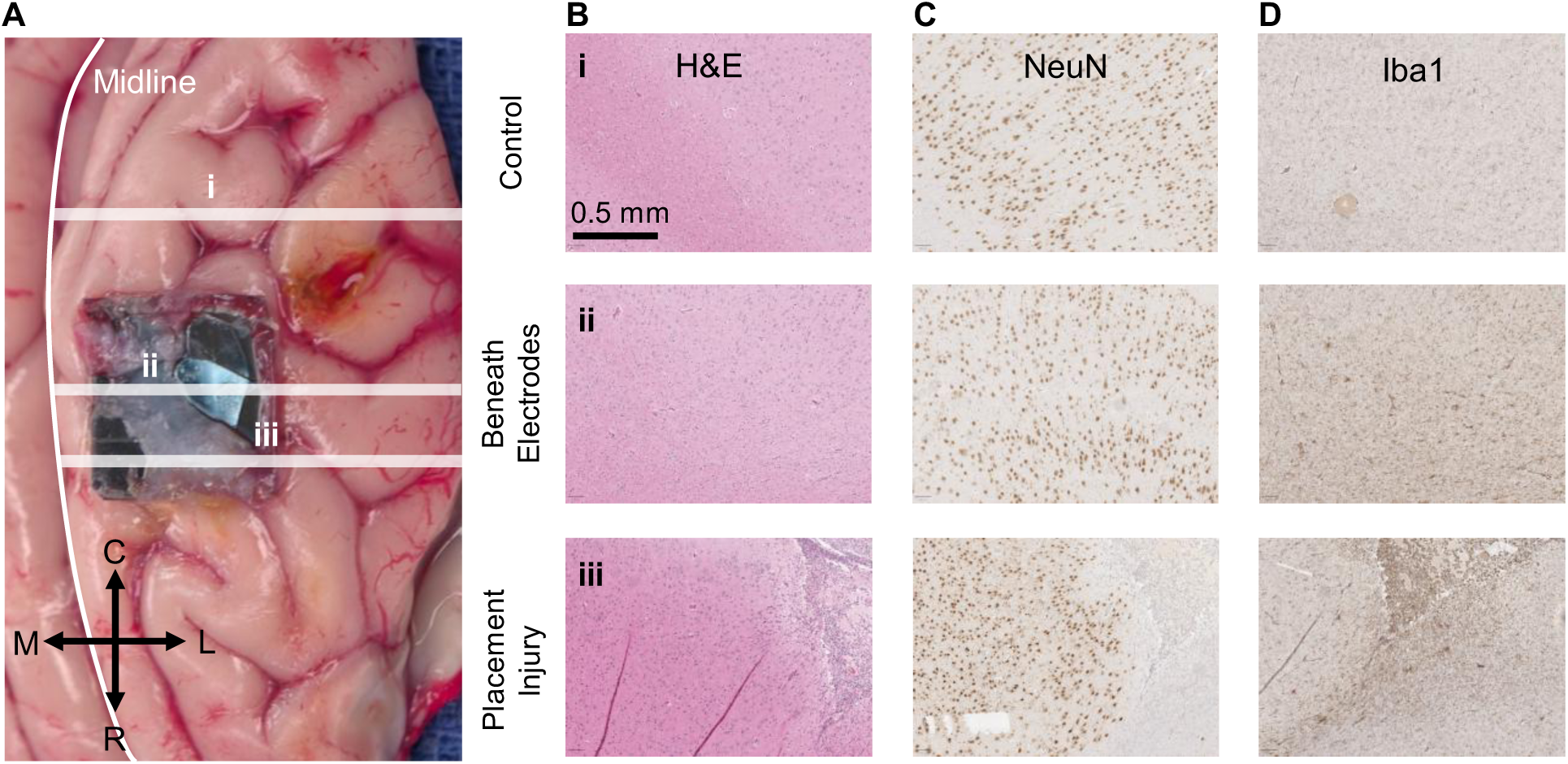
Post-mortem histological analysis of the porcine brain. (**A**) Photograph of the extracted brain showing rough estimates of the location of histological sections (control (i), beneath electrode array (ii), near device perimeter (iii)). The device fractured but remained intact during the extraction. Images of histological stainings for (**B**) hematoxylin and eosin (H&E), (**C**) NeuN, (**D**) Iba1. Histological sections (5-µm thick) of the left occipital cortex (i), taken as a control sample, show no discernable pathological changes. Sections from the left frontal cortex taken directly beneath the electrodes (ii) show no significant pathology by H&E stain or an immunoperoxidase stain for NeuN. However, a mild reactive microgliosis is highlighted with an immunoperoxidase stain for Iba1. Sections from the left frontal cortex taken at the site of a placement injury (iii) show a small focus of cortical tissue loss by H&E stain, consistent with mechanical injury. An immunoperoxidase stain for NeuN highlight the focal loss of neurons, and an immunoperoxidase stain for Iba1 highlight macrophages and reactive microglia associated with the cortical lesion.

**Fig. S10.**
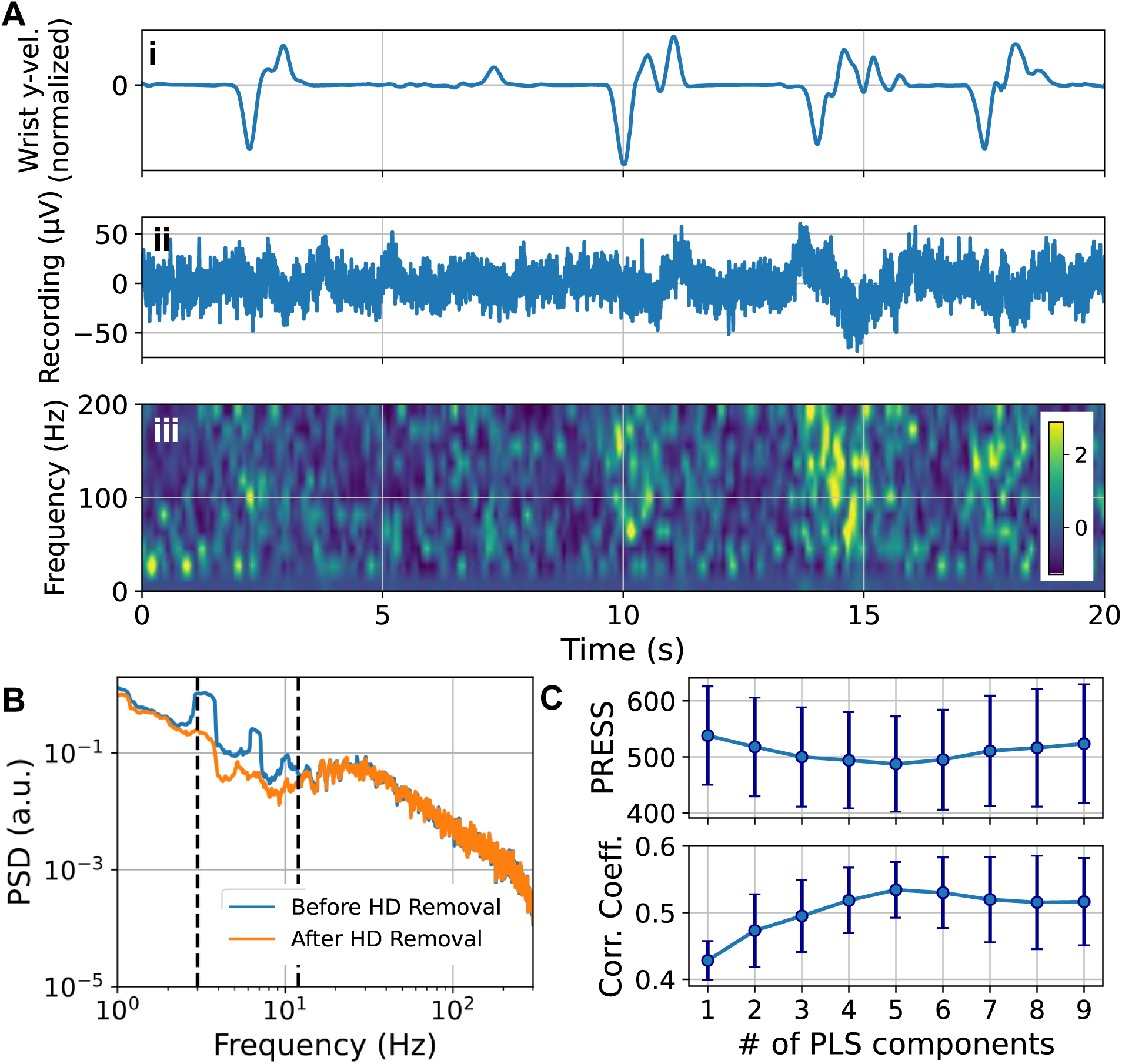
Motor feature decoder optimization and example channel recording. (**A**) Example representation of a band-pass filtered (0.3 – 300 Hz) channel recording (ii) and spectrogram (iii) aligned to the subject’s wrist velocity in y-direction (front-back) (i). (**B**) Example representation of a channel power spectral density (PSD) before and after removing hemodynamics (HD). Dashed lines indicate the band-pass filter range (3 – 12 Hz) that was applied to extract the HD time series components. (**C**) Linear parametric sweep of the number of partial least squares (PLS) components, and the resulting predictive error sum of squares (PRESS) (top) and Pearson’s correlation coefficient (bottom) between observed motor feature (normalized wrist velocity in y-direction) and PLS prediction. Error bars indicate SE. Plots in (A-ii), (A-iii) and (B) after HD removal represent the same data.

**Fig. S11.**
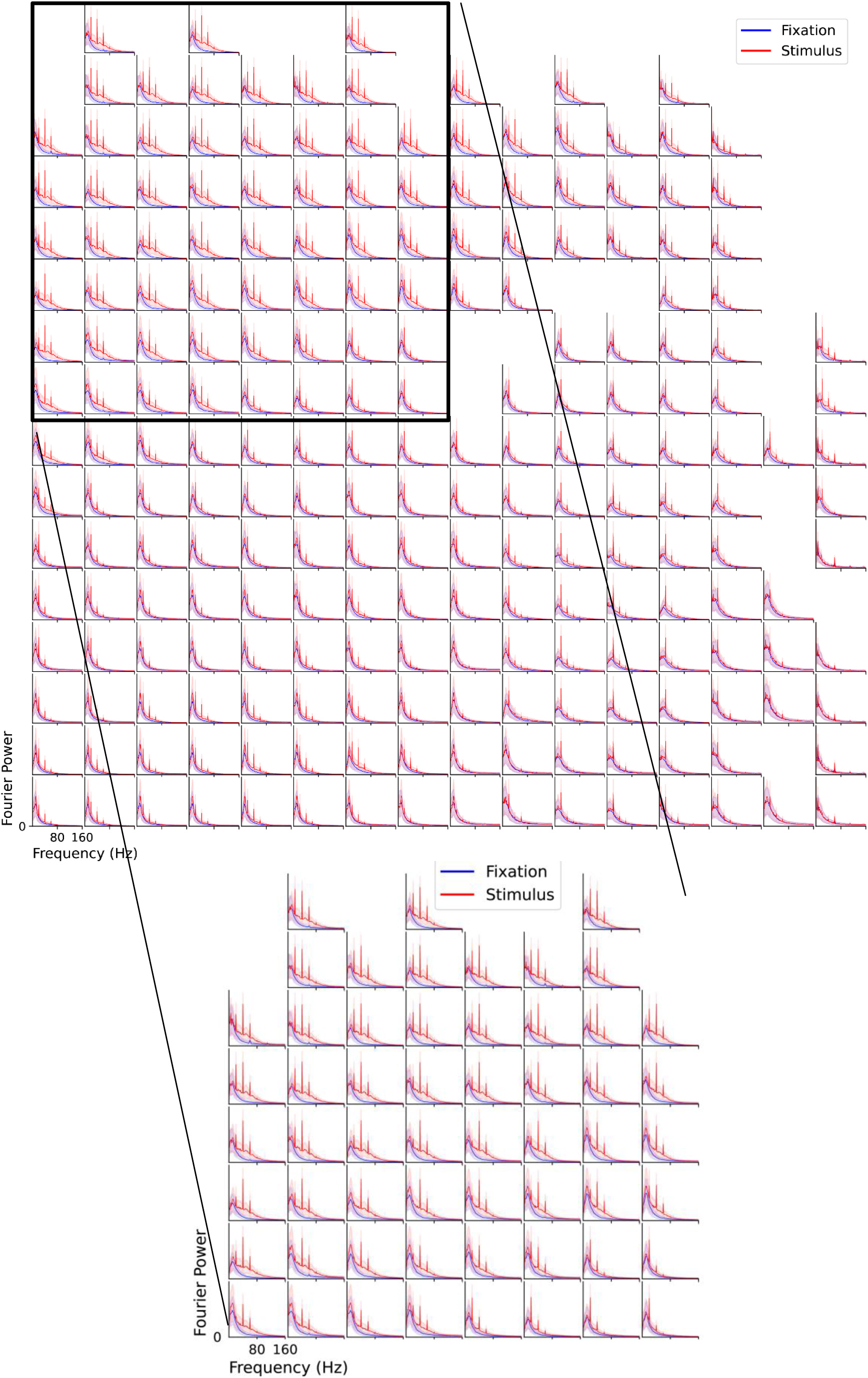
Comparison of Fourier spectrum during fixation and grating stimuli for each channel. Each panel corresponds to one non-saturated channel. Solid lines are the trial average of Fourier spectrum, and shaded areas mark the standard deviation across trials. Y axis scales are different for each panel, however they all start from 0. Channels over V1 (inset) show more pronounced grating stimuli-induced response, which agrees with the dot stimuli-induced findings (see **Fig. 5D-E, G**).

**Fig. S12.**
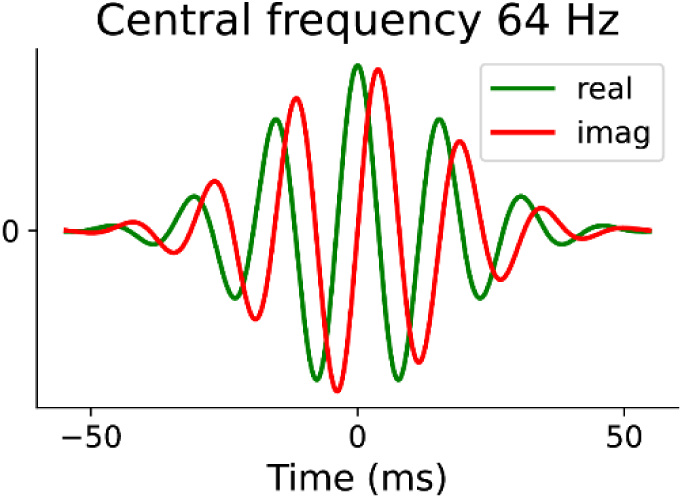
Complex Morlet wavelet example with central frequency as 64 Hz.

**Fig. S13.**
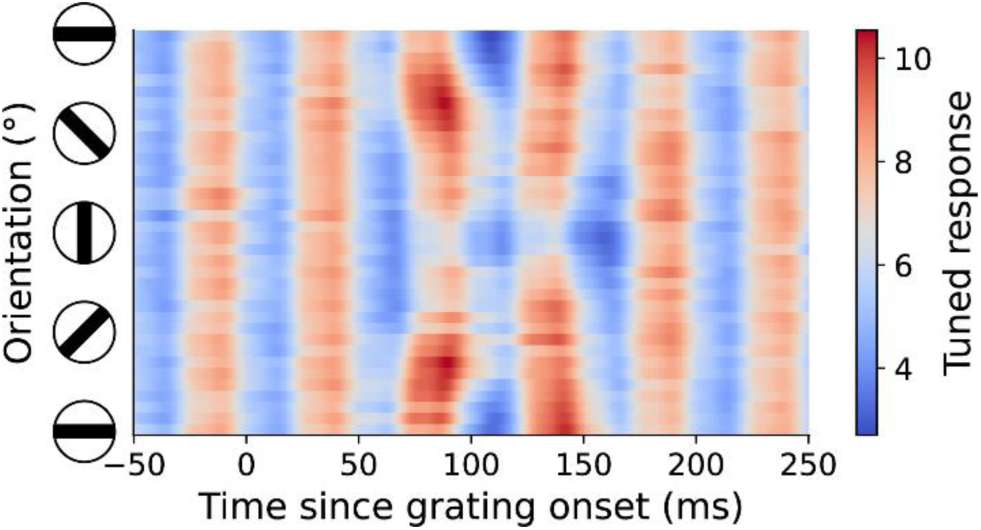
Grating-triggered response. Raw grating-triggered-average result of an example channel for the frequency band centered at 64 Hz.

**Fig. S14.**
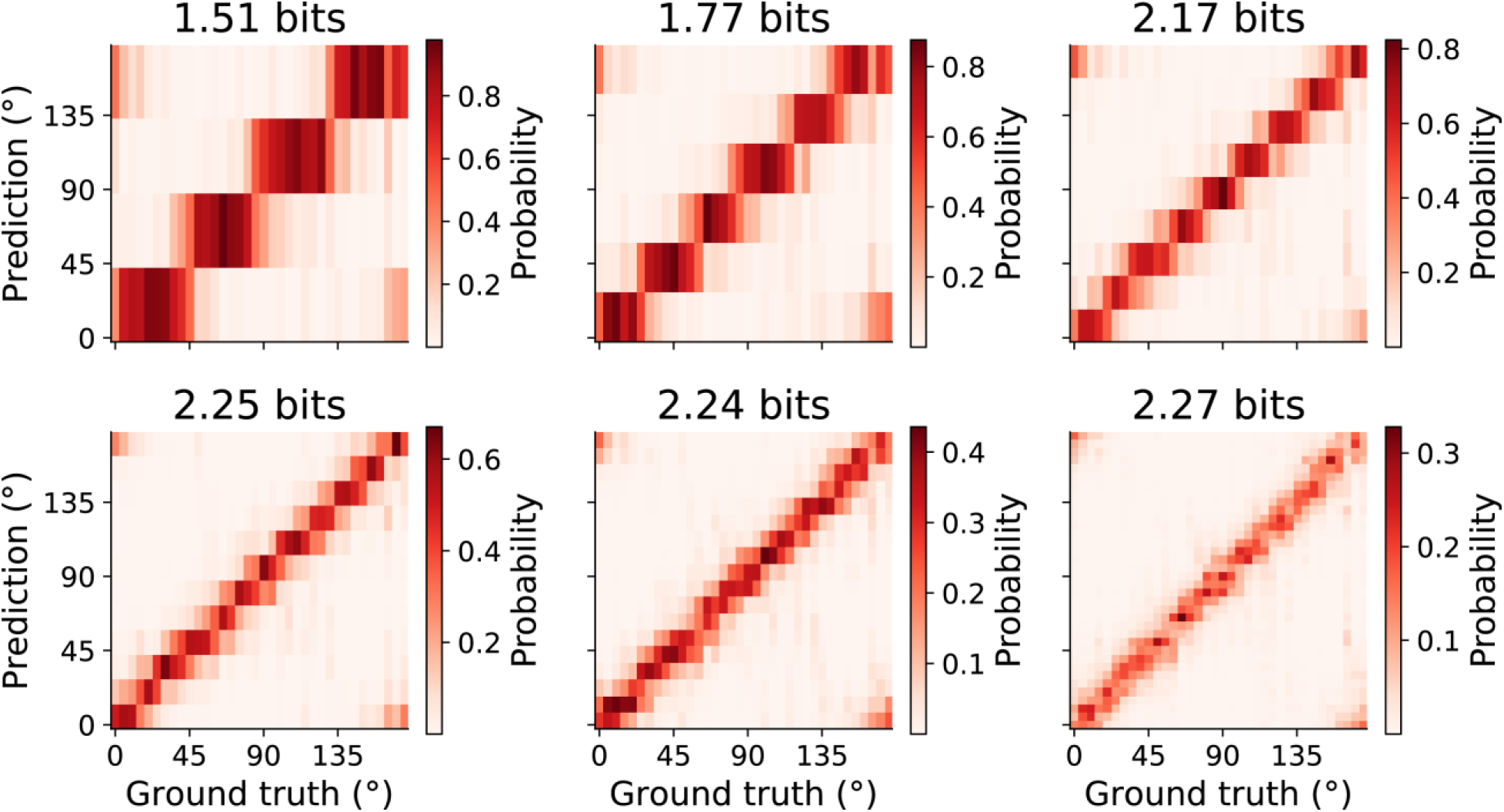
Mutual information between decoder prediction and BISC response for different orientation discretization. From left to right, top to bottom, number of bins used to discretize [0°, 180°) gradually increases.

**Fig. S15.**
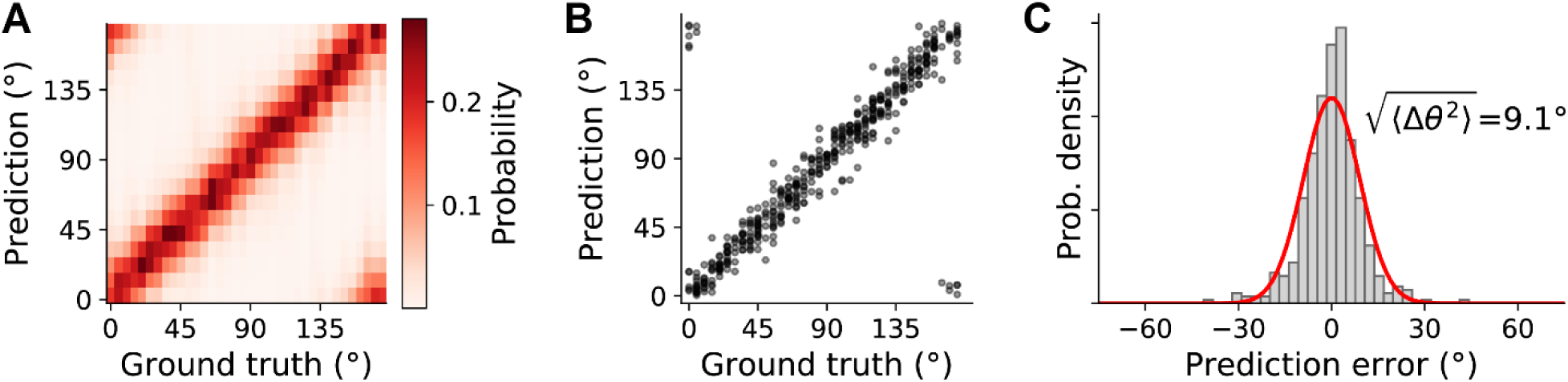
Orientation decoder trained as a regressor. The training objective is the Euclidean distance between the circular mean of decoder output and the ground truth orientation in the complex space. Decoder output (**A**), point estimation (**B**) and error histogram (**C**) are shown as in Fig. 5L**-M**.

**Fig. S16.**
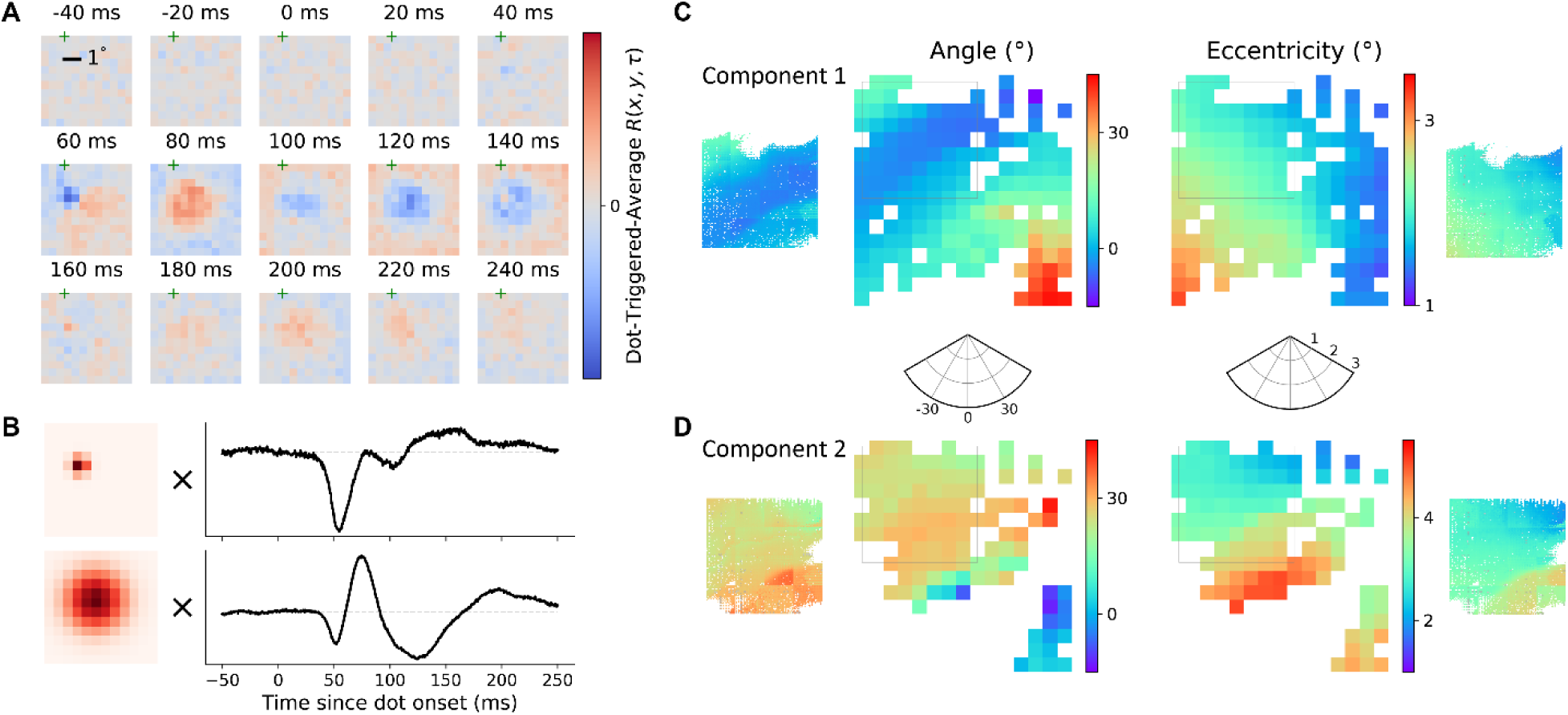
Receptive fields (RF) estimated from unfiltered response and the retinotopic maps of two components. (**A**) Dot-triggered-average of one example channel response without band-passed filtering. (**B**) Two component decomposition of the receptive field in (A). Each component is time-space separated and the spatial profile is approximated by 2D Gaussian function. (**C**) RF location of the first component, i.e. the one with smaller spatial extent. Dense recordings on one corner of the chip are analyzed in the same fashion, shown in inset figures on both sides. (**D**) RF location of the second component, i.e. the one with larger spatial extent.

**Fig. S17.**
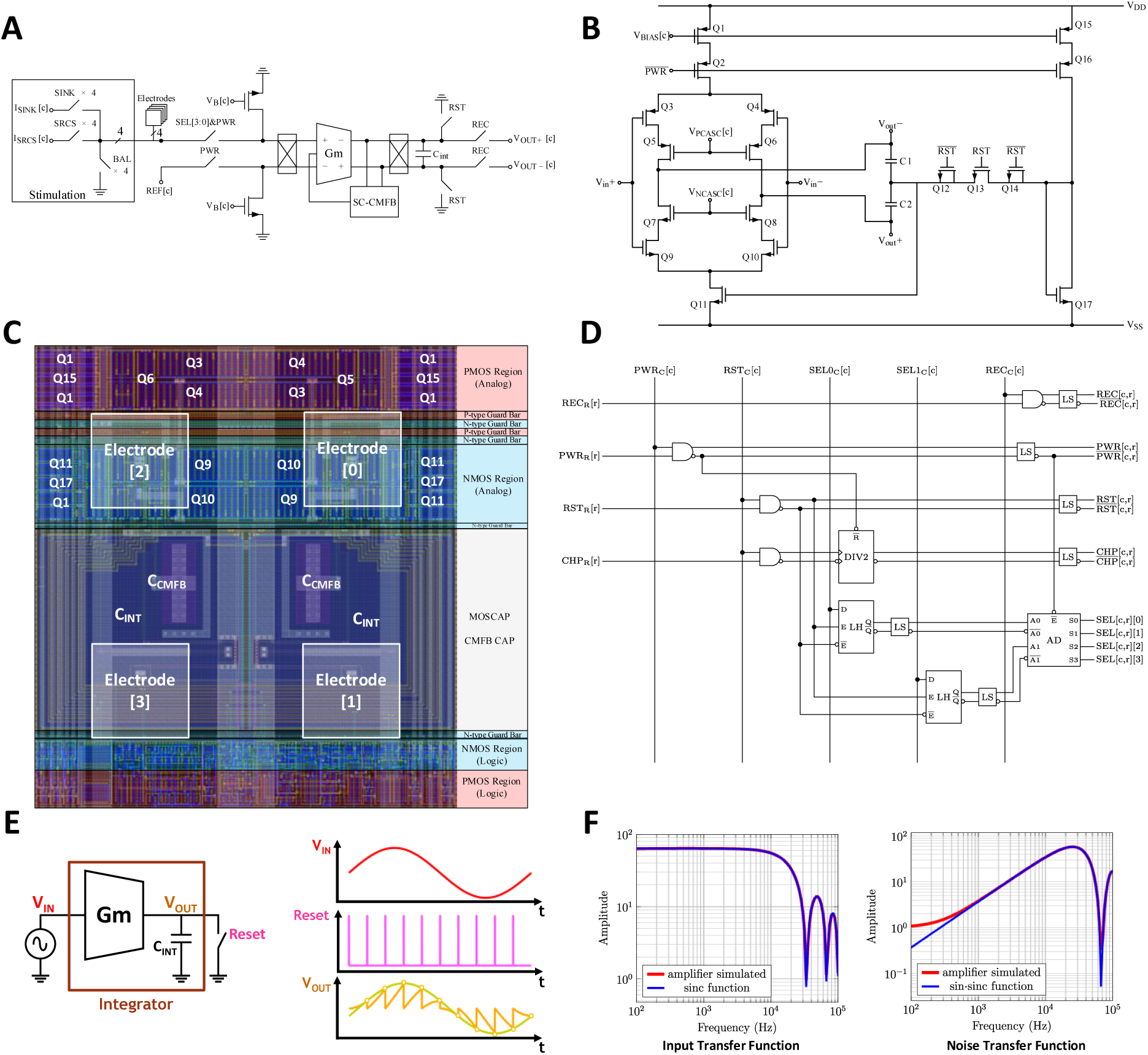
Architecture of the pixel. (**A**) Functional diagram of pixel. (**B**) Schematic of the Gm-C integrator, implemented as an inverter-based amplifier. (**C**) Layout of the pixel. (**D**) Schematic of digital logic used for recording in each pixel. [c] and [r] correspond to the pixel’s column and row addresses. (**E**) Depiction of the boxcar averaging principle used for amplification. The input is a sine wave, and the output switch resets periodically for a very short time. If the reset frequency is much higher than the input signal frequency, the output of the integrator becomes a sampled, amplified version of the input. (**F**) The theoretical and simulated transfer functions of the amplifier input and noise.

**Fig. S18.**
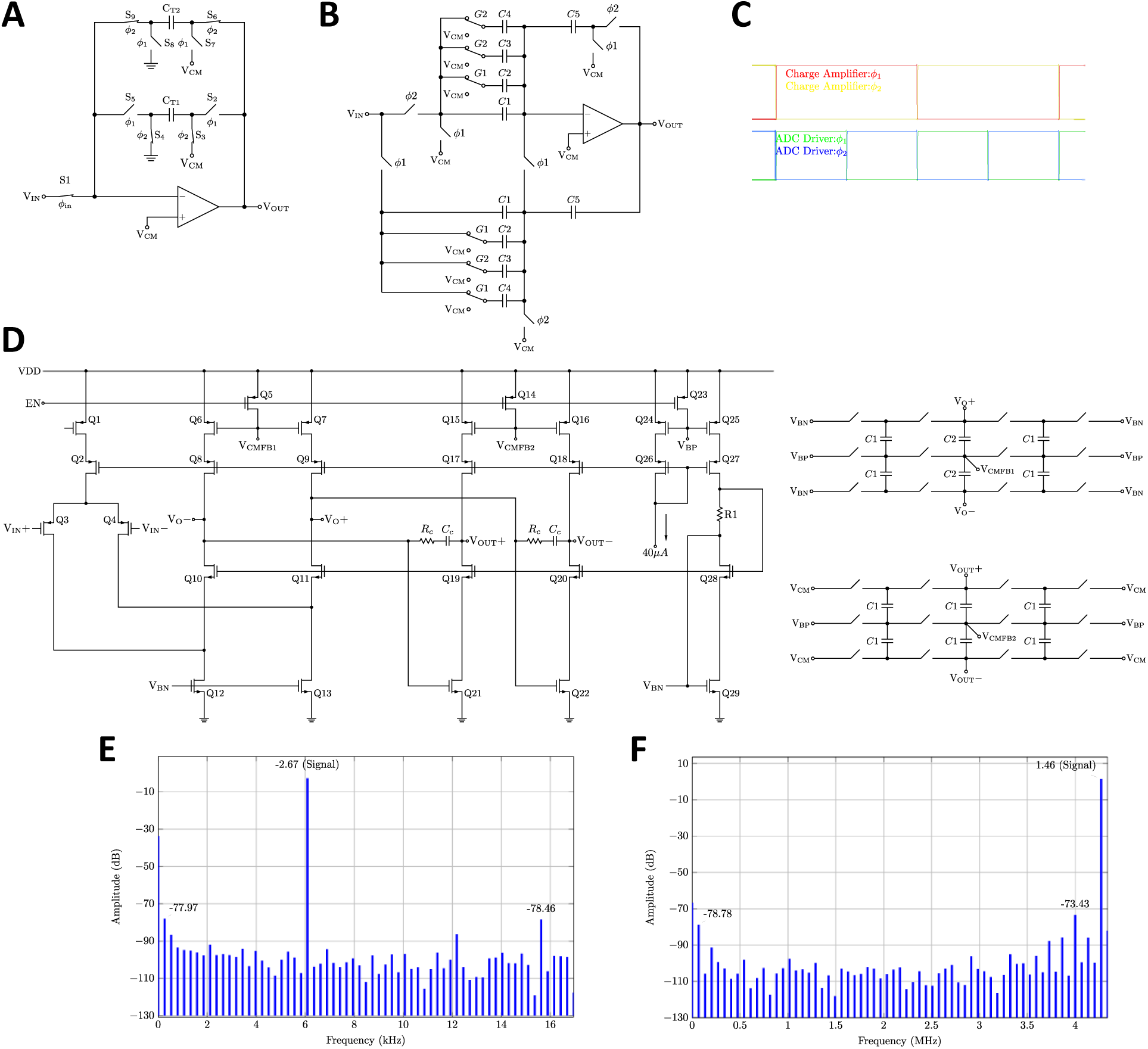
Design and characterization of the programmable gain amplifier (PGA). (**A**) Design of a ping-pong charge amplifier. This is the first stage of the PGA. This charge amplifier moves the charges integrated inside the pixels to its transfer capacitors C_T1_ and C_T2_. The ping-pong operation eliminates the need for two charge amplifiers, thereby saving power. A single ended version is shown for clarity while the actual implementation is differential. The operational amplifier (OPAMP) is implemented as the folded cascaded operational transconductance amplifier (OTA) in (D). This OTA has cascaded current sources at the second stage to boost DC gain. (**B**) Design of a programmable gain ADC driver. This is the second stage of the PGA following the charge amplifier. Correlated double sampling is used to enhance the linearity as described in ^112^. A single-ended version is shown for clarity and the OTA is the same as (D) without the cascoding at the second stage. (**C**) The timing diagram of the switches in (A) and (B). (**D**) The OTA design with common-mode feedback. (**E**) The simulated linearity of the neural amplifier and the charge amplifier. Spurious free dynamic range (SFDR) is over 75 dB. (**F**) The simulated linearity of the PGA. SFDR is over 71 dB.

**Fig. S19.**
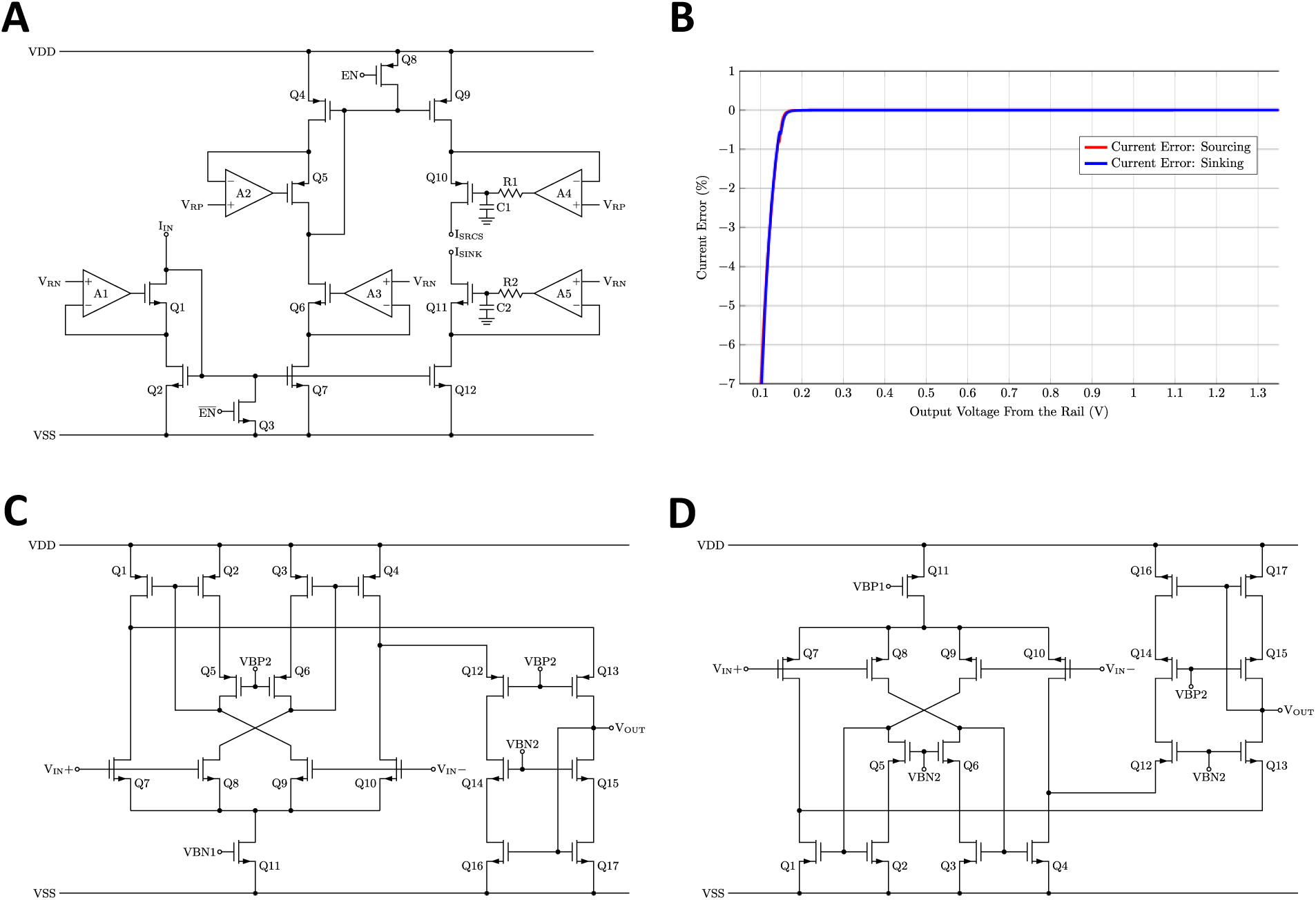
Design and characterization of the stimulation current generator. (**A**) Design of the regulated cascode current driver. Operational amplifier (OPAMP) A1, A3, A5 are implemented as PMOS recycling folded cascode input operational transconductance amplifier (OTA) and A2, A4 are implemented as NMOS input recycling folded cascode OTA. R1, R2 and C1, C2 are used to protect the output of OPAMP A4 and A5 by filtering the voltage spikes through the parasitic drain-to-gate capacitance of Q10 and Q11, when the stimulation polarity changes during monophasic stimulation. (**B**) The simulated voltage compliance under highest (1.12 mA, worst-case) stimulation current. The current error is less than 1% when the output voltages of the current driver are more than 150 mV from the rails (−1.35 V to 1.35 V). (**C**) The PMOS input recycling folded cascode OTA, based on ^130^. Compared to simple a 5-Transistor OTA, this OTA offers higher DC gain and gain-bandwidth product for the same power consumption. (**D**) The NMOS input recycling folded cascode OTA.

**Fig. S20.**
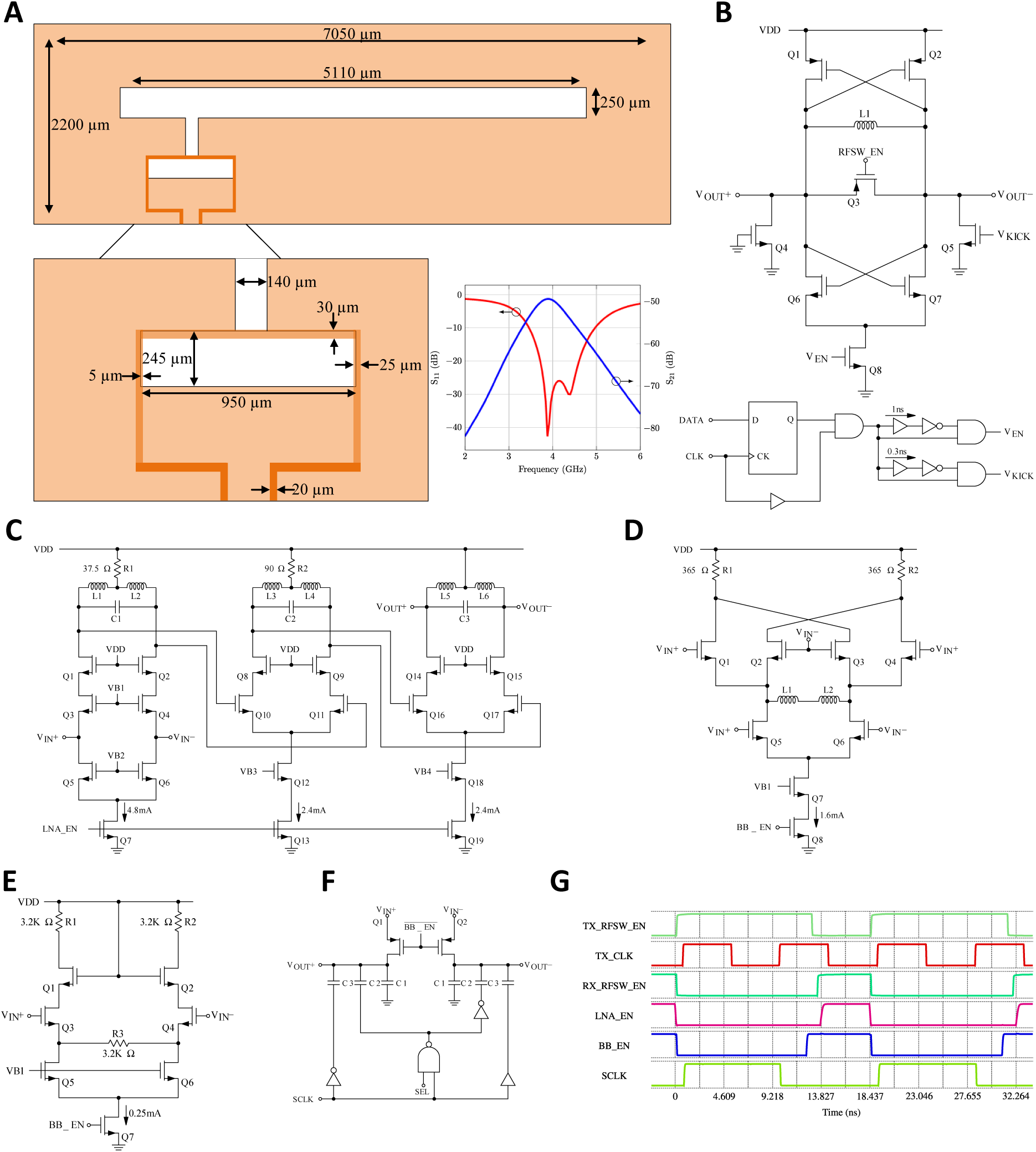
Design and characterization of the wireless transceiver. (**A**) Design of the on-chip ultra-wideband (UWB) antenna and simulated S-parameters. The feed ring is made of the redistribution layer (RDL) metal, and the bottom plate is a stack of METAL6 to METAL1; small metal slots are made in the bottom plate to satisfy density rules (not shown) (**B**) UWB transmitter based on LC complementary oscillator, and a V_EN_, V_KICK_ pulse generator. (**C**) Three-stage inductor loaded low noise amplifier (LNA). (**D**) Self-mixing mixer. (**E**) Baseband low-pass amplifier. (**F**) Charge redistribution sampling stage for subtracting a threshold voltage from the output of (E), with one-bit programmability. (**G**) Timing diagram of the wireless transceiver.

**Fig. S21.**
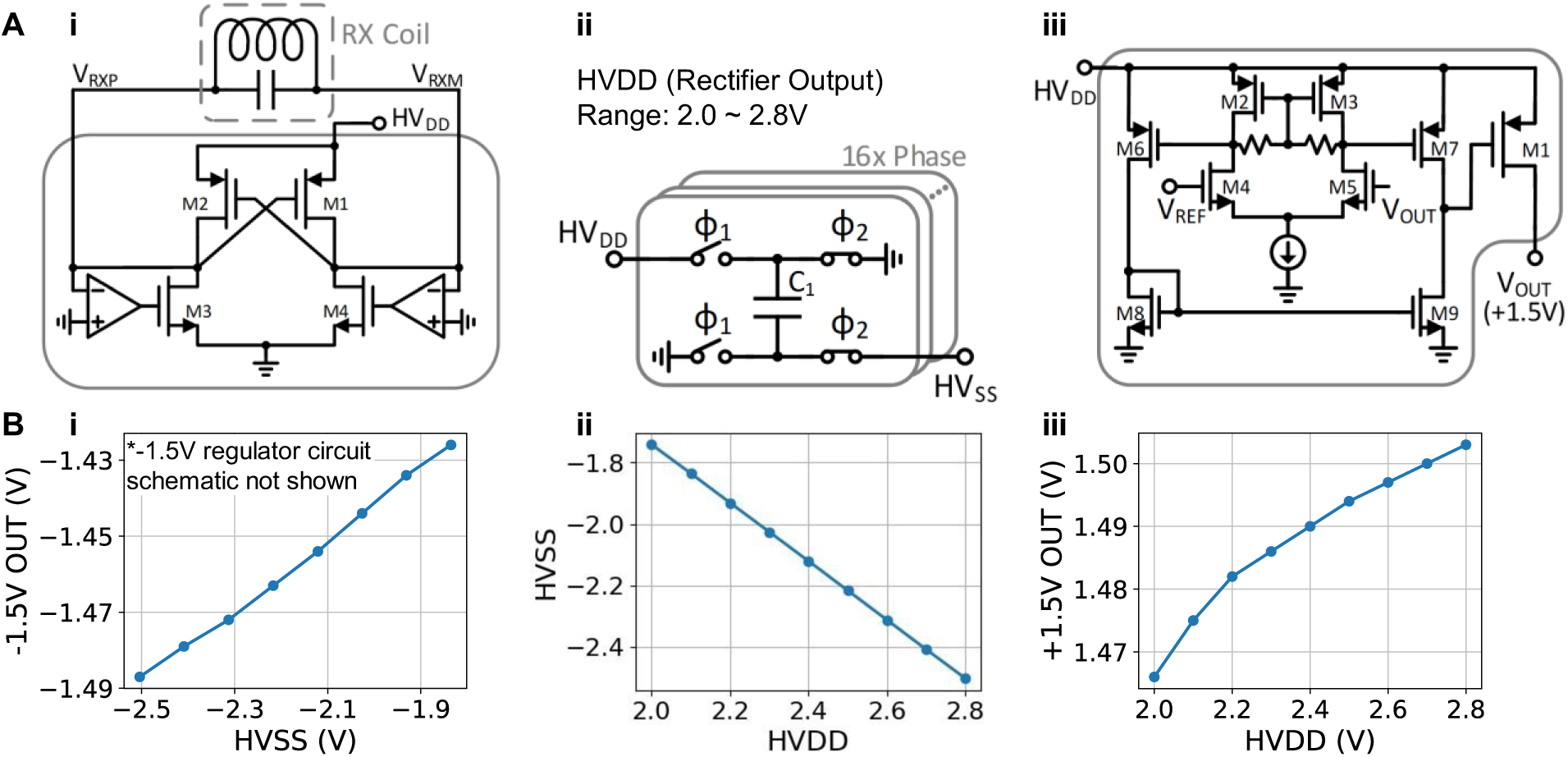
Design and characterization of the wireless power transfer circuit. (**A**) Key circuit blocks: active rectifier (i), switch cap DC-DC converter (ii), +1.5V regulator (iii). (**B**) Measured DC-DC line regulation of the key blocks: −1.5V regulator (i), switch cap DC-DC converter (ii), +1.5V regulator (iii). HV_DD_ denotes the rectifier DC output, and HV_SS_ denotes DC-DC converter output.

**Fig. S22.**
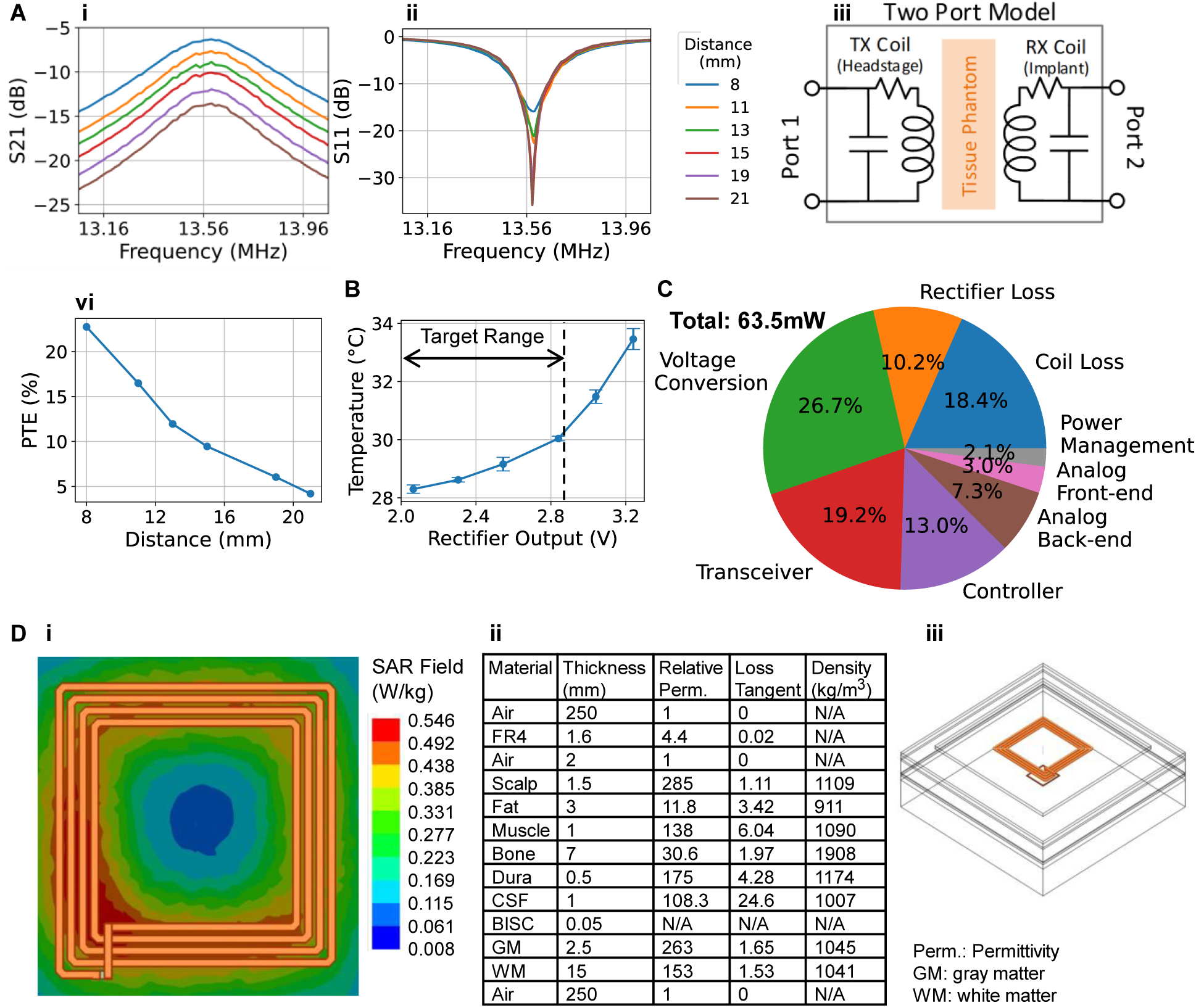
Power, thermal, and specific absorption rate (SAR) characterization. (**A**) Coil link characterization with two port S-parameter measurement. S21 (i) and S11 (ii) were measured with chicken breast as tissue phantom (iii), assuming an ideal conjugate matched impedance on the transmitting side (Port 1) and a 75 Ω load on the receiving side (Port 2) which is the linear load equivalence of the overall circuit load. S21 is converted to power transfer efficiency (PTE) for these loads (iv). (**B**) Temperature measurement of a fully passivated device in recording mode under different power delivery conditions. (**C**) Power consumption breakdown of the device when it is in recording mode. (**D**) SAR simulation result (i) assuming 1.5-cm implant depth and six-layer brain model (ii), (iii).

**Fig. S23.**
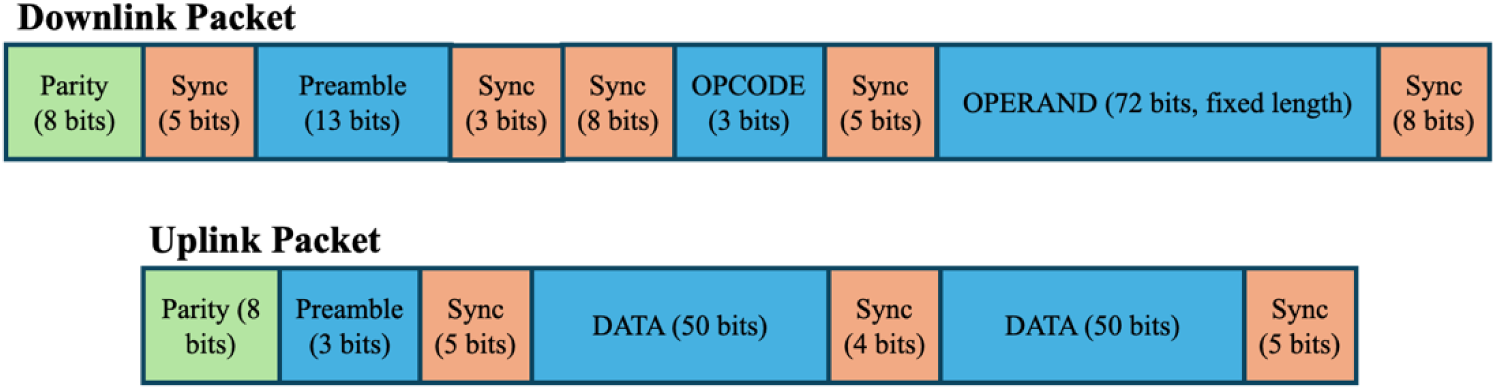
Communication packets in the BISC system. Downlink for the direction from the relay station to the implant. Uplink for the direction from implant to relay station.

**Fig. S24.**
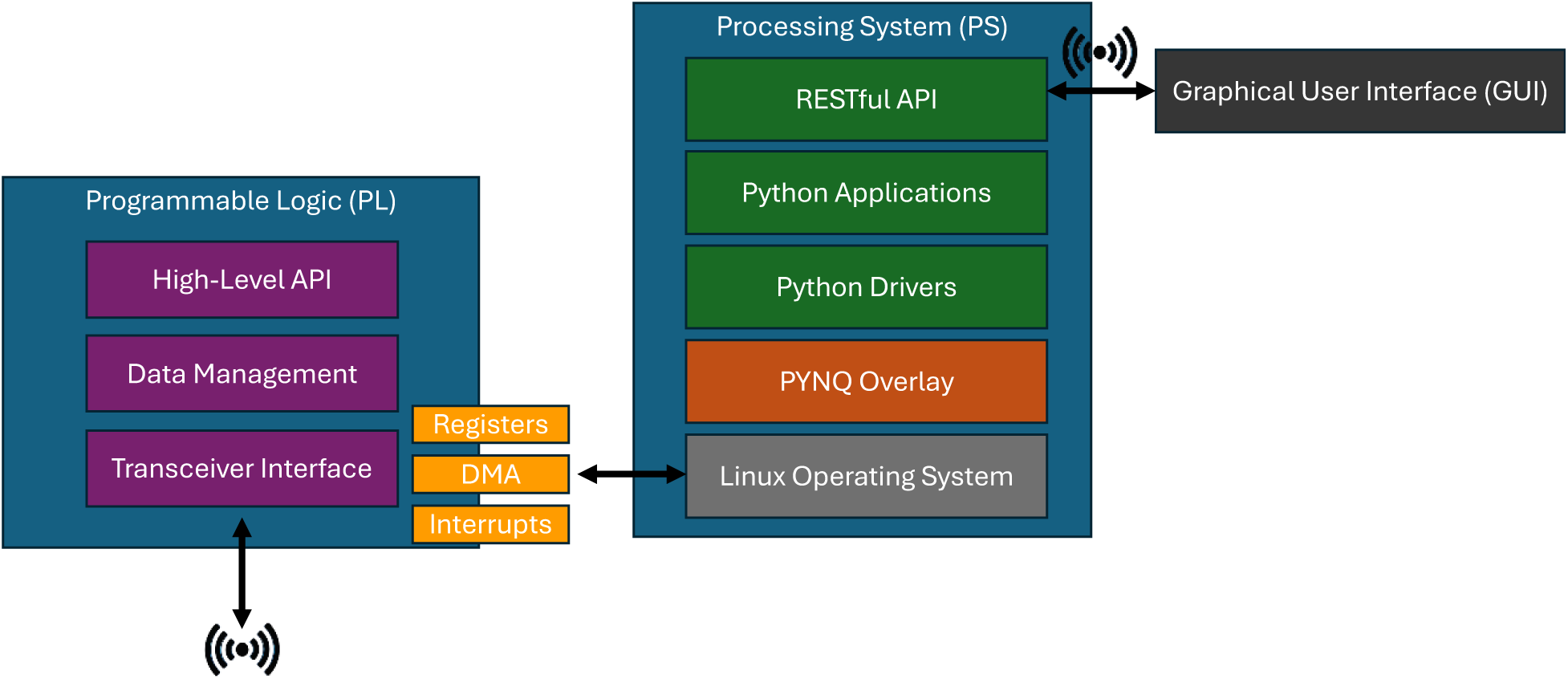
The software stack and hardware interface in the BISC system.

**Fig. S25.**
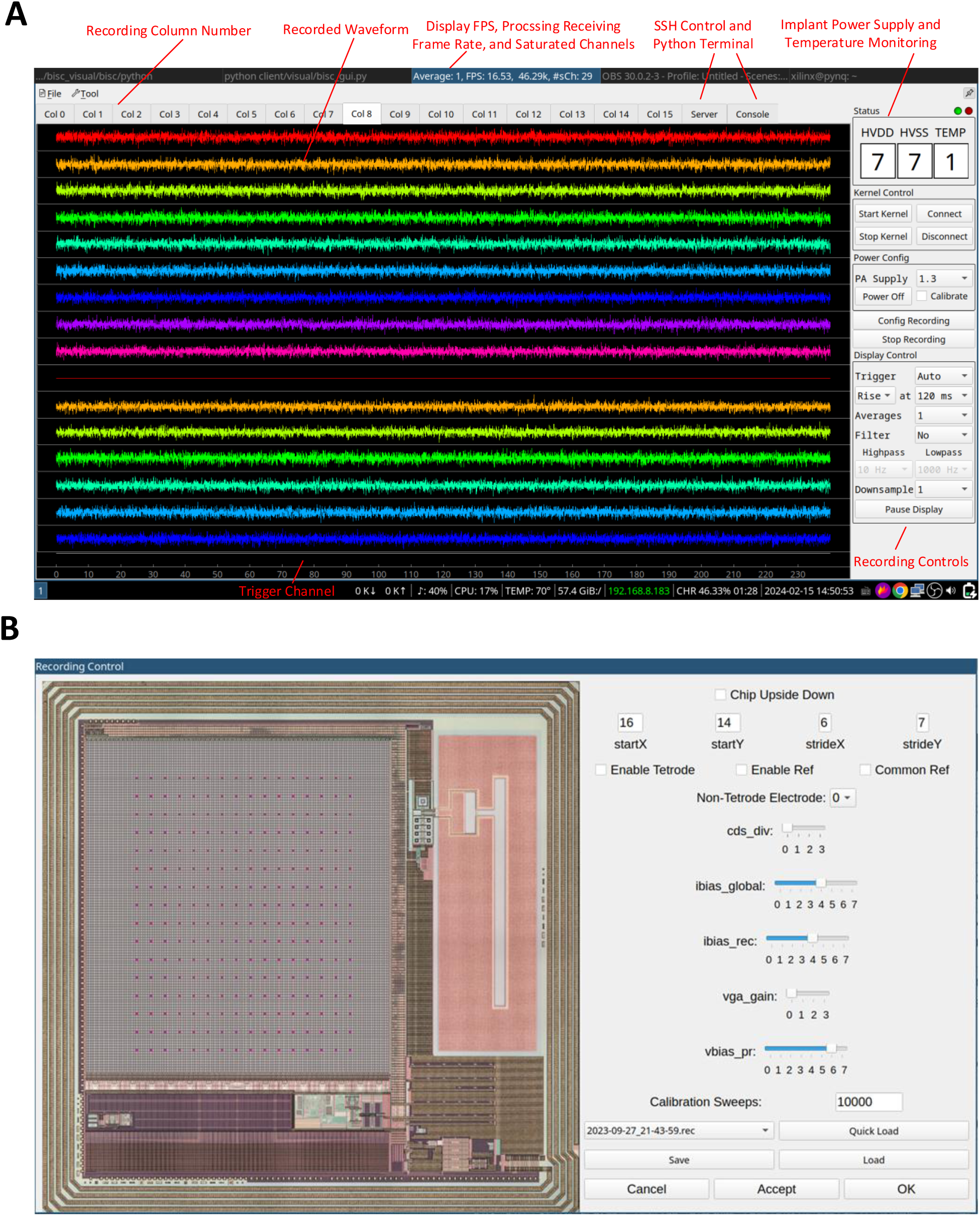
GUI software for recording. (**A**) The main interface displaying 16 channels (rows) in each tab (columns). The user can switch the displayed column on the fly by clicking the recording column number tabs at the top. The recorded data can be played back using the same GUI. The voltage and temperature readings from the implant are displayed on the right-hand side. (**B**) The interface window to configure address of recording pixels (shown as pink dots over the array). Other recording parameters including gain and high-pass corner can also be controlled using sliders.

**Fig. S26.**
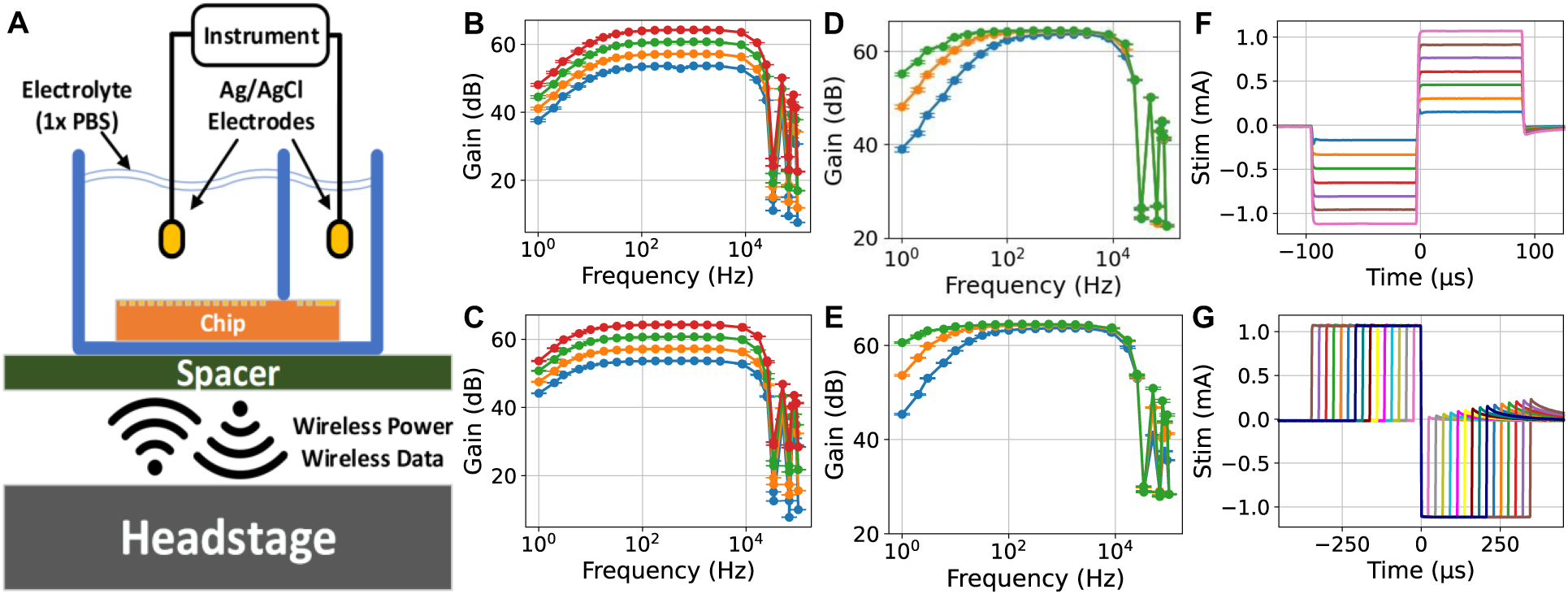
Bench-top *in vitro* characterization. (**A**) Description of the measurement setup. All measurements were taken wirelessly, with 1× phosphate buffered saline (PBS) electrolyte and Ag/AgCl electrodes. Electrodes were either driven by an arbitrary waveform generator (for recording) or connected to a transimpedance amplifier (for stimulation). (**B**) Frequency response across different gain configurations from a representative 16×16 recording. This plot is same as Fig. 2C but reproduced here for comparison with (C) (flat band gains: 53.7 ± 0.20, 57.2 ± 0.21, 60.7 ± 0.20, 64.2 ± 0.19 dB, values: mean ± SD. n = 255, 255, 245, 235). (**C**) Frequency response across different gain configurations from a representative 32×32 recording (flat band gains: 53.7 ± 0.28, 57.2 ± 0.21, 60.7 ± 0.39, 64.2 ± 0.29 dB, values: mean ± SD. n = 1012, 954, 919, 718). (**D**) Frequency response across different high-pass (HP) filter configurations from a representative 16×16 recording. This plot is similar to Fig. 2D but measured with highest PGA gain configuration (3-dB corner: 6.20 ± 2.25, 14.24 ± 2.35, 54.42 ± 1.99 Hz, values: mean ± SD. n = 150, 227, 250). (**E**) Frequency response across different high-pass (HP) filter configurations from a representative 32×32 recording (3-dB corner: 1.41 ± 2.63, 5.16 ± 1.88, 22.10 ± 1.42 Hz, values: mean ± SD. n = 299, 727, 969). (**F**) Representative example of amplitude controlled cathodic-first stimulation (amplitudes: 160, 320, 480, 640, 800, 960, 1120 μA). (**G**) Representative example of pulse width controlled anodic-first stimulation (pulse widths: T, 2T,…,15T where T = 23.04 μs). Error bars in (B) – (E) indicate SE.

**Table S1.** Instruction set architecture of the BISC controller.

| **Instruction Name** | **OPCODE** | **Action** | **Parameters** |
| --- | --- | --- | --- |
| Recording | 001 | Initiates a recording sequence over the configured array dimensions for an optionally specified number of samples | Number of data samples to take from the array |
| Stimulation | 111 | Triggers a pulse stimulation | Width of the stimulation pulse divided to negative, positive and neutral phases |
| Power-on | 110 | Powers-on the pixel amplifiers | N/A |
| Halt | 101 | Stops an on-going recording | N/A |
| Configuration | 010 | Sets the configuration of a target block in the controller | The target block and the unique configuration for the block |
| Programming | 011 | Configures the polarity of the pixels for stimulation | The coordinates of the pixels targeted for stimulation and their initial polarities |
| Query Configuration | 000 | Query for the current state of configuration of a specific block in the controller | Block address |

**Table S2.** Comparison with other wireless brain-computer interface devices.

| Reference | Neuralink<br>39,131 | CorTec<br>132 | Neuropace<br>133 | WIMAGIN<br>E<br>37,134 | WAND<br>33 | **BISC** |
| --- | --- | --- | --- | --- | --- | --- |
| In Vivo Model | Swine,<br>NHP* | Sheep | Human | Human | NHP | Swine, NHP |
| Integration Type | S | SL | SL | S | T | S |
| Dimensions w/o Electrodes (mm) | Radius: 11.5<br>Height: 8* | 60×38×7 | 60×27.5×7.5 | Radius: 25<br>Height: 12.54 | 36×33×15 | 12×12×0.05 |
| Weight (g) | n.r. | n.r. | 16 | n.r. | 17.95 | < 0.1 |
| Recording Electrode Type | Depth Polymer* | ECoG | ECoG | ECoG | Depth Probes | μECoG |
| Total # of Channel | 1024 | 32 | 4 | 64 | 128 | 65536 |
| Simultaneous Recording # of Channel | 1024 | 32 | 4 | 64 | 96 | 256 / 1024 |
| Noise (μV<sub>RMS</sub>) | 6.8 / 8.98<br>(300 – 10k Hz /<br>5 – 1k Hz) | n.r. | n.r. | 1<br>(0.5– 300 Hz) | 1.6 | 7.68 / 16.51<br>(10 – 4k Hz) |
| Amp Gain (V/V) | 411 – 2661 | 100 - 750 | n.r. | up to 1370 | n.r. | 484 – 1620 |
| High-Pass Filter (Hz) | 5 – 300 | 2 | 4 – 12 | 0.5 | 500 | 4 – 55 /<br>1 – 22 |
| Low-Pass Filter (Hz) | 10k | 325 | 30 – 120 | 300 | n.r. | 15.1k |
| Sampling Rate (S/sec) | 20k | 1k | 250 | 1k | 1k | 34k / 8.5k |
| ADC resolution (bits) | 10 | 12 | 10 | 12 | 15 | 10 |
| Power (mW) | n.r. | n.r. | n.a. | 75 | 172 | 64 |
| Power Link Type | IL* | IL<br>250 kHz | IL<br>20 – 50 kHz | IL<br>13.56 MHz | n.a. | IL<br>13.56 MHz |
| Battery | yes* | no | yes | no | yes | no |
| Data Link Type | BLE* | IrDA<br>880 – 900 nm | IL, OOK<br>20 – 50 kHz | FSK<br>402 – 405 MHz | BLE<br>2.4 GHz | UWB-IR<br>3.75 – 4.25 GHz |
| Data Rate (Mbps) | n.r. | n.r. | n.r. | 0.45 | 1.96 | 108/54 |

T: Transcutaneous (Headstage), S: Subcutaneous, SL: Subcutaneous with Lead Connection IL: Inductive Link, PAM: pulse amplitude modulation, UWB-IR: ultra-wideband impulse radio IrDA: Infrared Data Association, BLE: Bluetooth Low Energy, FSK: Frequency Shift Keying \* From press conference <sup>135</sup>

**Video S1.**
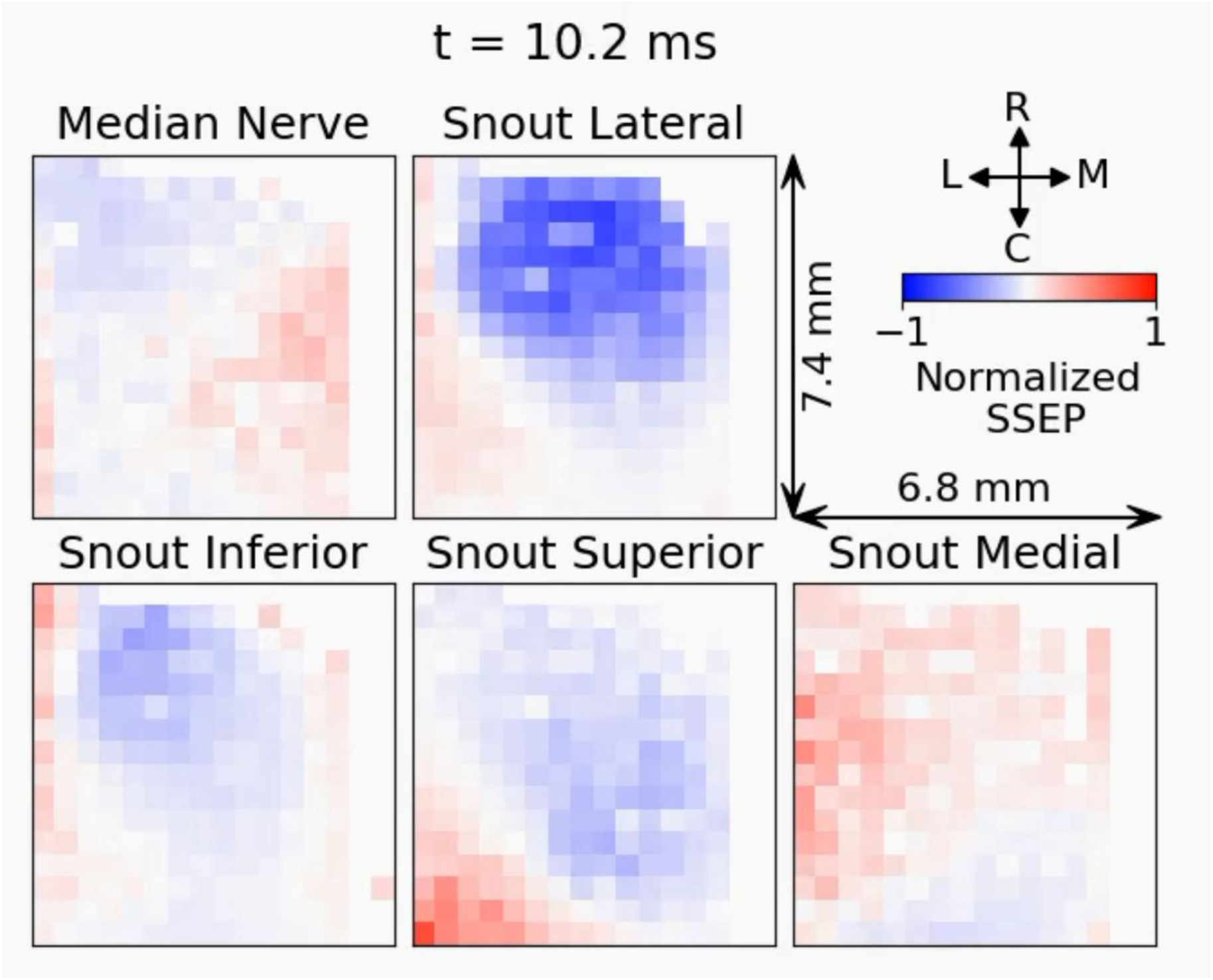
Normalized somatosensory evoked potential (SSEP) recording from a porcine model, trial averaged (n = 100).

**Video S2.**
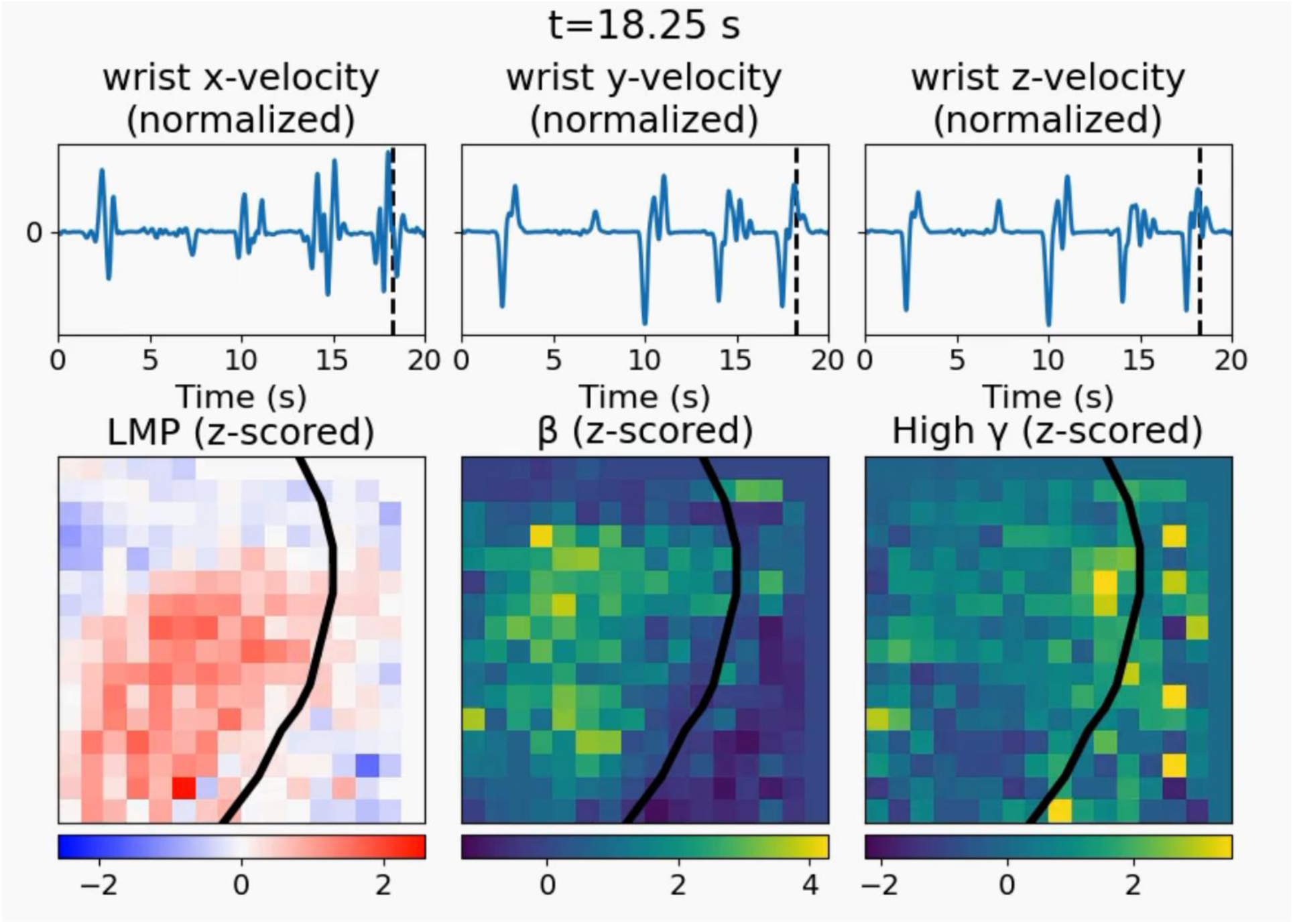
Motor cortex recording from a NHP model performing asynchronous reach-and-grab task.

**Video S3.**
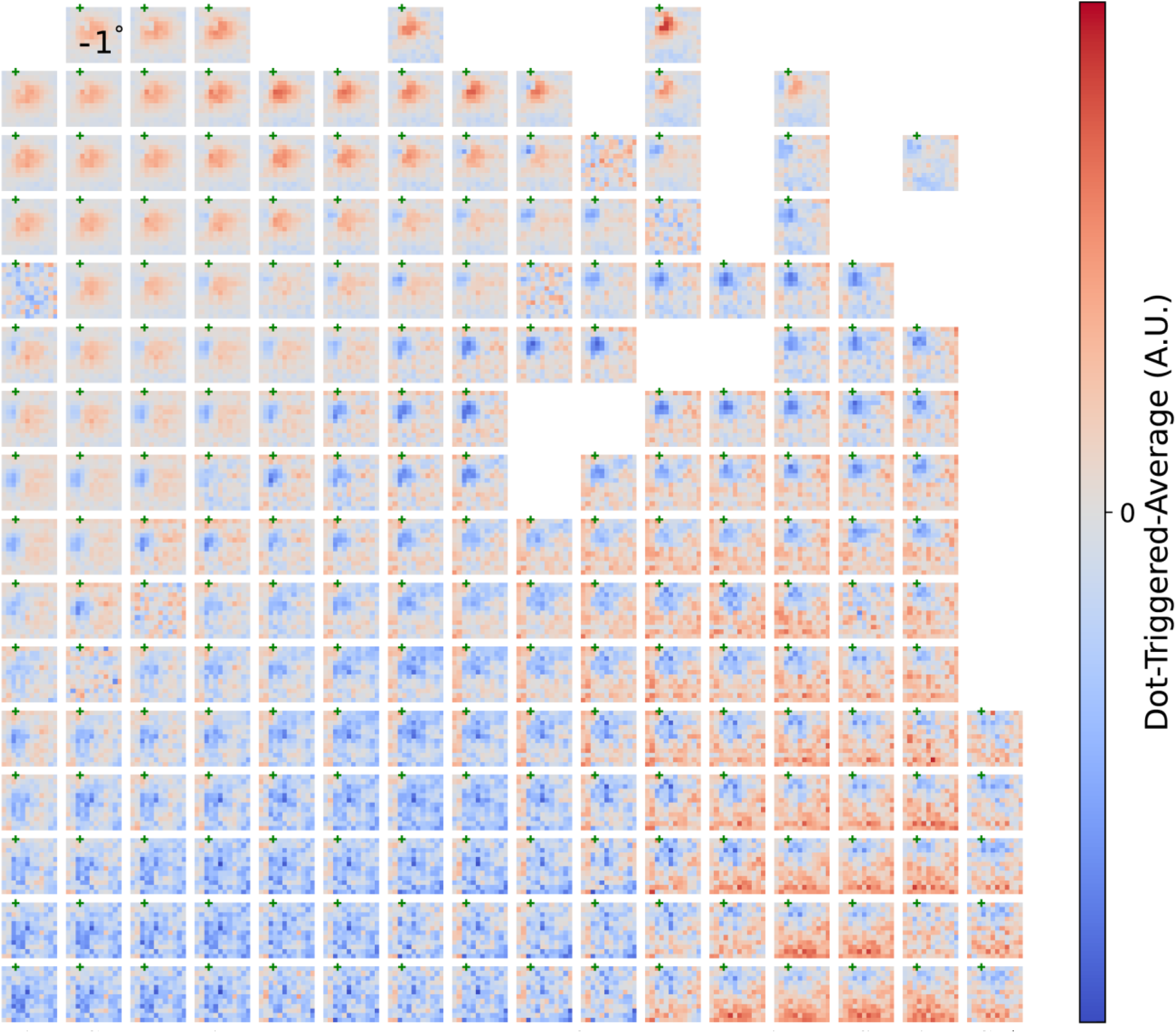
Dot-triggered-average responses of all channels without filtering. Color maps are not shared; however, they are all symmetrical around 0, i.e. same as in **Fig. S12A**. A snapshot at 75 ms is shown.

**Video S4.**
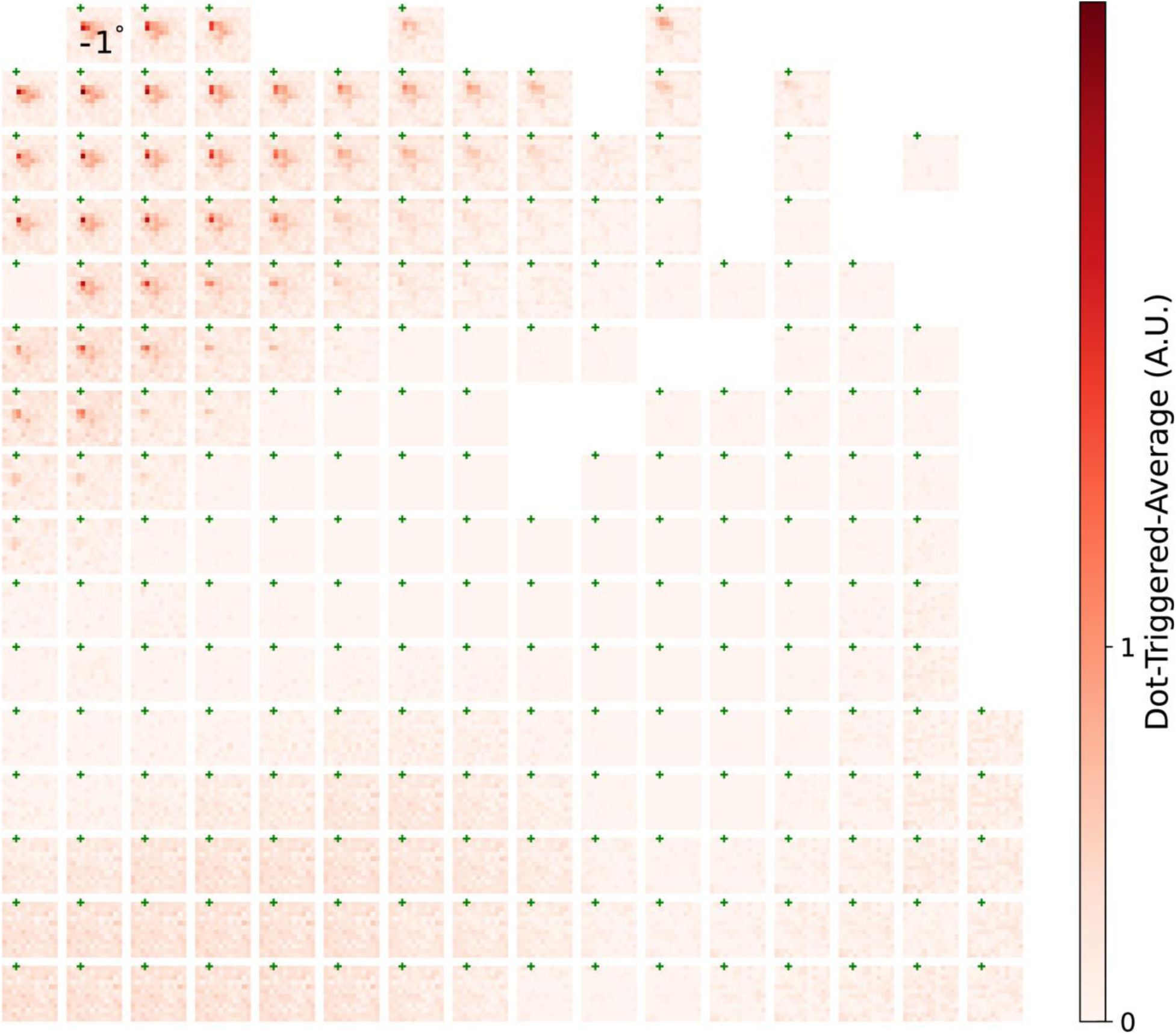
Dot-triggered-average responses of all channels after wavelet transformation (central frequency 8Hz). Color maps are shared across all channels, same as in Fig. 5D. A snapshot at 75 ms is shown.

**Video S5.**
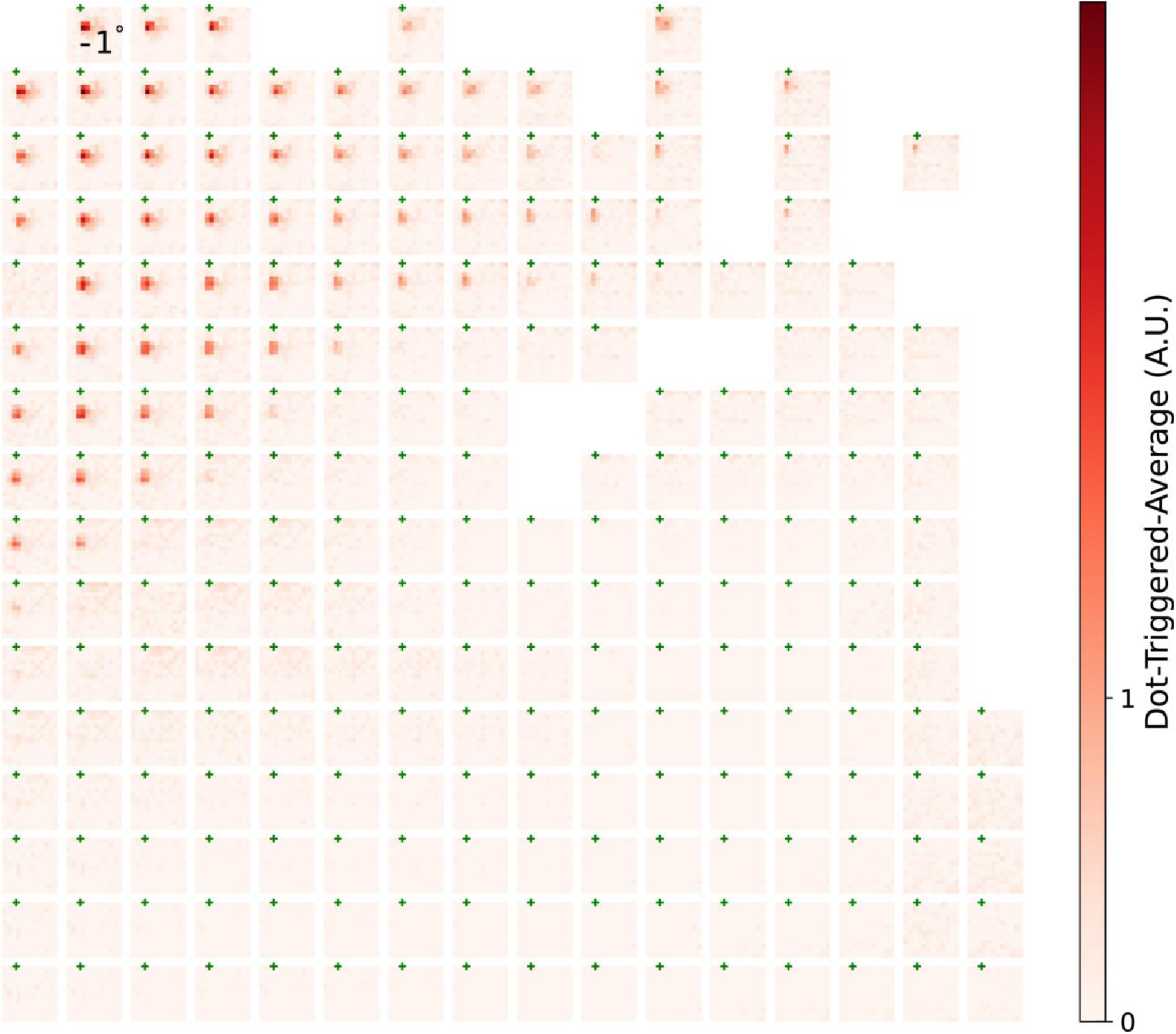
Dot-triggered-average responses of all channels after wavelet transformation (central frequency 16Hz). Color maps are shared across all channels, same as in Fig. 5D. A snapshot at 75 ms is shown.

**Video S6.**
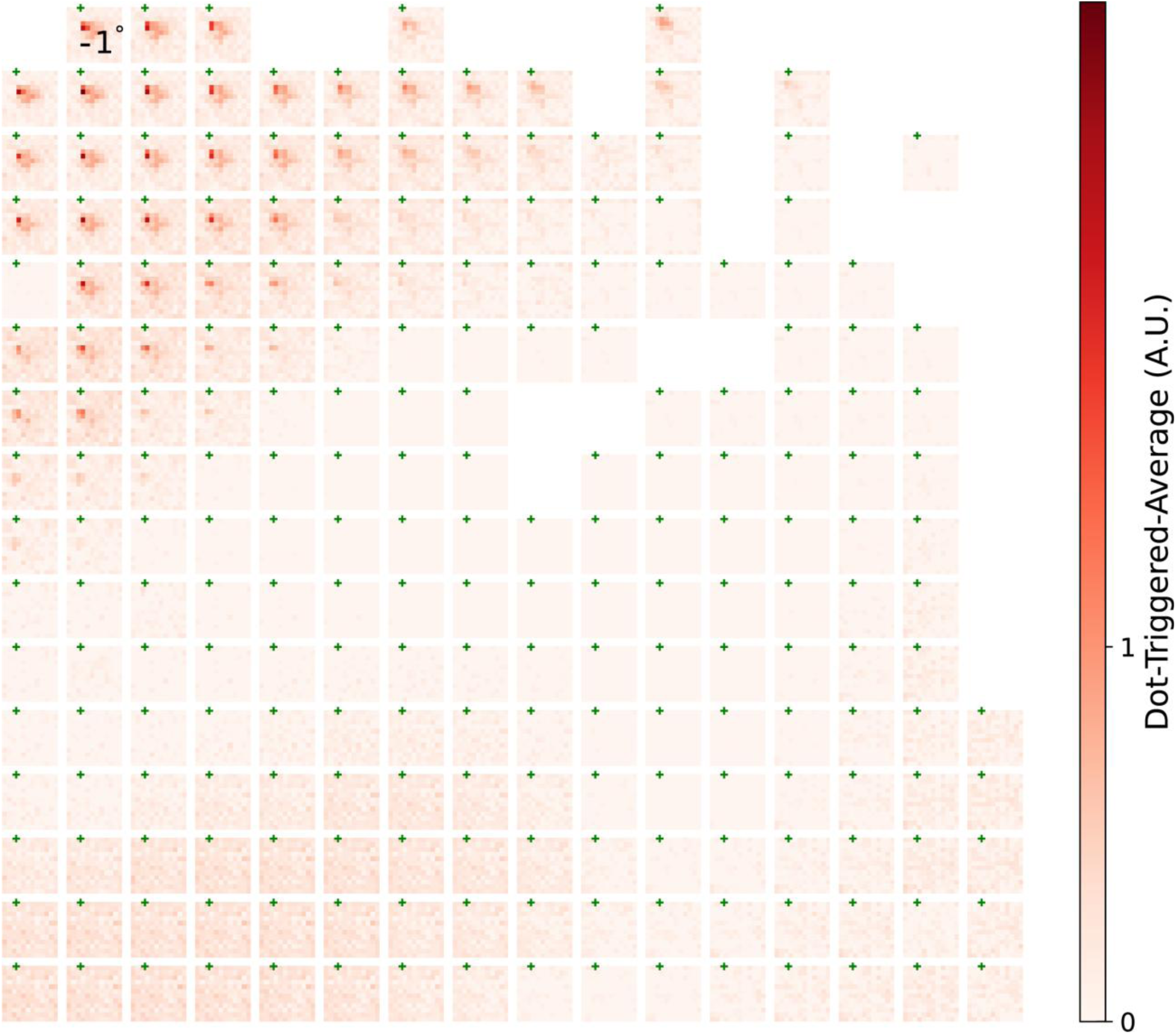
Dot-triggered-average responses of all channels after wavelet transformation (central frequency 32Hz). Color maps are shared across all channels, same as in Fig. 5D. A snapshot at 75 ms is shown.

**Video S7.**
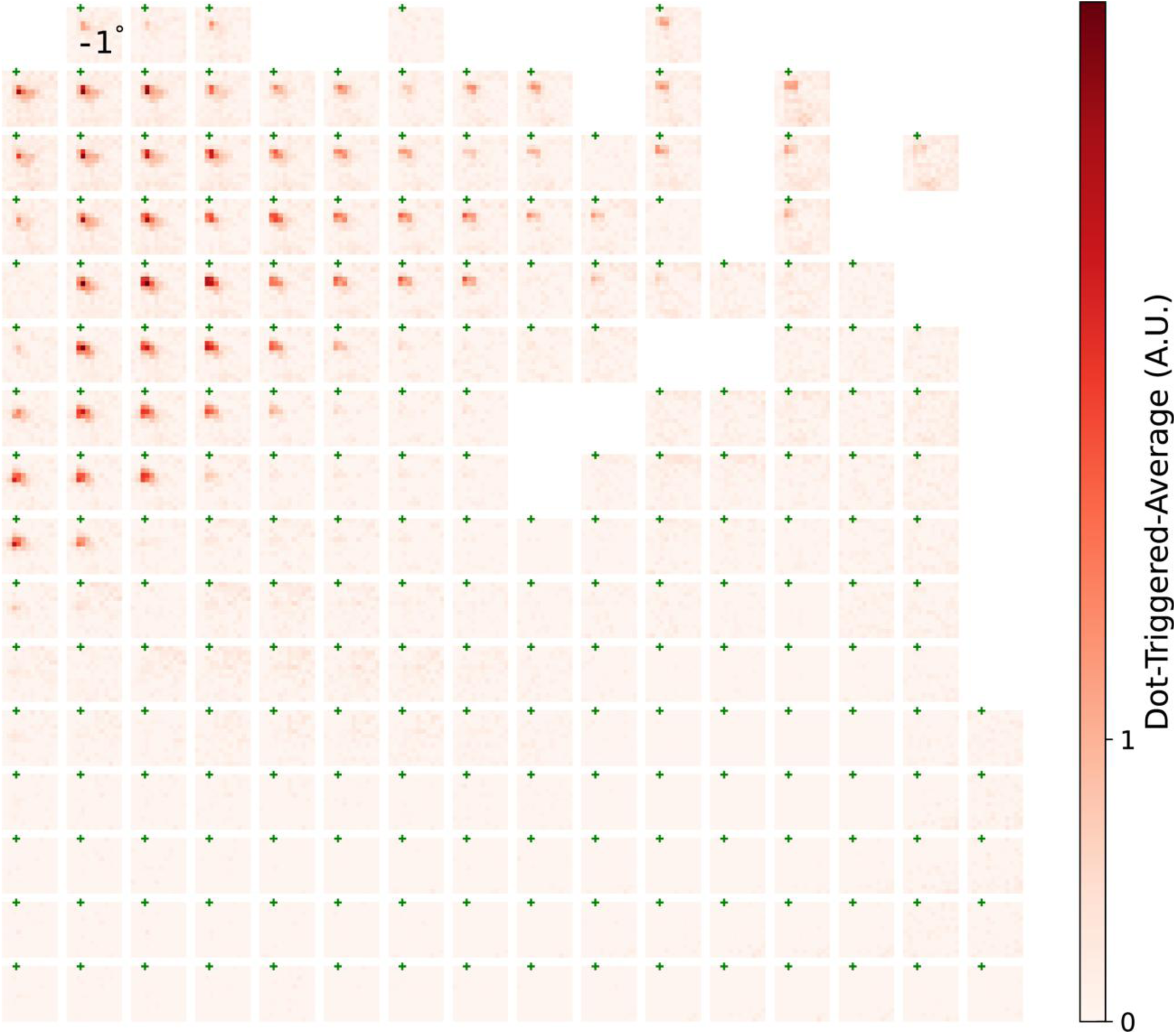
Dot-triggered-average responses of all channels after wavelet transformation (central frequency 64Hz). Color maps are shared across all channels, same as in Fig. 5D. A snapshot at 75 ms is shown.

**Video S8.**
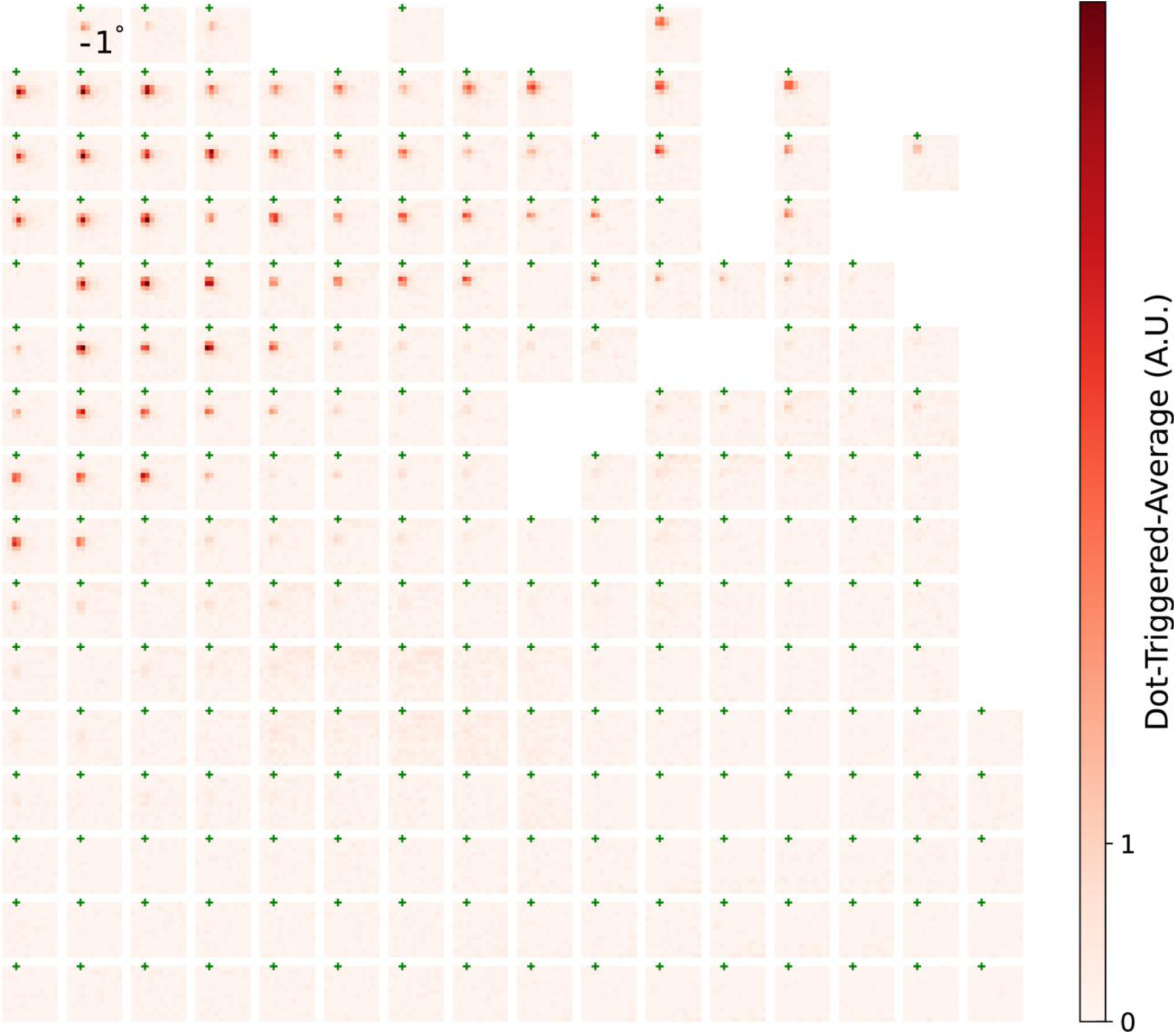
Dot-triggered-average responses of all channels after wavelet transformation (central frequency 128Hz). Color maps are shared across all channels, same as in Fig. 5D. A snapshot at 75 ms is shown.

**Video S9.**
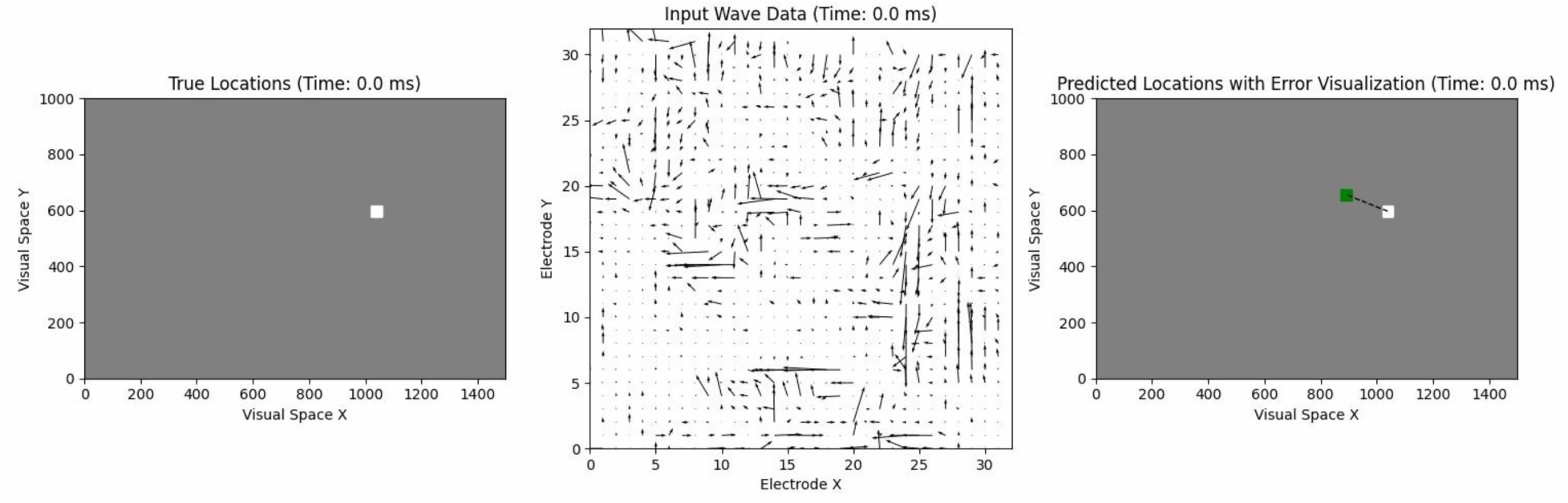
Dot-triggered traveling waves used for decoding stimuli location. The traveling waves are computed from gamma band (30 – 90 Hz) signals recorded from 32×32 spatially dense electrodes at 26.5 μm by 29 μm pitch. Spatiotemporal sequence of these traveling waves, measured within each dot presentation, is used to predict the current location of the dot stimuli presented to the subject.

